# Joint probabilistic modeling of paired transcriptome and proteome measurements in single cells

**DOI:** 10.1101/2020.05.08.083337

**Authors:** Adam Gayoso, Zoë Steier, Romain Lopez, Jeffrey Regier, Kristopher L Nazor, Aaron Streets, Nir Yosef

**Affiliations:** Center for Computational Biology, University of California, Berkeley, Berkeley, CA, USA; Department of Bioengineering, University of California, Berkeley, Berkeley, CA, USA; Department of Electrical Engineering and Computer Sciences, University of California, Berkeley, Berkeley, CA, USA; Department of Statistics, University of Michigan, Ann Arbor, Ann Arbor, MI, USA; BioLegend, Inc., San Diego, CA, USA; Chan Zuckerberg Biohub, San Francisco, CA, USA; Ragon Institute of MGH, MIT and Harvard

## Abstract

The paired measurement of RNA and surface protein abundance in single cells with CITE-seq is a promising approach to connect transcriptional variation with cell phenotypes and functions. However, each data modality exhibits unique technical biases, making it challenging to conduct a joint analysis and combine these two views into a unified representation of cell state. Here we present Total Variational Inference (totalVI), a framework for the joint probabilistic analysis of paired RNA and protein data from single cells. totalVI probabilistically represents the data as a composite of biological and technical factors such as limited sensitivity of the RNA data, background in the protein data, and batch effects. To evaluate totalVI, we performed CITE-seq on immune cells from murine spleen and lymph nodes with biological replicates and with different antibody panels measuring over 100 surface proteins. With this dataset, we demonstrate that totalVI provides a cohesive solution for common analysis tasks like the integration of datasets with matched or unmatched protein panels, dimensionality reduction, clustering, evaluation of correlations between molecules, and differential expression testing. totalVI enables scalable, end-to-end analysis of paired RNA and protein data from single cells and is available as open-source software.

## 1 Introduction

The advance of technologies for quantitative, high-throughput measurement of the molecular composition of single cells is continuously expanding our understanding of cell ontology, state, and function [1–3]. Flow cytometry has set the gold standard for classifying cell types and phenotypic states by quantifying the abundance of small panels of marker proteins on the surface of each cell. More recently, unbiased whole-transcriptome profiling with single-cell RNA sequencing (scRNA-seq) has enabled more comprehensive characterization of cell types [4] and a mechanistic understanding of molecular processes in single cells [5]. Flow cytometry and scRNA-seq offer different views of biological systems at the single-cell level, each with unique insights and limitations. In flow cytometry, for example, fluorescent detection of protein abundance limits how many proteins can be probed simultaneously, whereas scRNA-seq only provides a proxy for the functional information contained in the proteome that does not necessarily reflect protein abundance [6, 7]. The development of techniques such as CITE-seq [8, 9] for paired measurement of RNA and surface proteins from the same cell now provides the opportunity to jointly leverage both types of information. An important feature of CITE-seq is that it quantifies protein abundance by sequencing barcoded antibodies. This significantly increases the number of proteins that can be measured simultaneously relative to flow cytometry and provides the potential to approach proteome-wide profiling of single cells.

Such multi-omic single-cell measurements have the power to provide many new biological insights. First, connecting the extensive literature on cell surface markers with whole-transcriptome profiles could facilitate more consistent and robust annotations of cell types. More fundamentally, having access to a comprehensive molecular profile of each cell could enable a more nuanced, multifaceted understanding of its state [10, 11]. Furthermore, combining these two modalities presents new opportunities such as identifying novel surface markers for new or known cell types, estimating the rates of protein processing [12], and exploring how signals may be received through the cell’s outer membrane and transmitted to produce transcriptional changes. A joint analysis that combines the transcriptomic and proteomic views of a cell could help achieve these goals by using information from both molecular measurements to inform all downstream analysis.

However, combining the information contained in both molecular measurements poses multiple challenges. For example, while the RNA and protein counts observed in CITE-seq data reflect biological factors related to the cell’s underlying state, both measurements contain technical biases that are specific to each modality, making normalization difficult. The RNA data suffer from noise, limited sensitivity, and variation in sequencing depth – factors that have been addressed by a flourishing body of probabilistic methods [13–16]. The protein data, while less sparse than RNA, tend to be obscured by noise and background, which is caused by the detection of ambient or non-specifically bound antibodies. Next, there is the modeling challenge of combining the distinct sets of information and constructing a unified representation for each cell [17, 18]. Finally, there is the challenge of linking a joint data representation to downstream analysis tasks such as dimensionality reduction, clustering, differential expression, and dataset integration. As large-scale community efforts such as the Human Cell Atlas (HCA; [4]) begin to include CITE-seq datasets, these tasks become increasingly challenging because different experiments may use different antibody panels. This is problematic because algorithms designed for these tasks often require direct correspondence of features (genes and proteins) between datasets, which would entail computationally intersecting antibody panels and subsequently, a loss of some or all of the protein information.

These challenges present a need for a scalable, joint statistical model of RNA and protein measurements. Most CITE-seq studies thus far have limited their analysis such that variability between cells was represented by only one modality, while the other modality served to validate and aid with interpretation post-hoc [19–21]. For instance, two recent studies [19, 20] used the RNA data to cluster cells, and then the protein data to derive cluster annotations. Furthermore, statistical methods that have been applied to the protein data do not adequately address the technical factors inherent to this measurement.

Here, we present totalVI (Total Variational Inference), a deep generative model for paired RNA-protein data analysis that addresses these challenges simultaneously. totalVI learns a joint probabilistic representation of RNA and protein measurements that aims to account for the distinct noise and technical biases of each modality as well as batch effects. For RNA, totalVI uses a modeling strategy similar to our previous work on scRNA-seq (scVI; [13]). For proteins, totalVI introduces a new model that separates the protein signal into background and foreground components. It then estimates the probability that the counts of a protein observed in any given cell reflect protein abundance on the cell surface rather than technical artifacts caused by ambient antibodies or non-specific binding. The probabilistic representations learned by totalVI are built on a joint low-dimensional view of the RNA and protein data that is derived using neural networks. This joint view captures the biological variability between cells and can be interpreted as representing the underlying state of each cell. As such, this single low-dimensional representation of cell state reflects the coordinated regulation of transcription and translation that produces both molecules in the same cell.

The totalVI joint probabilistic model provides a cohesive solution for common analysis tasks such as dimensionality reduction, clustering, evaluation of correlations between genes and/or proteins, and differential expression testing. Prior to conducting these procedures, totalVI can be used for dataset integration, even in the case of partially- or non-overlapping protein panels, which result in “missing” protein entries. In addition to the integration of two or more input datasets, totalVI can impute the expression of proteins that are missing in certain samples, including ones with no measured proteins such as those from scRNA-seq experiments. Finally, while totalVI is based on deep neural networks, it provides a way to relate the coordinates of its latent dimensions to the expression of specific genes and proteins, thus affording interpretability and highlighting interesting trends in the data. In each of these tasks, totalVI accounts for batch effects, protein background, and other nuisance factors to enable more accurate analysis.

To evaluate the performance of totalVI, we conducted a series of CITE-seq experiments on murine spleen and lymph nodes that were designed to include complications like biological replicates, different protein panels, and up to 208 measured proteins per experiment. Using this data along with several public datasets, we demonstrated that totalVI effectively decouples protein foreground and background, integrates datasets with either matched or unmatched protein panels, imputes the measurements of proteins that were not included in one of the panels, and detects differential expression of both RNA and proteins. We demonstrated that totalVI compares favorably to the state of the art methods in each of these tasks (in cases where such a method is available), and that it scales to large dataset sizes. We applied totalVI to characterize the heterogeneity of B cells in the spleen and lymph nodes with paired RNA and protein information. totalVI and its downstream analysis procedures are available as open source software, which can be readily used as an end-to-end analytical solution for the increasingly common CITE-seq technology.

## 2 Results

### 2.1 The totalVI model

totalVI uses a probabilistic latent variable model [22] to represent the uncertainty in the observed RNA and protein counts from a CITE-seq experiment as a composite of biological and technical sources of variation. The input to totalVI consists of the matrices of RNA and protein unique molecular identifier (UMI) counts (Figure 1a). Categorical covariates such as experimental batch or patient are optional inputs used for integrating datasets, and referred to henceforth as “batch”. totalVI can integrate datasets with different antibody panels (including integration with standard scRNA-seq), and use its probabilistic model to impute any missing protein measurements.

**Figure 1:**
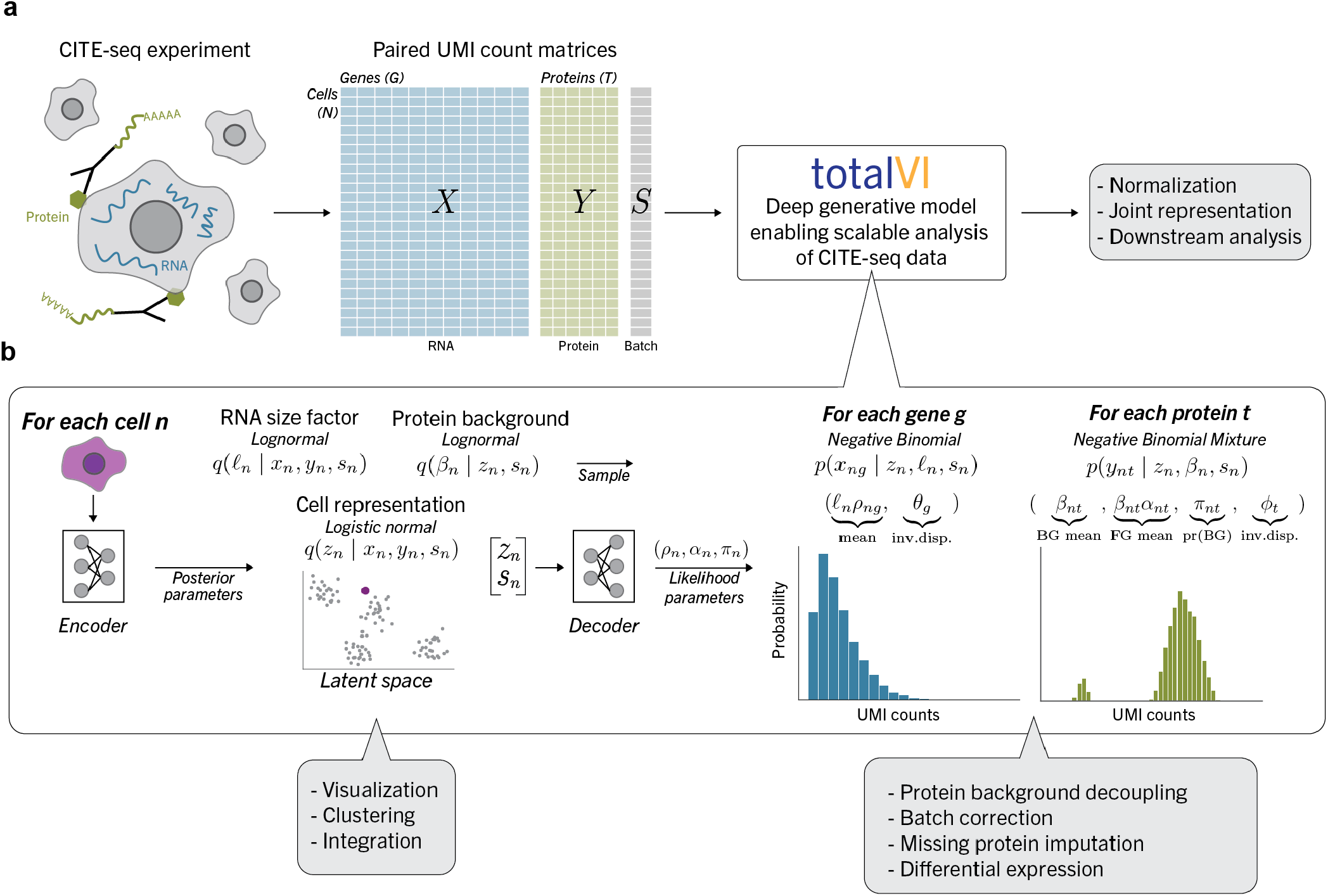
Schematic of a CITE-seq data analysis pipeline with totalVI. **a**, A CITE-seq experiment simultaneously measures RNA and surface proteins in single cells, producing paired count matrices for RNA and proteins. These matrices, along with an optional matrix containing sample-level categorical covariates (batch), are the input to totalVI, which concomitantly normalizes the data and learns a joint representation of the data that is suitable for downstream analysis tasks. **b**, Schematic of totalVI model. The RNA counts, protein counts, and batch for each cell *n* are jointly transformed by an encoder neural network into the parameters of the posterior distributions for *z_n_*, a low-dimensional representation of cell state, *β_n_*, the protein background mean indexed by protein, and *ℓ_n_*, an RNA size factor. The space of all joint cell representations (latent space) is independent of the batch and is used for dataset integration and cell stratification tasks. Next, a decoder neural network maps samples from the posterior distribution of *z_n_*, along with the batch, *s_n_*, to parameters of a negative binomial distribution for each gene and a negative binomial mixture for each protein, which contains a foreground (FG) and background (BG) component (Methods 4). These parameters are used for downstream analysis tasks.

The output of totalVI consists of two components that can be used for downstream analysis. The first component encodes each cell as a distribution in an unobserved (latent) low-dimensional space that represents the information contained in the RNA and protein data and controls for their respective noise properties. The second component provides a way to estimate the parameters of the distributions that underlie the observed RNA and protein measurements (i.e., likelihoods) given a cell’s latent representation. These distributions explicitly account for nuisance factors in the observed data such as sequencing depth, protein background, and batch effects (Figure 1b).

To model the RNA data, totalVI posits that for each gene *g* and cell *n* the number of observed transcripts, *x_ng_*, follows a negative binomial distribution, which has been shown to adequately handle the limited sensitivity and over-dispersion that are characteristic of this data [23]. The mean of the negative binomial, *ℓ_n_ρ_ng_*, depends on the cell’s batch, denoted *sn*, and two latent variables. The first variable, *ℓ_n_*, represents cell size and sequencing depth and follows a lognormal distribution. The second variable, *z_n_*, is a low-dimensional vector (here, 20 dimensions), which follows a logistic normal distribution and can be interpreted as the underlying biological state of the cell. A neural network maps *z_n_* and *s_n_* to *ρ_ng_*, which represents the normalized gene expression. This modeling strategy for the RNA data is closely related to previous work in this domain (e.g., scVI [13], ZINB-WAVE [16], and DCA [24]).

For the protein data, totalVI posits that the number of molecules for each protein *t* and cell *n*, *y_nt_* follows a two-component negative binomial mixture distribution. The mixture accounts for the fact that the observed protein counts may correspond to either real protein abundance on the cell surface, or to technical artifacts like ambient antibodies and non-specific binding. The mean of this mixture depends on *z_n_* (similarly to RNA) and on another latent variable, *β_nt_*, which follows a lognormal distribution and describes a baseline level of protein counts due to background. An additional neural network maps *z_n_* and *s_n_* to the protein likelihood parameters *α_n_* and *π_n_*, which represent an offset for the protein foreground mean, and the probability that the observed counts were generated from the background component, respectively.

totalVI optimizes the parameters of both of its components simultaneously using the variational autoencoder (VAE) framework [25]. In the first component, “encoder” neural networks are trained to map the raw observations to the parameters of the approximate posterior distributions of the latent variables *z_n_*, *ℓ_n_*, and *β_n_*. In the second component, the previously described “decoder” neural networks are trained to map *z_n_* and *s_n_* to the RNA and protein likelihood parameters *ρ_n_*, *α_n_*, and *π_n_*. The inverse dispersion parameters for genes *θ_g_* and proteins *ϕ_t_* are model parameters learned during inference. Accordingly, totalVI uses highly efficient techniques for stochastic optimization of VAEs, which makes it appropriate for the scale of CITE-seq data.

The latent variable *z_n_* in the first component of totalVI is particularly important for downstream analysis. As *z_n_* is linked to the mean of the likelihood for each protein and each gene in each cell by a neural network, its approximate posterior, *q*(*z_n_* | *x_n_*, *y_n_*, *s_n_*), captures information contained in both RNA and protein measurements. This distribution should also be robust to the nuisance factors accounted for, such as negative binomial noise and batch effects. We used the mean of *z_n_* under the approximate posterior as an integrated, joint view of cell state, which we then used as input to clustering and visualization algorithms. Additionally, we modeled *z_n_* with a logistic normal distribution, meaning each *z_n_* resides in a convex polytope amenable to archetypal analysis [26, 27]. Such an analysis connects each latent dimension to the expression of genes and proteins and aids with the interpretation of the model.

Other downstream tasks specific to genes and proteins (collectively referred to as features), are linked to the likelihood parameters from the second component of totalVI. In the following, we demonstrate how to use this part of the model in a range of tasks such as differential expression testing, missing protein imputation, and measurement denoising. A detailed specification of the model, along with further mathematical descriptions of the quantities used in downstream tasks is in Methods 4.

### 2.2 CITE-seq profiling of murine spleen and lymph nodes

We conducted a series of CITE-seq experiments that were designed to test the performance of totalVI on a variety of tasks while handling issues related to large protein panels that can vary between different batches. As a case study, we profiled murine spleen and lymph nodes, which contain heterogeneous immune cell populations that are well-characterized by surface protein markers. The CITE-seq experiments were executed on the 10x Genomics Chromium microfluidic platform using BioLegend TotalSeq-A antibodies.

In these experiments, cells were collected from two wild-type mice that were processed in two different experimental runs to serve as biological replicates (Methods 4.4). For each experimental run (completed on separate days), we stained cells from one mouse with two different panels of barcoded antibodies: the first contained 111 antibodies and the second contained 208 antibodies, of which the 111 antibodies were a subset (Supplementary Data: Antibodies). Spleen and lymph node cells were stained separately with the same antibody panels plus an additional hashtag antibody such that the tissues could be demultiplexed [28]. Cells stained with the same panel were pooled in the same 10x lane, resulting in four lanes (two per run).

We refer to these four spleen/lymph node datasets by their protein panel and experimental day: SLN111-D1, SLN208-D1, SLN111-D2, SLN208-D2. We refer to the combination of the four as SLN-all (experimental design in Table 1). After pre-processing and filtering, these datasets contained a total of 32,648 cells with approximately 3,300 UMIs per cell for RNA and 2,800 UMIs per cell for proteins (Methods 4.6).

**Table 1:** Summary of spleen and lymph node datasets. Each dataset was processed in a separate 10x lane. Each day indicates a 10x run. Cells captured is the number of cells reported by Cell Ranger.

| Dataset name | Antibody panel | Day | Mouse | Tissue | Cells (captured) | Cells (post-filtering) |
| --- | --- | --- | --- | --- | --- | --- |
| SLN111-D1 | 111 | 1 | A | Spleen; Lymph Node | 11,160 | 9,264 |
| SLN111-D2 | 111 | 2 | B | Spleen; Lymph Node | 9,017 | 7,564 |
| SLN208-D1 | 208 | 1 | A | Spleen; Lymph Node | 10,777 | 8,715 |
| SLN208-D2 | 208 | 2 | B | Spleen; Lymph Node | 8,921 | 7,105 |

**Table 2:** Sequencing statistics for spleen and lymph node datasets. Sequencing statistics calculated per 10x lane by Cell Ranger. RNA reads: mean reads per cell from RNA. Protein reads: mean reads per cell from antibody barcodes. RNA UMI: median UMI counts per cell from RNA. Protein UMI: median UMI counts per cell from antibody barcodes.

| Name | RNA reads | Protein reads | RNA UMI | Protein UMI |
| --- | --- | --- | --- | --- |
| SLN111-D1 | 34,717 | 4,733 | 4,392 | 2,785 |
| SLN111-D2 | 45,765 | 6,542 | 2,121 | 3,419 |
| SLN208-D1 | 33,569 | 5,513 | 4,561 | 2,956 |
| SLN208-D2 | 43,821 | 3,961 | 2,102 | 2,261 |

**Table 3:** Summary of filtering parameters for publicly available datasets. Ranges indicate criteria for retained cells.

| Dataset | No. cells | Pct. Mito | Protein Lib Size Range | No. Genes Expr. | RNA lib size |
| --- | --- | --- | --- | --- | --- |
| PBMC10k | 6,855 | < 10% | [400, 20,000] | < 4,500 | < 20,000 |
| PBMC5k | 3,994 | < 20% | [400, 20,000] | < 4,500 | < 20,000 |
| MALT | 6,838 | < 15% | [400, 20,000] | < 5,000 | < 30,000 |

**Table 4:**
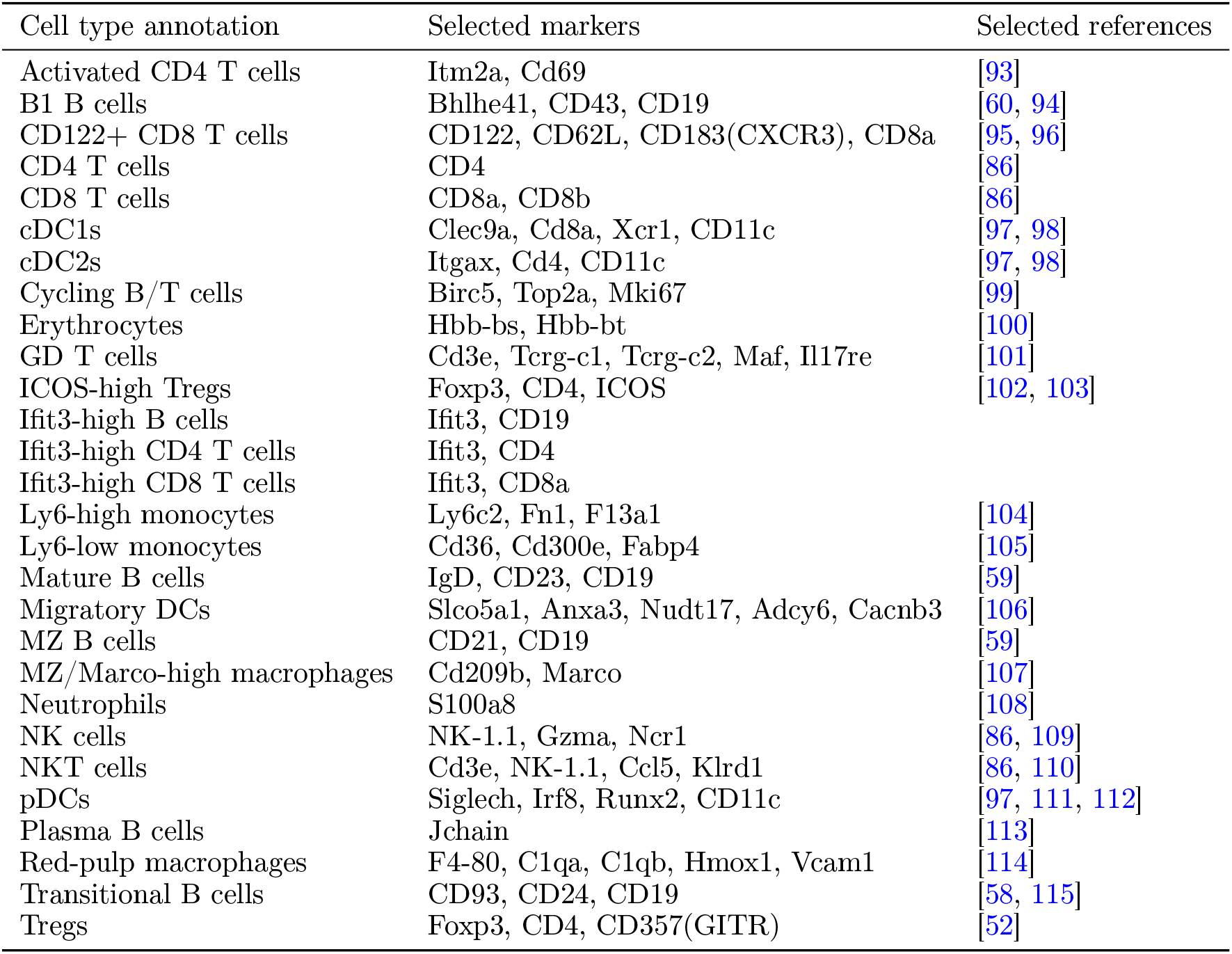
Annotated cell types and selected markers in the spleen and lymph node datasets. cDC1: Conventional dendritic cell 1. cDC2: Conventional dendritic cell 2. GD: Gamma/delta. MZ: Marginal zone. NK: Natural killer. NKT: Natural killer T. pDC: Plasmacytoid dendritic cell. Treg: Regulatory CD4 T cell.

### 2.3 totalVI is scalable and fits CITE-seq data well

The usefulness of probabilistic models like totalVI depends on how well they fit the observed data. Furthermore, they should generalize to unobserved data (i.e., not overfit) and scale to a realistic range of input sizes. Here we verify that totalVI satisfies these prerequisites.

We first estimated how well totalVI fits CITE-seq data that was available to it during training using posterior predictive checks (PPC; [14, 29]). To conduct PPCs, we generated replicated datasets (i.e., posterior predictive samples) by sampling from the fitted model (Methods 4.7). One replicated dataset is constructed by passing each observed cell *n* through the encoder part of the model, sampling from the posterior approximation of its latent variables (e.g., *z_n_*), and then sampling from the conditional distributions for each gene and protein given the latent variables (Figure 1b). As totalVI directly models the count data, the replicated datasets generated with this procedure should maintain the properties of the observed data. To that end, we assessed how well totalVI preserves the mean-variance relationship of the data by measuring the similarity between the coefficient of variation (CV) per gene and protein on the replicated and the raw data.

We then evaluated how well totalVI generalizes to data that was not available during training (i.e., a set of held-out cells). To do so, we generated replicated datasets conditioned on the held-out cells, and computed two somewhat opposing metrics of predictive performance. First, we assessed how well the average replicated data set matched the observed held-out data via mean absolute error. Second, we quantified how well the interval of values from replicated data sets covered the observed held-out data values (calibration error; [30]). We also computed the held-out predictive log-likelihood of the data. All of these held-out metrics were computed separately for genes and proteins.

As a baseline, we compared totalVI to factor analysis (FA), which can be viewed as a simplified version of totalVI. For instance, in FA, the mapping between the latent space and data space is specified by a linear function and the conditional distribution for each feature is Gaussian. As an additional control, we compared to scVI [13], which was restricted to the RNA part of the data. We expected the goodness of fit for totalVI and scVI on just the RNA part of the data to be comparable, as they share a similar generative model and neural network architecture.

Our evaluation relied on fitting the models to several CITE-seq datasets, spanning different species and tissues, including peripheral blood mononuclear cells (PBMC10k; [31]) and mucosa-associated lymphoid tissue (MALT; [32]) from humans, and our novel murine spleen and lymph node data (SLN111-D1). Across all datasets, totalVI performed best in terms of preserving the mean-variance relationship of genes and proteins in the raw data (Supplementary Figure S1a). On the held-out protein data, totalVI outperformed FA in both the mean absolute error and calibration error metrics. On the held-out RNA data, totalVI and scVI were largely comparable and outperformed FA (Supplementary Figure S1b-d).

To assess the scalability of totalVI, we concatenated all of our spleen and lymph node data (SLN-all) and recorded the training time for different sizes of subsets of this data. totalVI and scVI (which processes only RNA data) had similar dependence between run time and input size (Supplementary Figure S1e). Furthermore, we observed that totalVI can readily handle large data sets, for instance, processing the complete set of approximately 33,000 cells with over 4,100 features in less than an hour.

Overall, these results demonstrate that totalVI is a scalable probabilistic model that provides an accurate representation of CITE-seq data. With these prerequisites satisfied, we turned to benchmarking particular aspects of the model as they relate to common single-cell analysis tasks.

### 2.4 totalVI identifies and corrects for protein background

To analyze protein data in an accurate and quantitative manner, it is necessary to distinguish between true biological signal and technical bias in the protein measurement. Background is a type of technical bias that has previously been reported for antibody-based measurements and has been observed in both flow cytometry [33] and sequencing-based [8, 9] studies. In CITE-seq data, protein counts contain non-negligible background, the extent of which can be observed in the detection of counts for nearly every protein in all cells. This background arises experimentally from a combination of unbound ambient antibodies, which can be detected in empty droplets, and non-specific antibody binding, which can be detected in cells with no expected expression of a protein, such as CD19 in T cells (Methods 4.8, Supplementary Figure S2a-c, g). Recent methods have described background from ambient RNA [34, 35], but the presence of background is more pronounced in protein measurements (Supplementary Figure S2d-f). Failure to account for this protein background could lead to erroneous conclusions in downstream analyses. For instance, differential expression tests might report false positives driven by variation in background rather than by the biological signal of interest, as well as false negatives if high levels of background obscure low levels of signal.

Accurate identification of protein foreground and background is a prerequisite for denoising the protein measurements. Previous studies of CITE-seq data derived a single decision rule for every protein, specifying the minimal number of counts required to be considered foreground. In one study, the boundary was based on spiked-in negative control cells from another species, relying on the assumption that cross-reactivity of antibodies between species is minimal [8]. Another study fit a Gaussian mixture model (GMM) to distinguish a background and foreground component for each protein [36]. Using the same boundary for all cells, however, relies on the assumptions that all cells are subject to a similar background distribution of the protein in question and, for the commonly applied case of a two-component GMM, that the foreground component is comparable across cell types.

Since these assumptions might not hold in all cases, totalVI instead models protein background as cell- and protein-specific. In totalVI, the estimate of protein background accounts not only for the observed levels of a given protein in the cell in question, but also for the overall transcriptomic and proteomic profile of that cell. To do this, totalVI models each protein measurement as a mixture of a foreground and background component that depends on the cell’s representation in the latent space (Methods 4.1). The mixture is weighted by the parameter *π*, which can be interpreted as the probability that the counts of a protein in a given cell came from the background component (Figure 1b, Methods 4.8).

As a way of evaluating totalVI’s ability to quantitatively identify protein background using *π*, we made use of common marker proteins, whose counts are expected to come from the foreground component in some cell types and from the background component in others. For example, a high foreground probability for CD4 could be used as a positive predictor of CD4 T cells. We tested how well the foreground probability estimates (1 − *π*) of each marker protein in the SLN111-D1 dataset performed at classifying major cell types by computing the area under the receiver operating characteristic curve (ROC AUC), using manual cell type annotations as the true labels. As a baseline for comparison, we used the assignment probabilities from a two-component Gaussian mixture model (GMM) fit on all cells for each log-transformed protein (Methods 4.8).

For nine out of eleven known marker proteins, both totalVI and the GMM performed well at classifying cell types by protein foreground probability (ROC AUC > 0.97; Supplementary Figure S3a). For these marker proteins, distinguishing foreground and background components was relatively straightforward, and foreground and background levels appeared to be similar across all cells. The B cell marker CD19 was one example where the distributions of foreground and background protein expression were easily separated (Supplementary Figures S2a and S4a-d).

Other proteins, however, were not as simple, and had highly overlapping distributions of foreground and background counts (Supplementary Figure S2b, c). For two of these more challenging proteins, namely, the B cell marker CD20 and the T cell marker CD28, totalVI noticeably outperformed the GMM (Supplementary Figure S3b). These two proteins are illustrative of two types of errors that can be made when using a single decision boundary on all cells to distinguish protein foreground from background.

In the case of CD20 (encoded by Ms4a1 RNA), the GMM set a high cutoff on protein counts that resulted in numerous false negatives (Methods 4.8). Among the cells below the cutoff (considered CD20 negative by the GMM), totalVI distinguished cells with high and low foreground probability (Figure 2a). The cells with high foreground probability clustered with B cells in the totalVI latent space and had high Ms4a1 expression, confirming their identity as B cells. On the other hand, cells with similarly low protein expression but with low probability foreground clustered with T cells and had no Ms4a1 expression (Figure 2a-c).

**Figure 2:**
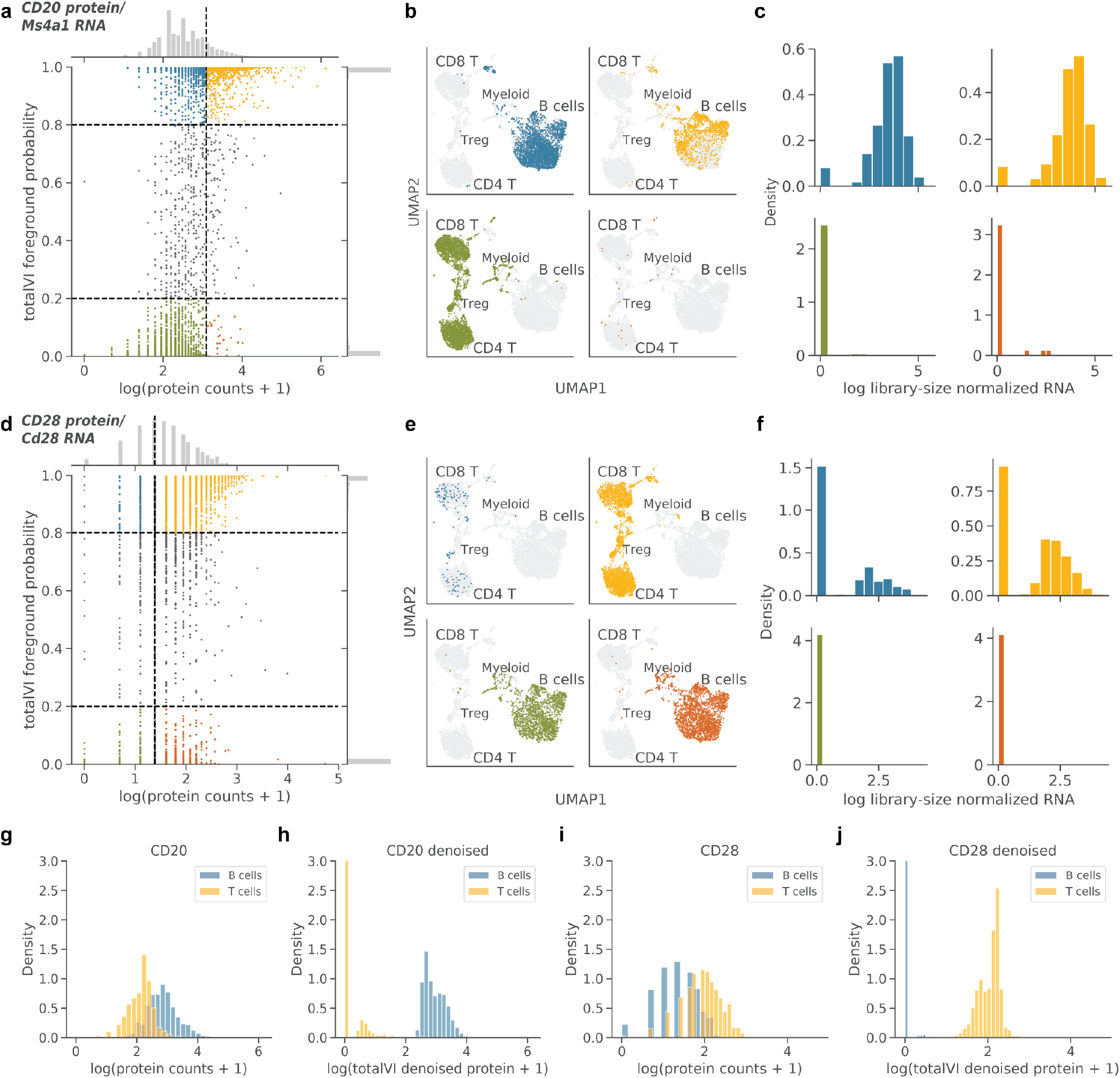
totalVI identifies and corrects for protein background. totalVI was applied to the SLN111-D1 dataset. **a-c**, CD20 protein (encoded by Ms4a1 RNA). **(a)** totalVI foreground probability vs log(protein counts + 1). Vertical line denotes protein foreground/background cutoff determined by a GMM. Horizontal lines denote totalVI foreground probability of 0.2 and 0.8. Cells with foreground probability greater than 0.8 or less than 0.2 are colored by quadrant, while the remaining cells are gray. **(b)** UMAP plots of the totalVI latent space. Each quadrant contains cells from the corresponding quadrant of (a) in color with the remaining cells in gray. **(c)** RNA expression (log library-size normalized; Methods 4.8) for cells colored in (a). **d-f**, Same as (a-c), but for CD28 protein (encoded by Cd28 RNA). **g, h**, Distributions of log(protein counts + 1) **(g)** and log(totalVI denoised protein + 1) **(h)** for CD20 protein in B and T cells. y-axis is truncated at 3. **i, j**, Same as (g, h), but for CD28 protein.

In the case of CD28, the GMM set a low cutoff on protein counts that resulted in numerous false positives. Among the cells above the GMM cutoff, totalVI distinguished the cells with high foreground probability (primarily T cells) from the cells with low foreground probability (primarily B cells) (Figure 2d-f). Across all proteins, the totalVI foreground probability tended to fall near zero or one (Supplementary Figure S4e).

Despite using a two-component mixture, totalVI can also decouple background and foreground for proteins that have greater than two modes of expression globally. For example, totalVI correctly decoupled background and foreground for CD4 in peripheral blood mononuclear cells, which globally follows a trimodal distribution (Methods 4.8, Supplementary Figure S5a, b). Through the above examples, we see that by leveraging information from all cells, genes, and proteins, totalVI is capable of identifying protein foreground and background in a way that more accurately reflects cell state than a single decision boundary on protein expression.

To illustrate background identification, we demonstrated that decoupling foreground from background can be used to “classify” a cell as expressing or not expressing a certain protein. For downstream analysis, totalVI uses *π* in a more quantitative manner to remove protein background. Specifically, totalVI can denoise the protein data by setting the background component to zero, while also accounting for the measurement uncertainty of the foreground component. This effectively subtracts the expected amount of background from the overall expected expression (Methods 4.3 and 4.8, Figure 2g-j, Supplementary Figure S5f, g).

Statistics computed on the expectation of denoised values may be biased due to spurious relationships between features that arise in the denoising process [37]. We used the expectation of denoised values for visualization (Supplementary Figure S5c-e). However, to address this concern for statistical analyses, the totalVI differential expression test uses distributions over the denoised values as opposed to testing directly on a denoised data matrix; we further describe this procedure below. For other analyses focused on the relationships between features, we developed a novel sampling method for denoised data that controls for nuisance variation while avoiding denoising-induced artifacts (Methods 4.3). We applied this method to construct feature-feature correlation matrices and found that totalVI preserved the independence of negative control genes, while the naive calculation of correlations on expected denoised values generated false positive correlations (Methods 4.9, Supplementary Figure S6a, b, d, e). These results lend confidence that downstream analysis with totalVI is not subject to spurious feature relationships that can arise from data denoising. When observing the correlations between proteins and their encoding RNA, we found that totalVI correlations were higher in magnitude than raw correlations across the majority of RNA-protein pairs (Supplementary Figure S6c, f). It is worth noting that the totalVI model has no explicit information about the relationship between any RNA-protein pairs, such that any correlation learned by the model is not predetermined by known RNA-protein relationships. In the sections below, we demonstrate how denoising protein expression within totalVI results in more accurate downstream analysis.

### 2.5 Integration of multiple datasets

As the number of CITE-seq studies grows, we anticipate increased interest in performing integrated analyses of datasets from different labs, experiments, and technologies; however, these categorical covariates often confound the shared biological signal between datasets. In the case of scRNA-seq data, methods like Seurat v3 [38] and Scanorama [39] were designed to correct expression values for these so-called batch effects and have demonstrated state-of-the-art results. These methods can also be extended to integrate CITE-seq datasets, but their application is limited. First, their corrected values are not designed to be used for differential expression testing or other related downstream tasks, since the uncertainty of the correction is not considered. Second, they require each dataset to have the same features, which may not be the case due to differences in antibody panels between datasets.

Integration is built into the totalVI model via an assumption of independence between the latent space and the batch. Consequently, the batch is treated as a covariate in the totalVI generative model, allowing for a statistically robust integration procedure. Furthermore, totalVI can handle datasets with different antibody panels due to a novel modification in which datasets with different proteins are processed using a fixed neural network architecture designed for the union of the proteins [40]. totalVI can also impute the expression values of missing proteins by making out-of-sample predictions, which incorporates the information learned from those proteins in the batches in which they were observed (Methods 4.3, [41]).

We benchmarked totalVI against state-of-the-art methods using general metrics designed to test how well datasets are mixed and how well the original structure of each dataset is maintained. We used these metrics to demonstrate that totalVI can effectively integrate datasets with equal and unequal protein panels. Finally, we benchmarked totalVI’s accuracy of predicting protein expression in cases where measurements are available in only one of the datasets.

We considered three metrics to quantify integration performance. First, the latent mixing metric quantifies how well the different batches mix in the low-dimensional latent space by considering the frequency of observed batches within local cell neighborhoods [42]. The second performance criterion, the measurement mixing metric, describes how well the different batches mix in the high-dimensional data space. This test uses the Mann-Whitney U statistic to evaluate the extent to which the expression of a given feature is systematically higher in one of the batches post-integration. Finally, as the mixing metrics could be trivially optimized by randomizing the data, we also tested how well the patterns of feature expression in every individual dataset are preserved in the integrated dataset.

In particular, we used the idea of autocorrelation, which measures for each feature the extent to which cells that are near each other in latent space have a similar expression level [43]. The feature retention metric summarizes the difference in the autocorrelation of features computed using the latent space of each individual dataset versus the integrated latent space. Thus, it tests whether the integrated latent space captures similar genes and proteins as the latent space of each dataset in isolation (Methods 4.10).

We demonstrated totalVI’s integration capabilities on two of the spleen and lymph node datasets (SLN111-D1, SLN208-D2), which were generated with different panels containing either 111 or 208 antibodies (111 is a subset of the 208). For these datasets, a linear dimensionality reduction with principal components analysis (PCA) revealed batch as a major source of variation (Figure 3a). We compared totalVI to Seurat v3 [38] and Scanorama [39], which, like totalVI, produce both batch corrected expression values and an integrated latent space. We benchmarked two versions of totalVI – totalVI-intersect, in which protein features were intersected, and totalVI-union, in which the union of protein features was used as input.

**Figure 3:**
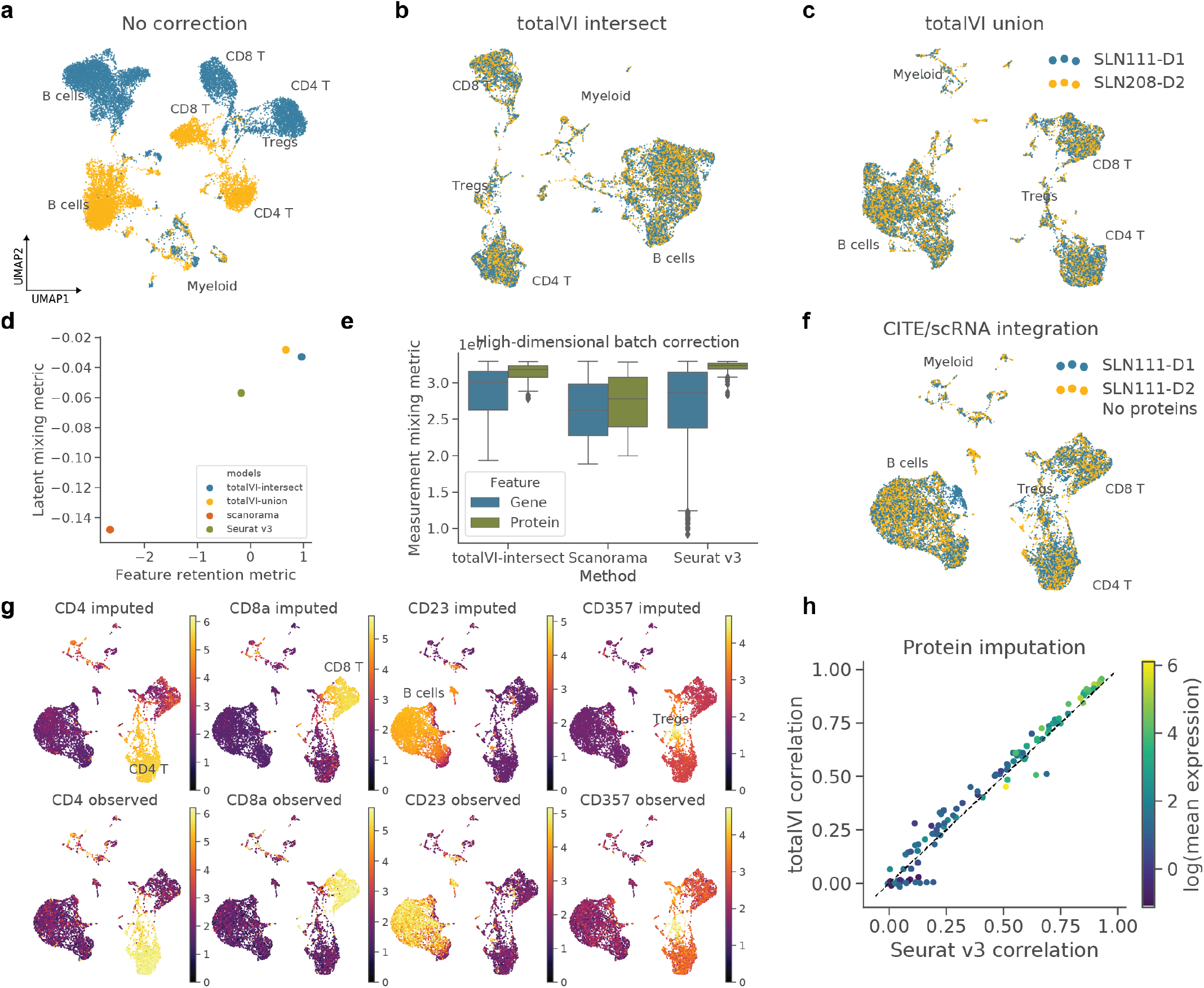
Benchmarking of integration methods for CITE-seq data. **a-c**, UMAP plots of SLN111-D1 and SLN208-D2 with no integration (PCA of paired data with intersection of protein panels), and after integration with totalVI-intersect, in which the protein panels were intersected, and totalVI-union, in which the unequal protein panels were preserved, colored by dataset. **d, e**, Performance of integration methods based on three metrics: **(d)** latent mixing metric, feature retention metric and **(e)** measurement mixing metric (higher values are better for each; Methods 4.10). **f**, UMAP plot of SLN111-D1 integrated with SLN111-D2 (proteins held out) by totalVI. **g**, UMAP plots colored by totalVI imputed and observed protein expression (log scale) of key cell type markers (range 0-99th percentile of held-out values for each protein). **h**, Pearson correlation of imputed versus observed protein expression (log scale) for totalVI-union and Seurat v3 colored by average expression of the protein.

For each of these methods, we computed UMAP embeddings [44] of their latent spaces (Figure 3b, c), as well as the three evaluation metrics. Generally, we found that after integration, cells of similar types were co-located in the latent space, as evidenced by the shared expression of key marker proteins like CD4, CD8a, and CD19 (Supplementary Figure S7). In the latent mixing and feature retention metrics, totalVI outperformed the other methods, while comparing favorably in the measurement mixing metric (Figure 3d, e). totalVI-union and totalVI-intersect performed similarly, indicating that the presence of missing data did not diminish totalVI’s integration capabilities. We also repeated this analysis on two public datasets of PBMCs (PBMC10k [31], PBMC5k [45]) and observed similar performance for totalVI (Supplementary Figure S8a-f).

Since totalVI-union can integrate datasets with different protein panels, we reasoned it would be feasible to integrate a CITE-seq dataset with a standard scRNA-seq dataset that has not measured proteins and impute the missing protein measurements. We assessed this by integrating SLN111-D1 and SLN111-D2, where we held out the proteins of SLN111-D2. We first observed that totalVI can learn a biologically meaningful integrated latent representation despite the large amount of missing data (Figure 3f). Indeed, the localization of observed protein expression in the latent space revealed the same broad immune cell types like B cells, CD4 T cells, CD8 T cells, Tregs, and myeloid cells. Next, we decoded the latent representations of cells from SLN111-D2 conditioned on them being in SLN111-D1, where protein data was observed, and reported the mean of the negative binomial mixture for each protein as the imputed protein values (Methods 4.3). For key cell type marker proteins, totalVI-imputed proteins shared similar patterns of expression as the held-out observed proteins (Figure 3g).

To further quantify imputation accuracy, we computed the Pearson correlation between imputed and observed protein values on the log scale. We did not correct for background in this analysis since the comparison is to the observed data. We compared totalVI to Seurat v3, which imputes protein values based on smoothing of protein values from mutual nearest RNA neighbors. totalVI had higher correlation than Seurat v3 for the majority of the proteins (Figure 3h). As expected, imputation was inaccurate in both methods for proteins with low counts, which may only have foreground signal in few cells, or were poorly detected. We observed similar results when applying this task to PBMCs (Supplementary Figure S8g, h). Taken together, these results suggest that imputed proteins may be used as a proxy for real protein measurements in other downstream tasks.

### 2.6 Differential expression

totalVI can leverage its estimates of uncertainty from a single model fit to detect differentially expressed features between sets of cells while controlling for noise and other modeled technical biases. In a test between two sets of cells, totalVI estimates the posterior odds that any feature is differentially expressed (using Bayes factors [13, 46, 47]; Methods 4.3). Here, the Bayesian equivalent of a null hypothesis for a particular feature is that the log fold change (LFC) of expression between the two sets is contained within a small interval centered at zero. Likewise, the probability that the LFC is outside this interval is considered the *probability of differential expression* (DE). The Bayes factor thus quantifies the degree to which the data support the hypothesis that a feature is differentially expressed versus the hypothesis that it is not differentially expressed. A distribution over the LFC is estimated using posterior samples of *z_n_* and the subsequent data likelihood parameters (Figure 1b), adjusted for sequencing depth (RNA), background signal (protein), and batch effects (both). We used the median of this LFC distribution as an estimate of effect size.

To evaluate totalVI as a framework for DE analysis in the common scenario of multiple experiments, we integrated all four spleen and lymph node datasets (SLN-all; totalVI-intersect). totalVI provided a descriptive representation of this data, as clusters of cells in the joint latent space corresponded to immune cell types or states. Indeed, we found consistent differences in the expression of known immune cell markers, thus allowing us to manually annotate each cluster with cell type labels (Figure 4a, Supplementary Figure S9, Methods 4.11). These annotations were consistent with the latent space derived with totalVI-union (Supplementary Figure S10), and we used them throughout this section.

**Figure 4:**
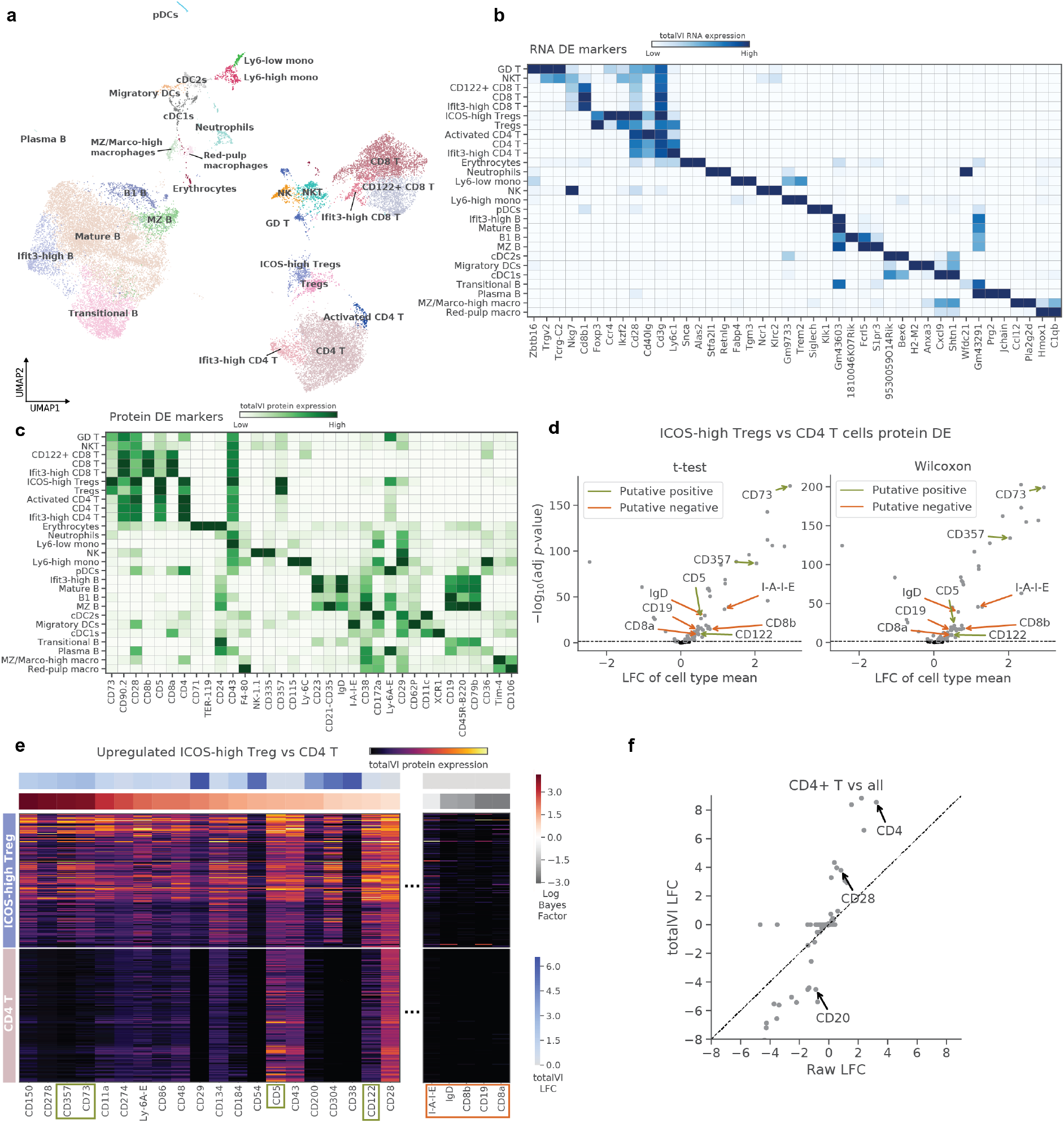
totalVI identifies differentially expressed genes and proteins. totalVI intersect was applied to the SLN-all dataset. **a**, UMAP plot of SLN-all, after clustering and annotating the data (Methods 4.11). **a, b**, Heatmap of markers derived from one-vs-all tests for **(a)** RNA and **(b)** proteins. For each cell type, we display the top three protein markers and top two RNA markers in terms of LFC. **d**, Volcano plot of protein differential expression test between ICOS-high Tregs and CD4 T cells for a Welch’s t-test and Wilcoxon rank-sum test. Putative positives and negatives are denoted by green and orange arrows, respectively. Significant proteins are colored in grey, all others are in black. **e**, totalVI protein expression for proteins (columns) upregulated in ICOS-high Tregs versus CD4 T cells. Cells (rows) are ordered by cluster, and subsampled to be equal in number per cluster. Columns are normalized in the range [0, 1]. The left section in the heatmap contains all the proteins called differentially expressed by totalVI with a positive log fold change. Proteins are sorted by Bayes factor (significance). The rightmost section contains the putative negatives, which are not called differentially expressed by totalVI. **f**, Comparison of log fold changes estimated by totalVI and observed in the raw data from a one-vs-all test of CD4 T cells.

Beyond markers used for annotation, we found that a totalVI one-vs-all DE test (in which one cell type is compared to all others) identified many additional features as differentially expressed by one or more subsets of cells (Methods 4.12, Figure 4b, c). For example, totalVI identified the gene Klrc2 as differentially expressed in both natural killer (NK) cells and gamma/delta T cells, which has previously been shown to be upregluated in these populations relative to alpha/beta T cells [48]. On the protein side, totalVI identified CD335 (NKp46) as among the top protein markers for NK cells, which is a canonical marker used for sorting [49], and CD43, which is associated with the development of NK cells [50].

Overall, the Bayes factors inferred by totalVI for the RNA data were highly correlated with those produced by scVI (Supplementary Figure S11a), which has been independently evaluated [47]; therefore, we focused on evaluating the protein DE test. For the protein DE test, we focused on testing for accurate detection of true positive and negative cases of DE and reproducibility across datasets. Throughout, we compared totalVI to two methods – a Welch’s t-test applied to the log-transformed protein counts and a Wilcoxon rank-sum test. These general methods are commonly used for scRNA-seq data, so we used them as baselines for protein DE testing (Methods 4.12).

We first evaluated the extent of false positives using isotype control antibodies. As isotype controls lack target specificity, differences in the abundance of isotype controls between cell types may stem from differences in background or other technical sources of variation. We applied totalVI to the SLN208-D1 dataset, which contained nine isotype controls, and performed a one-vs-all DE test. In all but one of the comparisons in the one-vs-all test, totalVI called zero of the isotype controls as differentially expressed, outperforming the baseline methods. totalVI also outperformed a version of itself (totalVI-wBG) in which the protein background component is not subtracted (Supplementary Figure S11b).

To gain further insight into the extent of false positive and false negative DE calls, we compared ICOS-high regulatory T cells (ICOS-high Tregs) and conventional CD4 T cells from SLN-all in a one-vs-one DE test. This test is challenging because these two cell types share many of the same upregulated and downregulated features when compared with other immune cell types. Our analysis was based on a list of putative positives and negative surface proteins curated from previous studies that used flow cytometry. In this test, we expected CD73, CD357 (GITR), CD122, and CD5 to be upregulated (positives) in ICOS-high Tregs relative to conventional CD4 T cells [51–54].

We found that all three methods identified these positives as significantly upregulated; however, the two baseline methods also incorrectly called all putative negatives as upregulated, which included I-A/I-E (MHC II), IgD, CD19, CD8b, and CD8a, which have no expected expression in either of these cell types (Figure 4d). Globally, the two baseline methods called the majority of the proteins in the panel (78/111 proteins in both cases) as differentially expressed, many of which are likely the result of differences in background. Filtering proteins by the observed LFC in the baseline methods may reduce false positives in these cases, but the increased accuracy would still be limited (e.g., CD5 and IgD had a similar LFC and therefore could not be distinguished; Figure 4d). The totalVI test, on the other hand, correctly classified the putative negatives and putative positives in our curated list (Figure 4e), while calling 28 proteins differentially expressed in total. To further support the utility of removing background from the protein data, we performed this test using totalVI-wBG, which improved upon the baseline methods, but also falsely called I-A/I-E (MHC II) and CD8b as positives (Supplementary Figure S12a).

To test for reproducibility across biological replicates, we applied each method separately to the SLN111-D1 and SLN111-D2 datasets in a one-vs-all DE test. Comparing the results obtained from each dataset (Bayes factors for totalVI and adjusted *p*-values for the benchmark methods) by Spearman correlation, we found that totalVI had the highest consistency between replicates (Spearman’s *ρ* = 0.91; Supplementary Figure S11c-e). We also tested for reproducibility across experimental designs: one in which the two CITE-seq datasets (SLN111-D1, SLN111-D2) had equal protein panels, and another in which proteins were measured in only one of the datasets. We used totalVI to integrate the two batches in each scenario and conducted one-vs-all DE tests. The resulting Bayes factors were reproducible between the two scenarios (Spearman’s *ρ* = 0.86; Supplementary Figure S11f).

Finally, the LFC estimates calculated by totalVI also better captured the underlying biological signal. For example, in a test of CD4 T cells vs all from SLN-all, the canonical marker CD4 had a higher LFC than in the raw data (Figure 4f). Additional markers like CD28 (T cell marker) and CD20 (B cell marker), which we previously highlighted as having highly overlapping foreground and background components, had respectively higher and lower LFCs compared to LFCs derived from the raw data. This increase in contrast is driven by totalVI’s ability to probabilistically remove the background.

### 2.7 Interpretation of totalVI latent space

Deep-learning-based methods for dimensionality reduction tend to rely on “black-box” models, making it difficult to interpret the coordinates of their inferred low-dimensional latent spaces. For example, for autoencoder-based single-cell methods like scVI or DCA [13, 24], there is no straightforward way to determine which expression programs are associated with each dimension of the latent space. This is in contrast to linear methods like PCA, GLM-PCA or LDVAE [27, 55], where each latent dimension is associated with a loading vector that describes the contribution made by each feature and thus enables direct interpretation. The interpretability of linear methods, however, comes at the expense of reduced capacity to fit complex data such as that obtained by scRNA-seq [27].

In totalVI, although the relationship between the latent space and the observation space is non-linear, the model still provides a way to relate each latent dimension to the expression of individual genes and proteins. Specifically, the latent vector *z_n_* associated with each cell *n* follows a logistic normal distribution, such that the values in *z_n_* are non-negative and sum to one. Thus, each cell represents a mixture over the different vertices of a probability simplex [26, 56, 57]. Each of these vertices, commonly referred to as archetypes, corresponds to a point in the latent space of totalVI, where the value of *z* is 1 in one coordinate and 0 in all others. The individual cells can thus be thought of as making a tradeoff, choosing the archetypes that best describes their function. The archtypes, on the other hand, represent a summary of expression programs, the combination of which characterizes a cell.

The expression profile associated with each archetype can be obtained using the decoder network of totaVI, as it is trained to map points in latent space to the parameters that underlie the distribution of each gene and protein (Figure 1b). To explore these archetypal gene and protein expression profiles, we decoded the archetypal points from the SLN-all dataset (Supplementary Figure S13a, b). We found that some archetypal points in this dataset correspond to specific cell types, whereas others corresponded to more global sources of variation that are not constrained to one cell type (Supplementary Figure S14a).

For example, archetype 16 was associated with high protein expression of CD93 and CD24, which mark the transitional B cell subset (Supplementary Figure S14b). Conversely, archetype 7 was associated with interferon-response genes such as Ifit3 and Isg20 and reflected within cell type variability in several subsets, including CD4 and CD8 T cells, B cells, Ly6-high monocytes and neutrophils (Supplementary Figures S14c and S15). Therefore, archetypal analysis enables a data-driven alternative to clustering for characterizing the heterogeneity observed both between and within cell subsets.

Archetypal analysis can also provide insight into the inner-workings of the model. In particular, we sought to evaluate the contribution of the protein data to the model and to the inferred latent space. To this end, we computed for every archetype, which percent of its top associated features are proteins. The representation of proteins amongst the top ranking features is higher than naively expected by their share of the feature population (Supplementary Figure S13c). Furthermore, the top genes often did not encode the top proteins in each archetype. Together, these results suggest that the protein data significantly influences the locations of the cells in the latent space.

### 2.8 Characterization of B cell heterogeneity in the spleen and lymph nodes with RNA and proteins

We next demonstrate how a joint representation of RNA and protein can be used to characterize cell identities within a specific immune compartment and in the context of multiple samples. To this end, we used the totalVI-intersect model fit on the SLN-all dataset and focused on the B cell population (Methods 4.14, Figure 5a).

**Figure 5:**
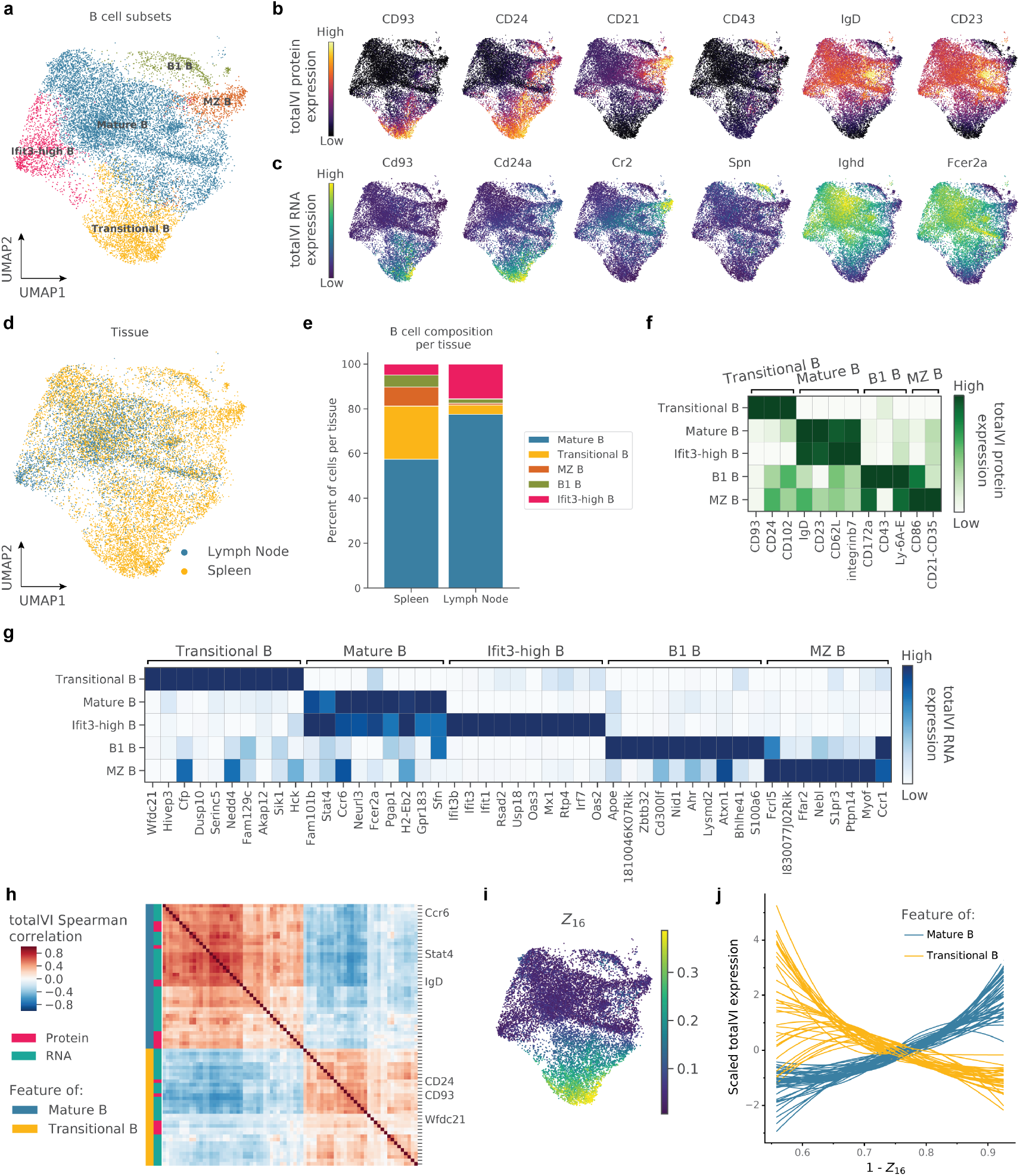
Characterization of B cell heterogeneity in the spleen and lymph nodes with RNA and protein. totalVI-intersect was applied to the SLN-all dataset. Data were filtered to include B cells. **a**, UMAP plot of totalVI latent space labeled by cell type. **b, c**, UMAP plots of totalVI latent space colored by **(b)** totalVI protein expression of six marker proteins and **(c)** totalVI RNA expression of the six genes that encode the corresponding proteins in (b). **d**, UMAP plot of totalVI latent space labeled by tissue. **e**, Cell type composition per tissue. **f, g**, totalVI one-vs-all differential expression test on B cell subsets filtered for significance (Methods 4.12) and sorted by the totalVI median LFC. **(f)** The top three differentially expressed proteins per subset and **(g)** the top ten differentially expressed genes per subset, arranged by the subset in which the LFC is highest. **h**, totalVI Spearman correlations in transitional B cells between RNA and proteins, which were selected as described in Methods 4.14. Features were hierarchically clustered and are labeled as either RNA or protein, and by the cell type with which the feature is associated. **i**, UMAP plot of totalVI latent space colored by *Z*_16_ (the totalVI latent dimension associated with transitional B cells). **j**, totalVI expression of features in (h) as a function of (1 − *Z*_16_). Each feature was standard scaled and smoothed with a loess curve.

We start with characterizing cell identities using prior biological knowledge by curating a set of six surface markers that are commonly used for isolating B cell subsets (Table 4), and visualizing their expression on the UMAP representation of the totalVI latent space (Figure 5b). We found that these markers stratified the B cells into groups that were largely consistent with unsupervised clustering (Methods 4.14). RNA expression of these markers followed similar patterns to the proteins they encode (Figure 5c). This analysis identified several B cell subsets, including transitional B cells (marked by CD93 and CD24) and mature B cells (marked by IgD and CD23). Additionally, we detected clusters that represent the non-follicular subsets of B1 and marginal zone (MZ) B cells (marked by CD43 and CD21, respectively).

Comparing the composition of the different B cell subsets in the spleen and in the lymph nodes (Figure 5d), we found that these tissues contained overlapping but not identical subsets of B cells in a manner that was consistent with previous studies (Figure 5e, [58, 59]). In particular, clusters spanned the developmental range from recent bone-marrow emigrants in the splenic transitional B cell subset to a mature subset that had substantial representation in both the spleen and lymph nodes. As expected, the B1 and MZ B cell subsets were found primarily in the spleen.

Moving beyond visualization of known markers, we used totalVI in a more unbiased approach to quantify the differences between the B cell clusters with a one-vs-all DE test (Methods 4.12). With a single DE test, totalVI was able to provide information about these B cell subsets using both their protein and RNA expression (Figure 5f, g). As expected, the totalVI DE test found the set of six known surface markers in Figure 5b to be among the top three protein markers defined by the corresponding one-vs-all test (Figure 5f). Most RNA molecules encoding the marker proteins were also differentially expressed in the respective one-vs-all test. The complete set of differentially expressed genes, however, contained many additional molecules at similar or higher level of significance. This list included informative genes whose products are not necessarily present on the surface of the cell, such as the transcription factor Bhlhe41 that marks B1 B cells (Figure 5g, [60]).

More globally, the protein data combined with the transcriptome-wide view enabled a more refined characterization of variation within the four major sub-populations identified above by surface markers. One example of this is a sub-population of mature B cells, labeled here as Ifit3-high B cells. The Ifit3-high B cells expressed all of the protein and RNA markers of mature B cells and could not be clearly distinguished from the remaining mature B cells based on protein data alone (maximal LFC across all proteins was less than 0.19). Nevertheless, based on transcriptome-wide DE analysis, this cluster could still be distinguished as a sub-type of mature cells by the elevated expression of a host of interferon response genes (Figure 5g). This observation was also supported by a gene signature analysis with Vision [61], which identified two interferon response signatures that were enriched in the Ifit3-high B cell cluster (Methods 4.14, Supplementary Figure S15a, b). The expression of interferon response genes was not necessarily expected in this steady state condition (i.e., with no induced inflammation), however, we found the Ifit3-high B cell cluster as well as Ifit3-high T cell clusters to be represented in both biological replicates, and therefore took it to capture part of the biology in the SLN-all dataset (Supplementary Figure S15c, d).

As a second example, we explored the variability within the subset of transitional B cells and its relationship with the process of B cell development. Interestingly, latent dimension 16 (*Z*_16_) captured a gradual transition within this cluster: from a small population of Rag1 expressing cells (indicating early development [58]) to cells that were closer to the mature cluster (Figure 5i, Supplementary Figure S16b, c). We used totalVI to explore how development from transitional to mature B cells may be associated with coordinated changes in gene and protein expression. To do this, we calculated the totalVI Spearman correlations separately within transitional and mature B cells for a set of features that distinguished between the two subsets (Methods 4.14). Within the transitional B cells, hierarchical clustering clearly stratified this feature set into two anti-correlated modules, one corresponding to proteins and RNA associated with transitional B cells and the other to mature B cells (Figure 5h). These modules, however, were less apparent in a hierarchical clustering of these features in mature B cells (Figure S16a), indicating that the apparent coordination may be a characteristic of the transitional state. Focusing on the transitional B cells, we found that the features in the two modules significantly correlated with the axis of maturation captured by latent dimension 16 (Supplementary Figure S16d). For cells progressing along this axis, the expression of features in the transitional module decreased and those in the mature module increased (Figure 5j, Methods 4.14). These results therefore point to a program of transitional B cell maturation that consists of coordinated activation and repression of multiple genes and surface proteins, leading to a gradual transition in cell state.

Taken together, these analyses demonstrate the ability of totalVI to reveal biologically-relevant patterns in feature expression and to identify differentially expressed RNA and surface proteins that could independently inform future experiments and analyses. While the proteins measured did not cover the entire proteome, they provide a crucial link to the surface marker literature that has traditionally defined cell types and present an opportunity to identify new markers for which we have antibodies. Many of the most defining RNA features in B cells do not encode surface proteins, but nonetheless are likely to play defining roles in cell state through transcriptional regulation. Through this multi-level analysis based on a combined RNA-protein view in a joint latent space, we could thus obtain a more complete picture of cell identity.

## 3 Discussion

We have developed totalVI, a scalable, probabilistic framework for end-to-end analysis of paired transcriptome and protein measurements in single cells. totalVI couples a low-dimensional representation of the paired data with disparate downstream analysis tasks such as visualization, identification of cell types and cell states, integration of datasets, data denoising, imputation of missing protein measurements, and differential expression testing. totalVI builds upon previous autoencoder-based methods that have been successfully applied to a variety of common tasks in single-cell transcriptomics data analysis [13, 24]. To handle the addition of proteins, totalVI assumes that RNA and protein measurements are generated from the same latent space of cells that captures their state – an assumption shared by a number of multi-omics methods [36, 62, 63].

A distinction of totalVI is that it explicitly models modality-specific technical factors that contribute to the observed data, obviating the need for a preprocessing step of normalization, which may induce biases [64]. For proteins specifically, totalVI leverages the concept of mixture density networks [65], establishing for each protein within each cell a negative binomial mixture in which the component with smaller mean corresponds to background. We demonstrated that this mixture could be effectively used to identify protein measurements that are the result of protein background and likewise those that correspond to real surface proteins. This enabled a denoised view of the data, which led to more accurate differential expression results.

We demonstrated how totalVI can be used to conduct a tiered analysis in our characterization of the heterogeneity of B cells in the murine spleen and lymph nodes. Based on a joint latent space containing both RNA and protein information from integrated datasets, we identified clusters, visualized and interpreted those clusters using known B cell surface markers, and performed a higher-resolution differential expression analysis that revealed a combination of RNA and protein markers that characterized each cluster. This analysis demonstrates an approach that can be applied to other biological systems to identify new cell types and surface markers, and provide a more detailed understanding of cell states and molecular processes through the combination of transcriptomic and proteomic information.

Going beyond the characterization of cell types, totalVI can uncover relationships between RNA and protein molecules within a cell. For example, totalVI could be used to investigate the relationship between the level of an RNA transcript and the level of its encoded protein in different biological settings, which remains an open question [66]. Previous works aiming to quantify RNA-protein relationships with paired measurements have reported weak positive correlations [8, 9]. However, it is unclear to what extent these low correlations can be attributed to technical noise in RNA and protein measurements or to biological factors such as transcription, translation, and post-translation dynamics. We found that the totalVI correlations were higher in magnitude than raw correlations across the majority of RNA-protein pairs, suggesting that the low correlations between RNA and proteins observed in previous studies could have been the result of technical noise. Future work quantifying correlations and regulatory relationships between RNA and protein features could inform our understanding of co-regulated gene networks, signal transduction pathways, or transcription and translation dynamics [12].

totalVI is the culmination of an iterative design process that consists of defining a candidate generative process and inference procedure, and then criticizing the model’s performance [22]. This process encodes domain knowledge into the hierarchical model that disentangles sources of variation. In modeling the protein data, we added components like protein background parameters that reflect our understanding of the CITE-seq experimental data-generating process, which we further discuss in Appendix A. Furthermore, we addressed a number of practical concerns with neural-network-based approaches, including their interpretability and sensitivity to hyperparameter choices (e.g., number of layers, learning rate, etc.) [67]. To address interpretability, we constrained the totalVI latent space to be the probability simplex, which enabled us to link variation in latent dimensions to particular genes and proteins with archetypal analysis [26]. This revealed that some of totalVI’s latent dimensions were associated with specific cell types, while others corresponded to more global sources of variation, which could be observed within several cell types. With regard to the second concern, we used a default set of hyperparameters across all datasets and analysis tasks (Appendix C), demonstrating state-of-the-art results.

While the totalVI model was designed in consideration of the CITE-seq experimental protocol, we envision totalVI being used to inform experimental design by optimizing methods that affect signal-to-noise (Appendix A). For instance, totalVI could help identify optimal antibody titrations that improve the identification of foreground and background in protein measurements. totalVI could also be used to identify sequencing depths for RNA and protein libraries that balance the information gained per measurement in various analysis tasks with the cost of additional sequencing [68, 69].

totalVI is available as part of the scvi software package, thus allowing for a consistent programmatic interface and workflow for users analyzing different types of single-cell data. This software package includes tutorials that demonstrate functionality described here, as well as the interaction between totalVI and Scanpy [70], a popular Python-based single-cell analysis pipeline. Since totalVI processes datasets in mini-batches of hundreds of cells, it has a small memory footprint and therefore can typically be used with free cloud computing environments like Google Colab. Overall, the accessibility and scalability of totalVI make it readily available to contribute to and extract knowledge from growing community efforts such as the Human Cell Atlas [4].

The flexibility and scalability of totalVI make it easily applicable to future datasets with larger protein panels, and enable extensions that incorporate additional paired measurements. For example, we expect totalVI to naturally handle intracellular proteins measured with barcoded antibodies. Further additions of modalities like chromatin accessibility [71] or clonotype features [72] is made straightforward within the totalVI codebase with consideration of the modality-specific likelihood. By combining multiple views of cellular processes, totalVI could reveal a more complete picture that redefines cell states and elucidates mechanistic relationships between molecular components of the cell.

## Supporting information

Supplementary Data: Antibodies

## Acknowledgements

We thank Ellen Robey, Lydia Lutes, and Derek Bangs for help designing experiments. We thank the BioLegend Inc. and their proteogenomics team, especially Bertrand Yeung, Andre Fernandes, Qing Gao, Hong Zhang, Tse Shun Huang, for providing reagents and expertise and for help with sample preparation, library generation, and sequencing of CITE-seq libraries. We thank David DeTomaso for general data analysis advice, and Pierre Boyeau and Achille Nazaret for help with integrating totalVI in the scvi package. We thank members of the Streets and Yosef laboratories for helpful feedback. Research reported in this manuscript was supported by the NIGMS of the National Institutes of Health under award number R35GM124916 and by the Chan-Zuckerberg Foundation Network under grant number 2019-02452. A.G. is supported by NIH Training Grant 5T32HG000047-19. Z.S. is supported by the National Science Foundation Graduate Research Fellowship. A.S. and N.Y. are Chan Zuckerberg Biohub investigators.

## Author contributions

A.G. and Z.S. contributed equally. A.G., Z.S., A.S., and N.Y. designed the study. A.G., Z.S, R.L., J.R., and N.Y. conceived of the statistical model. A.G. implemented the totalVI software with input from R.L. K.L.N. designed and produced antibody panels and provided input on the study. Z.S. designed and led experiments with input from A.S. and N.Y. A.G. and Z.S. designed and implemented analysis methods and applied the software to analyze the data with input from A.S. and N.Y. A.S. and N.Y. supervised the work. A.G., Z.S., R.L., J.R., A.S., and N.Y. participated in writing the manuscript.

## 4 Methods

### 4.1 The totalVI model

totalVI estimates a conditional distribution for cell *n*, *p_ν_* (*x_n_*, *y_n_ s_n_*), in which *x_n_* is the *G*-dimensional vector of observed RNA counts (*G* genes), *y_n_* is the *T*-dimensional vector of observed protein counts (*T* proteins) and *s_n_* is the *B*-dimensional one-hot vector describing the batch index (experiment identifier). In total, there are *N* cells. We use *ν* to refer to the set of all generative parameters, which are described throughout this section. This distribution is estimated using the framework of variational autoencoders (VAE; [25]).

We begin by describing the generative process, for which a graphical summary is in Figure S17 and an algorithmic summary is in Algorithm 1. We then describe the inference procedure, as well as how downstream analysis tasks are directly linked to posterior queries of the model.

#### Algorithm 1

The totalVI generative model. The gamma distribution is parameterized by its shape and mean. Let *v* be the set of model parameters described here. A dataset has *G* genes and *T* measured proteins.

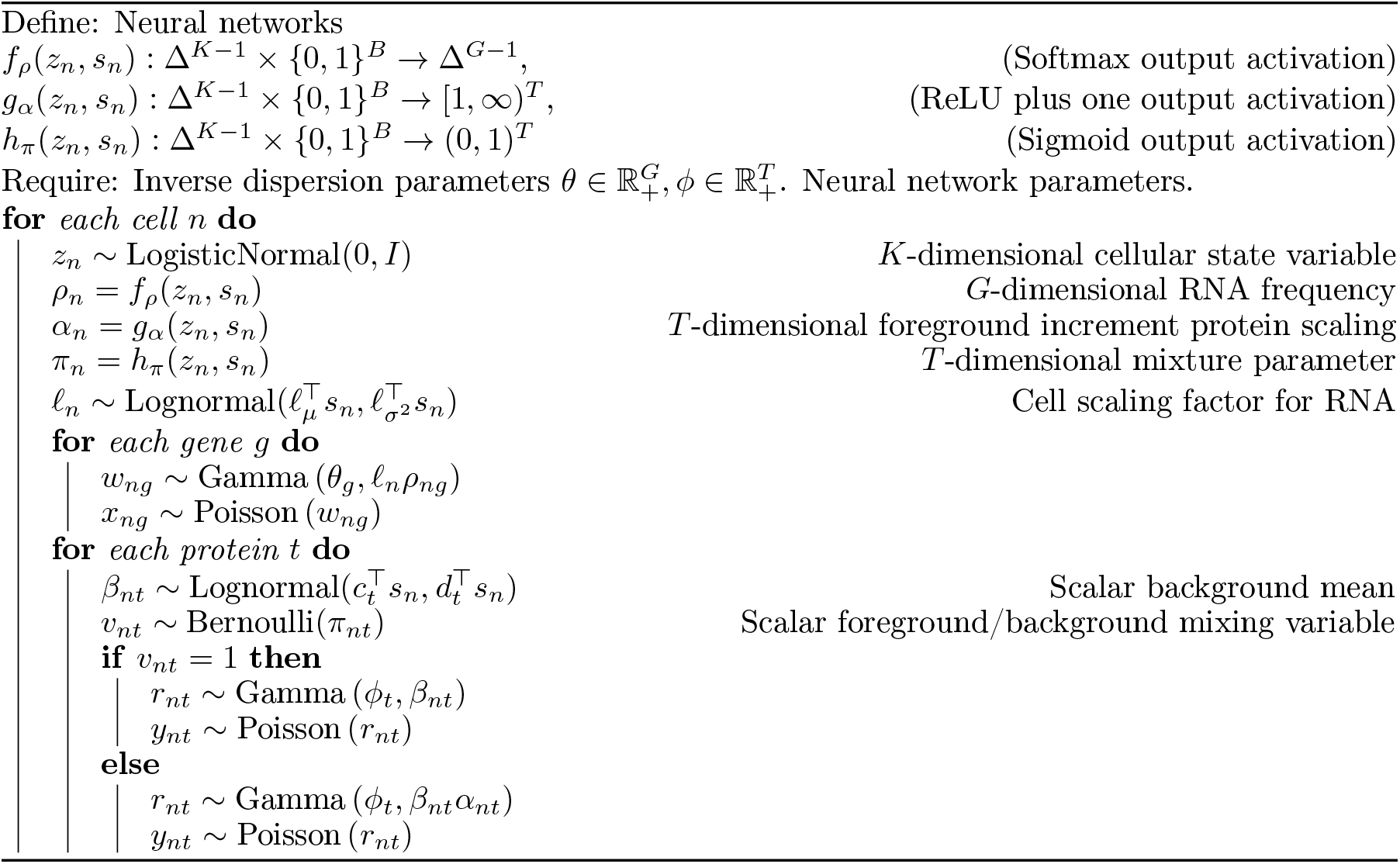

#### Priors

The latent cell representation *z_n_* ∼ LogisticNormal(0, *I*), where the logistic normal distribution is a distribution over the probability simplex. This specification enables cells to be interpreted with archetypal analysis (Methods 4.13). Typically in VAEs, *z_n_* follows an isotropic normal distribution, which is chosen for computational convenience [25]. In this setting, a logistic normal distribution arises as transforming a sample from a normal distribution with a softmax function. A full description is in Appendix C. For all experiments, we set *z_n_* to 20 dimensions. The latent RNA size factor 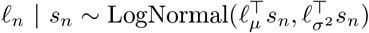, where *ℓ*_μ_ ∈ ℝ^*B*^ and 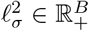 are set to the empirical mean and variance of the log RNA library size (defined as total RNA counts of a cell) per batch. We use a protein-specific prior for the protein background intensity, where 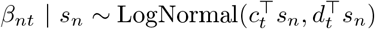. The parameters for the background intensity, *c_t_* ∈ ℝ^*B*^ and *d_t_* 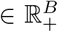, are protein specific and are treated as model parameters learned during inference. This prior is motivated by the observation that some component of the background is due to ambient antibodies. A prior can also be thought of as regularizing the posterior distribution, thus reducing the influence of outliers [73].

#### RNA likelihood

Given *z_n_*, *ℓ_n_*, and *s_n_*, an observed expression level *x_ng_* follows a negative binomial distribution, which we present here as a Gamma-Poisson mixture:

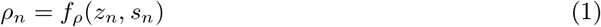

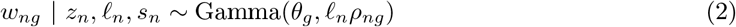

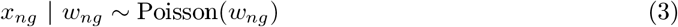

The gamma distribution is parameterized by its shape and mean. The mean is equal to *ℓ_n_ρ_ng_*, where *ℓ_n_*, a scaling factor, is multiplied by *ρ_ng_*, interpreted as a normalized gene frequency (because *ρ_n_* is nonnegative and sums to one). *ρ_n_* is the output of a neural network *f_ρ_*, which takes *z_n_* and *s_n_* as input (Algorithm 1).

Integrating out *w_ng_* results in the following conditional distribution:

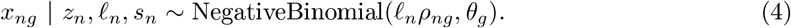

The parameter *θ_g_*, which is the shape of the gamma distribution, is also the inverse dispersion of the negative binomial. Further details on this connection are in Appendix B. We perform inference on the model with *w_ng_* integrated out. We also treat *θ_g_* as a model parameter learned during inference. Overall, this likelihood is equivalent to that presented in scVI [13], without zero-inflation.

#### Protein likelihood

To capture observed protein counts arising from the background or foreground, we model *y_nt_* with a negative binomial mixture, given *z_n_*, *β_n_* and *s_n_*. This conditional distribution is described by the following process:

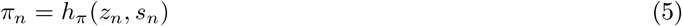

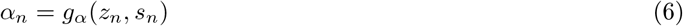

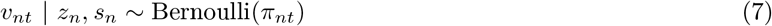

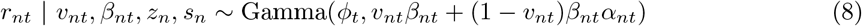

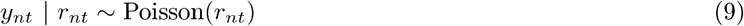

Here *v_nt_* controls which mixture component generates the counts. Its parameter, *π_nt_*, is the output of the neural network *h_π_*(*z_n_*, *s_n_*). Notably, *α_nt_*, which is the output of the neural network *g_α_*(*z_n_*, *s_n_*), is greater than one. This ensures that one of the mixture components is always larger than the other, allowing us to interpret one component as background and one component as foreground. Furthermore, *π_nt_* is interpreted as the probability that any cell-protein pair has observed counts due to background alone. For one mixture component, *y_nt_ z_n_*, *β_nt_*, *s_n_*, *v_nt_* follows a negative binomial distribution, as can be seen by integrating out *r_nt_*. Finally, integrating out *v_nt_* too shows that *y_nt_* given *z_n_* and *s_n_* follows a negative binomial mixture distribution, where *ϕ_t_* is a protein-specific inverse dispersion parameter.

### 4.2 Inference for totalVI

Here we describe the inference procedure for totalVI except for the case of missing data (i.e., totalVI union). The model evidence, *p_ν_* (*x*_1:*N*_, *y*_1:*N*_ | *s*_1:*N*_), cannot be computed as the integrals are analytically intractable, so Bayes rule cannot be directly applied to find a posterior distribution. Therefore, we use variational inference [74] to approximate the posterior distribution with a distribution having the following factorization:

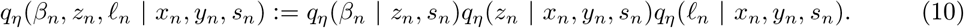

Here *η* is the set of parameters of an inference network, commonly called the *encoder* – a neural network that takes a cell’s combined expression as input and outputs the parameters of the approximate posterior (e.g., mean and variance). Factors of the posterior approximation share the same family as their respective priors (e.g., *q*(*β_n_* | *z_n_*, *s_n_*) is lognormal). As described previously, we integrate out the latent variables *v_nt_*, *r_nt_* and *w_ng_* (Algorithm 1), yielding *p_ν_* (*y_nt_* | *z_n_*, *β_nt_*, *s_n_*), which is a mixture of negative binomials and *p_ν_* (*x_ng_* | *z_n_*, *s_n_*, *l_n_*), which is a negative binomial distribution.

We optimize the evidence lower bound (ELBO) [74] of log *p_ν_* (*x*_1:*N*_, *y*_1:*N*_ | *s*_1:*N*_) with respect to the variational parameters *η* and model parameters *ν* using stochastic gradients [25]. In other words, we learn the model parameters and posterior distributions simultaneously. In the VAE framework, the generative neural network is referred to as the *decoder*. Each iteration of training consists of randomly choosing a mini-batch of data (256 cells), estimating the ELBO based on this mini-batch, and updating the parameters via automatic differentiation operators. The terms corresponding to Kullback-Leibler divergences of the ELBO (see Appendix C) follow a deterministic warm-up scheme [75], which helps to avoid shallow local maxima. We use the Adam optimizer [76] with weight decay to update the model parameters. Learning rate reductions and early stopping are performed based on the ELBO of a validation set.

All neural networks are feedforward and and use standard activations (e.g., exponential, softmax, sigmoid) to encode the variational and generative distributions. We use the same hyperparameters for all our experiments. Appendix C gives further implementation details.

#### Inference in the case of missing proteins

Here we adapted the training procedure from [77] to handle missing protein data. As any single batch may correspond to an experiment that used a different protein panel (or no proteins in the case of a scRNA-seq experiment), the missingness of protein features depends on the batch index *s_n_*. Further, suppose all batches share the same set of genes. Across all batches, there are *T* proteins. For cell *n*, we denote the observed protein expressions 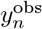 and the unobserved protein expressions 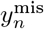. The log likelihood of the observed data decomposes as

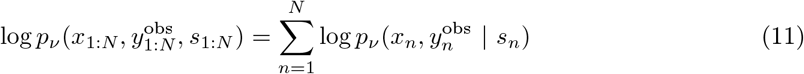

The generative process for the observed data is the same as in Algorithm 1, with appropriate modification to only generate the features present in a particular batch. Thus, *ν* is the same set of model parameters described previously. Again, we use variational inference to approximate the posterior distribution with the distribution in Equation 10. In fact, all approximate posteriors share the same encoder parameters *η*. We optimize the ELBO of Equation 11 similarly to the procedure used when there is no missing data (i.e., we optimize the ELBO with respect to the model parameters *ν* and variational parameters *η*). To handle mismatched dimensions in the encoder, we substitute zeros for missing proteins, and for the decoder, we only calculate the ELBO terms corresponding to observed data [40]. Therefore, this procedure naturally extends to the case when there is no observed protein data for a cell *n*, which would be the case when the cell is obtained from a scRNA-seq experiment. Since the quality of missing protein imputation depends on (i) the goodness of fit of totalVI to the protein for the data in which it was observed and (ii) the statistical distance of the aggregated posterior distributions of *z_n_* for each of the batches [77, 78], we add a domain adaptation regularization term to the ELBO when training [79]. A scaling factor on this regularization term decays from one to zero early in training.

### 4.3 Posterior predictive distributions linked to downstream tasks

For tasks like differential expression, denoising, and finding correlations, totalVI estimates functionals of posterior predictive distributions [22]. Define *C_n_* = {*x_n_*, *y_n_*, *s_n_*} as the set of observed data for cell *n*. First, consider the connection between the posterior predictive distribution of RNA data to totalVI denoised RNA expression. The posterior predictive RNA expression 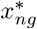 for gene *g* given *C_n_* is distributed following:

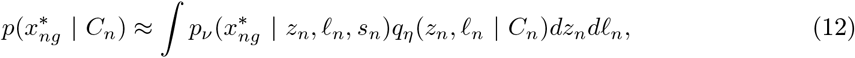

To produce denoised RNA expression, we compute the posterior predictive mean of 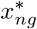. To further control for variation due to *ℓ_n_*, we condition on *ℓ_n_* = 1. By the law of total expectation,

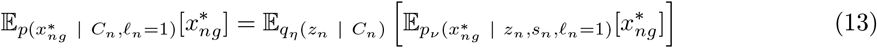

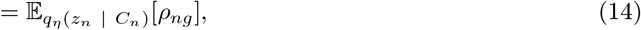

where *ρ_ng_* is the expectation of the RNA likelihood with the additional condition that *ℓ_n_* = 1.

For each cell *n*, we can compute the denoised RNA expression by averaging samples of *ρ_n_* generated by the following process:

1. Sample *z_n_* from *q_η_*(*z_n_* | *C_n_*)
2. Set *ρ_n_* = *f_ρ_*(*z_n_*, *s_n_*)

There are two important considerations for these posterior predictive distributions. First, we use the approximate posterior as a surrogate for the posterior. Second, these posterior predictive distributions are not tractable to compute in closed form, so we can only sample from them with ancestral sampling. Functionals of the posterior are computed using Monte Carlo integration.

#### Denoised protein expression

After training the model, we can generate “denoised” protein expression – protein expression effectively absent of background and controlled for sampling noise. Consider the perturbed protein generative process in which we set the background intensity to zero:

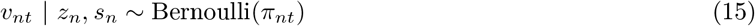

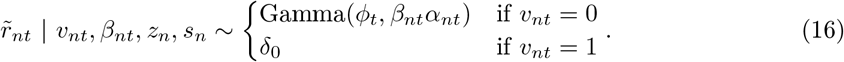

Here *δ*_0_ is a point mass distribution at 0. After marginalizing out *v_nt_*, 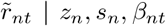 follows a zero-inflated Gamma distribution with mean (1 − *π_nt_*)*β_nt_α_nt_*.

For denoising, we return the posterior predictive mean of 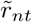. Indeed, the posterior predictive mean is equal to (1 − *π_nt_*)*β_nt_α_nt_* averaged over many posterior samples of *q*(*β_nt_*, *z_n_* | *C_n_*). In other words, we return the foreground mean, weighted by the probability that the observation was derived from the foreground. This can also be stated as subtracting the expected background from the expected total signal.

#### Missing protein imputation

To impute protein expression 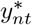 for cell *n* and protein *t* missing in batch *s_n_*, but that is observed in a batch *s*′ ≠ *s_n_*, do the following:

1. Sample *z_n_* from *q_η_*(*z_n_* | *C_n_*)
2. Sample *β_nt_* from *q_η_*(*β_nt_* | *z_n_*, *s* = *s*′)
3. Sample 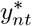 from 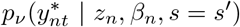

This process returns samples of 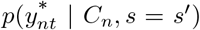. Intuitively, we encode the cell into the latent space, which is designed to mix the batches (i.e., be an integrated low-dimensional representation of the data), and obtain the parameters for the protein likelihood (decode) conditioned on the cell being in batch *s* = *s*′. Thus, the quality of imputation relies on how well batches mix in the totalVI latent space. Ultimately, we report the expected value of the imputed distribution

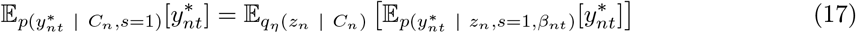

We may also impute the denoised expression, by exchanging 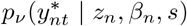 with 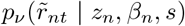. This change would additionally remove the protein background contribution to the prediction.

#### Differential expression

With a single model fit, totalVI can detect differentially expressed features between sets of cells, i.e., the model does not need to be retrained for every test. Here we use the Bayesian framework of [47] to detect differential expression (DE) of genes and proteins. Let

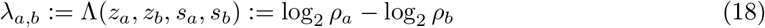

be the log fold change (LFC) of RNA expression between cells *a* and *b*. Then the probability that gene *g* is differentially expressed (DE) is

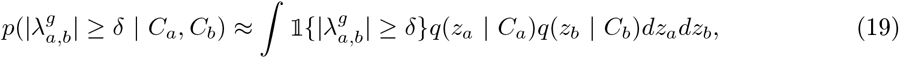

where *δ* is a threshold for the effect size. Intuitively, we are measuring the fraction of posterior samples that the absolute LFC greater than or equal to *δ*. For all experiments we set *δ* = 0.2. We compare the DE probability to the probability that the LFC is in the null region 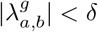 using a Bayes factor:

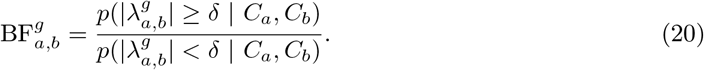

This can also be extended to groups of cells. Let *A* = {*a*_1_,*a*_2_,…, *a_m_*} be the indices of one subpopulation of interest, and *B* = {*b*_1_, *b*_2_,…, *b_n_*} be the other subpopulation of interest. We then exchange the posterior distributions in Equation 19 with the aggregated posterior:

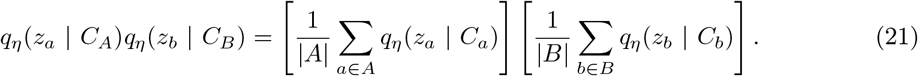

In this sampling procedure, a cell representation *z_a_* (resp. *z_b_*) is sampled given one randomly chosen cell in subpopulation *A* (resp. subpopulation *B*). Then, it is determined if 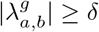 via an indicator function. The DE probability is estimated based on many samples.

Furthermore, by integrating over the batch variable *s_n_*, we effectively compare cells as if they were in the same batch [13]. For genes, this is equivalent to computing

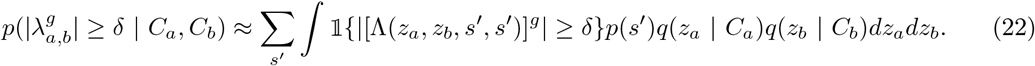

Here *p*(*s*′) is a uniform prior over batches. Every time we sample from the posterior, we decode the samples using the same batch indicator, averaging the DE probability over every possible batch indicator.

For proteins, we use the same framework, but define

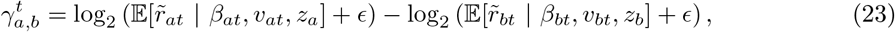

where the conditional expectation is equal to

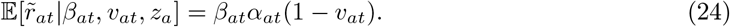

This is interpreted as the foreground mean if the cell was generated from the foreground, and zero otherwise. The added constant *E* is a “prior count” that helps define the log fold change when 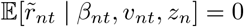. For all analysis, we set *ɛ* = 0.5. As with genes, we are interested in calculating 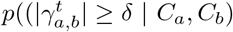, where in this case we integrate with respect to the distribution

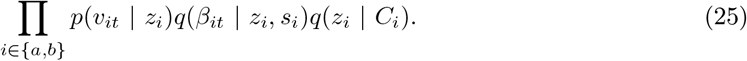

We consider features with a log(BF) > 0.7 as differentially expressed. This is roughly equivalent to calling features significant if the odds ratio (here equivalent to a Bayes factor) is greater than 2. Finally, we use the posterior samples of *λ_a,b_* (resp. *γ_a,b_* for proteins) as the estimate of effect size for each gene (resp. protein). Specifically, we use the median of the samples, which is robust to outliers and is also the Bayes estimator under *L*_1_ loss.

#### Denoised correlation matrix construction

We seek a feature-feature correlation matrix (e.g., gene-gene correlations, gene-protein cross-correlations) that summarizes biological variation, instead of technical variation. As totalVI explicitly models nuisance factors (for genes as well as proteins), we can query the model while controlling for this nuisance variation. Furthermore, because naive computations of correlations on denoised values (parameters of conditional distributions) were shown to induce spurious gene-gene correlations [37], we develop a novel sampling scheme that helps remove technical variation while avoiding such artifacts.

In order to ensure our correlation matrix does not include variation from the modeled technical factors, we perturb the data generating process to fix the library size (*ℓ_n_* = 10000) as well as incorporate the denoised protein expression conditional distribution. In particular, we compute a correlation matrix using samples from the distribution

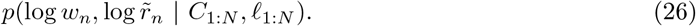

This is also a posterior predictive density whose samples are generated with ancestral sampling. As 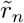 is zero-inflated, we add the same “prior count” before taking the logarithm. For this distribution, we sample ancestrally using the aggregated posterior

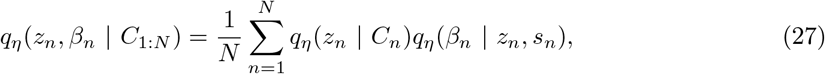

One could in principle replace the aggregated posterior with the prior in case of analyzing dataset-wide correlations. However, this approach is more flexible as it can be applied to calculate the correlation matrix for a subpopulation *A* = {*a*_1_, *a*_2_,…,*a_m_*}, where *A* is the set of indices for the subpopulation, by conditioning the distribution on *x_A_* and *y_A_*.

The distinction between this procedure and those that induced spurious correlations is that the latter effectively estimates a correlation matrix using the expected value of the posterior predictive distribution, rather than estimating the correlation matrix of the posterior predictive distribution.

#### Out-of-batch generalization

totalVI learns a transformation from *z_n_* and *s_n_* to the parameters of the conditional distributions for each feature (decoder). In an out-of-batch prediction, we predict the expression of a cell (e.g., the mean of conditional distribution) given a batch *s* ≠ *s_n_*. Here we describe a general way to sample posterior quantities for a cell while also “transforming” it into a different batch that was also observed for other cells. Special cases of this have already been described in the protein imputation and differential expression sections. Consider, for instance, the RNA counts in cell *n* and gene *g*. We can calculate posterior predictive samples of *x_ng_* while conditioning on any arbitrary observed batch *b*. Then,

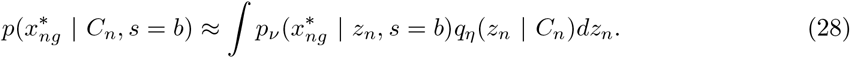

Furthermore, we can integrate over the choice of batch by sampling from

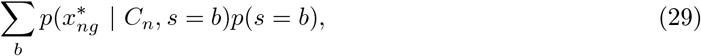

where *p*(*s*) is a uniform prior over batches. We take the expected value of this particular distribution as batch-corrected, denoised gene expression data. This “transforming” can also be applied to other likelihood parameters like *π_n_*.

### 4.4 CITE-seq experiment on mouse spleen and lymph node

Table 1 shows a summary of the experimental design that generated the mouse spleen and lymph node CITE-seq datasets. Below, we describe in further detail how these datasets were collected and processed.

#### Cell preparation

Two female C57BL/6 (B6) mice at 5 weeks of age were euthanized using CO2. From each mouse, six lymph nodes were harvested, pooled in RPMI + 10% FBS media on ice, mechanically dissociated with a syringe plunger, and passed through a 70 *μ*m strainer to generate a single cell suspension. Likewise, the spleen was harvested, placed in RPMI + 10% FBS media on ice, mechanically dissociated with a syringe plunger, and passed through a 70 *μ*m strainer to generate a single cell suspension. For the spleen, red blood cells were lysed in Red Blood Cell Lysis Buffer (BioLegend #420302) following the manufacturer’s protocol. All animal care and procedures were carried out in accordance with guidelines approved by the Institutional Animal Care and Use Committee at BioLegend, Inc.

#### Antibody panel preparation

We prepared panels containing 111 and 208 antibodies from BioLegend in the TotalSeq-A format (Supplementary Data: Antibodies). We performed a buffer exchange on each panel using a 50kDa Amicon spin column (Millipore #UFC505096) following the manufacturer’s protocol to transfer antibodies into RPMI + 10% FBS. Spleen and lymph node cell suspensions were stained with different hashtag antibodies [28].

#### CITE-seq protocol and library preparation

The CITE-seq experiment was performed following the TotalSeq protocol with two slight modifications. First, the 10 minute centrifugation at 14,000g to remove antibody aggregates was conducted prior to buffer exchange. Second, cells were stained, washed, and resuspended in RPMI + 10% FBS to maintain viability. After staining, washing, and counting, 12,000 spleen cells and 12,000 lymph node cells were mixed and loaded into a single 10x lane. We followed the 10x Genomics Chromium Single Cell 3’ v3 protocol to prepare RNA, antibody-derived-tag (ADT) and hashtag-oligo (HTO) libraries [80].

#### Sequencing and data processing

RNA, ADT, and HTO libraries were sequenced with an Illumina NovaSeq S1. Reads were processed with Cell Ranger v3.1.0 with feature barcoding, where RNA reads were mapped to the mouse mm10-2.1.0 reference (10x Genomics, STAR aligner [81]) and antibody reads were mapped to known barcodes (Table 2). Hashtags were demultiplexed separately for each 10x lane with HTODemux in Seurat v3 using the kmeans function [38]. No read depth normalization was applied when aggregating datasets.

### 4.5 Additional datasets

We also used publicly available CITE-seq datasets from 10x Genomics. These included “10k PBMCs from a Healthy Donor - Gene Expression and Cell Surface Protein” (PBMC10k, [31]), “5k Peripheral blood mononuclear cells (PBMCs) from a healthy donor with cell surface proteins (v3 chemistry)” (PBMC5k, [45]), and “10k Cells from a MALT Tumor - Gene Expression and Cell Surface Protein” (MALT, [32]).

### 4.6 CITE-seq data pre-processing

For each dataset, after initial cell and gene filtering, we retained at least the top 4,000 highly variable genes (HVGs) as defined by the Seurat v3 method, merging HVGs from different batches when appropriate [38]. Dataset specific filtering is described below.

#### Spleen and lymph node

An initial cell filter removed cells expressing fewer than 200 genes. Cells labeled as either doublets or negative for hashtag antibodies by HTODemux were also removed. A protein library size filter retained cells with between 400 and 10,000 total protein UMI counts. We also filtered on the number of proteins detected. For cells stained with the 111 antibody panel, we removed cells with fewer than 90 proteins detected, while the cutoff was set to 170 for cells stained with the 208 antibody panel. Cells with a high percentage of UMIs from mitochondrial genes (15% or more of the cell’s total UMI count) were removed. An initial gene filter removed genes expressed in 3 or fewer cells in any given batch. In addition to the top 4,000 HVGs selected by the Seurat v3 method, we retained genes that encode the proteins targeted by the 111 antibody panel. This resulted in 4,005 total genes. After all filters, the distribution of cells per dataset was: (SLN111-D1, 9,264 cells), (SLN111-D2, 7,564 cells), (SLN208-D1, 8,715 cells), (SLN208-D2, 7,105 cells). This is a total of 32,648 cells. Unless otherwise stated, we filtered out isotype control antibodies (9 total in the 208 panel) and hashtag antibodies. The protein CD49f was also removed due to having very low total UMI counts.

#### PBMC10k, PBMC5k, & MALT

For each of these datasets, we first removed doublets using DoubletDetection [82]. Cells with high mitochondrial content (percentage of UMIs from mitochondrial genes), high number of genes detected, high UMI counts, and with fewer than 200 genes expressed were removed. Next, cells with outlier protein library size (on either end) were removed. Genes with expression in three or fewer cells were removed. Finally, the top 4,000 HVGs were retained. Dataset specific parameters are in Table 3. In the case where the PBMC datasets are integrated, the 4,000 HVGs are selected by merging HVGs computed on each dataset separately as in the Seurat v3 method.

### 4.7 Posterior predictive checks and held-out metrics

Posterior predictive checks are useful to check the fit of Bayesian models. They work by comparing the observed data to posterior predictive samples from the model [29]. Much of the benchmarking done here was inspired by previous work done to benchmark the scHPF model [14]. We primarly compared totalVI to factor analysis, which is a linear-Gaussian alternative to totalVI, and is easily extendable to multiple modalities are features are treated conditionally independent of the latent representation. We also compared performance on RNA only to scVI [13]. Posterior predictive samples for totalVI and scVI were obtained by calling the `generate` function in the scVI package. We ran scVI with 20 latent dimensions and negative binomial conditional distribution in order to be consistent with totalVI. Factor analysis (FA) models were fit using the sklearn package [83] on the combined RNA and protein measurements using one of two normalization procedures. The first procedure consisted of transforming each value by log(count + 1). The second procedure consisted of log library size normalizing the RNA features and protein features separately. For example, considering only the RNA measurements for a cell, we normalized each cell to sum to 1 by dividing by the library size of RNA, multiplied by 10,000, added 1 to each value, and took a log transformation:

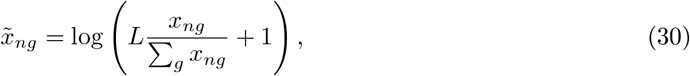

where *L* = 10000. This process was then applied to the protein measurements. We refer to this type of normalization as *log library size normalization*, and for short, *log rate*. These normalization procedures are necessary as FA assumes a Gaussian distribution, so training on the raw data would lead to poor model fit. Posterior predictive samples for FA models were computed using the fitted parameters and posterior distribution derived in [84]. We note that normalization procedures were inverted so that FA posterior predictive samples were on the same scale as the raw data.

For each dataset, each model was trained on a train set comprising of 85% of the cells. An additional 5% of cells were held-out as a validation set for totalVI early stopping. The remaining 10% of cells were also held-out as a test set. For each model’s posterior predictive samples (25 for each model) based on the train set, we calculated the coefficient of variation (CV) for each feature, and calculated the mean absolute error between the average CV and the observed raw data CV. We also used posterior predictive samples to assess generalization to unseen data. In this setup, we generated posterior predictive samples (150 for each model) conditioned on the test set. We considered the mean absolute error between the observed held-out data and the posterior predictive mean.

Moreover, we computed a held-out calibration error [30] for each model based on the test set. For each cell *n* and gene *g*, let *I_ng_* be the indicator that the observed value is contained in the interval of all posterior predictive samples. The calibration error for genes is then calculated as

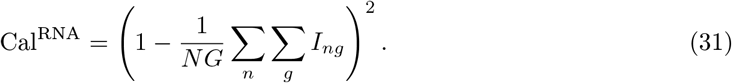

The calibration error for proteins is computed separately following the same procedure.

Finally, for totalVI and scVI only, and for only the RNA data, we computed the held-out predictive log likelihood. In this metric, *z_n_* and *ℓ_n_* were sampled from the variational posterior for each cell *n* and the average negative conditional log likelihood, −log *p*(*x_n_* | *z_n_*, *l_n_*, *s_n_*) was computed. This is also called the reconstruction loss in the VAE literature. This is also an approximation of −log *p*(*x_n_* | *x_n_*, *y_n_*, *s_n_*), the negative predictive log likelihood. We note that we cannot compare the log likelihood of totalVI and scVI, which use discrete conditional distributions to factor analysis models, which use continuous conditional distributions.

### 4.8 Background decoupling benchmarking

We reported the totalVI background probability as the posterior predictive mean of *π_nt_*, thus

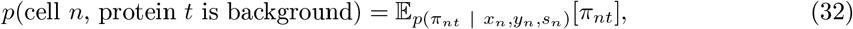

where the expectation is approximated using Monte Carlo integration. The totalVI foreground probability is one minus the background probability.

#### Observing protein background in empty droplets and non-expressing cell types

To observe different sources of protein background, we considered both empty droplets and cell types with known expression of surface markers. We defined empty droplets as non-cell barcodes from the SLN111-D1 dataset with between 20 and 100 RNA UMI counts (approximately 75,358 barcodes). We chose these criteria so that empty droplets were likely to represent ambient molecules rather than sequencing errors (with very low UMI counts) or cell debris (with higher UMI counts) [85]. To observe non-specific binding of antibodies, we considered B cells (which are known to express CD19 and CD20, but not CD28) and T cells (which are known to express CD28, but not CD19 or CD20). Using cell type annotations as described below (Methods 4.11), we grouped all high-quality, non-doublet B cell clusters (excluding plasma B cells), and alpha/beta T cell clusters (including all CD4, Treg, and CD8 T cell clusters). We observed that for these three proteins, both empty droplets and the non-expressing cell type contained protein background (non-zero protein counts) with varying degrees of overlap with the foreground signal of the expressing cell type. In this text, we describe the protein counts of the non-expressing cell type above the counts in empty droplets as non-specific antibody binding, although we acknowledge there could be multiple sources of this cell-specific background (Appendix A).

#### GMM cutoff for protein foreground/background

As a baseline determination of a cutoff to distinguish cells with foreground or background protein expression, we used a Gaussian mixture model (GMM). We applied scikit-learn’s `GaussianMixture` with default parameters to fit a GMM with two components to the log(protein counts + 1) for each protein for all cells in the SLN111-D1 dataset. We interpreted the posterior probability of the component with the higher mean as the foreground probability and that of the lower mean as the background probability. The GMM cutoff between foreground and background was determined to be the protein expression level at which the foreground probability was closest to 0.5.

#### Classification of cell type by marker proteins

We sought to evaluate totalVI against a GMM at predicting major cell types by the foreground probability of commonly used surface markers. Restricting all cells to just those that fell into the categories of B cells or T cells as described above, we tested how well totalVI or a GMM could classify cell types based on commonly used protein surface markers. We used a GMM fit on all cells of the SLN111-D1 dataset for each protein as described above. For each protein, we computed a receiver operating characteristic curve (ROC) (`sklearn.metrics`) by thresholding the totalVI or GMM foreground probability estimates (described above), using cell type annotations as true labels. We reported the area under the ROC (ROC AUC). The cell type considered as the positive population was either B cells, T cells, CD4 T cells, or CD8 T cells depending on the marker. In tests considering each of these positive populations, all remaining cells among the B and T cells were considered the negative population. Marker proteins tested included, for B cells: CD19, CD45R-B220, CD20, I-A-I-E (MHC II); for T cells: CD5, TCRb, CD28, CD90.2; for CD4 T cells: CD4; for CD8T cells: CD8a, CD8b [86–89]. Although we are aware of documented exceptions to these markers appearing strictly on a single cell type (e.g. CD5 is expressed on a portion of B1 B cells), these exceptions are rare. In these cases where marker expression is not mutually exclusive, cell types can still be distinguished by the gradation in levels of the marker between cell populations. Thus, these exceptions do not negate the utility of these markers in broad cell type classification (which is apparent in both totalVI and GMM performance at this classification task).

#### Visualization and raw data normalization

For the SLN111-D1 dataset, we visualized the totalVI latent space in two dimensions using Scanpy’s [70] implementation of the UMAP algorithm [44]. We applied log library-size normalization to the raw RNA counts as in Equation 30.

#### Distinguishing foreground and background in trimodal protein distributions

Despite using a two-component mixture, totalVI can decouple the background and foreground of proteins that have more than two modes globally. totalVI is capable of distinguishing foreground and background in this setting because the mixture is conditionally dependent on *z_n_*, which allows the foreground and background expression modes to be defined locally in the latent space. For example, as has been reported using flow cytometry [90], CITE-seq data of peripheral blood mononuclear cells contains three modes of CD4 expression corresponding to CD4 T cells, monocytes, and background. totalVI detected that both CD4 T cells and monocytes had foreground expression of CD4, while the CD4 expression of the remaining cells was attributable to background expression.

#### Denoised protein expression

Denoised protein expression was calculated as described in Methods 4.3. B cells and T cells were defined by annotations, as described above.

### 4.9 RNA-protein correlation analysis

#### Evaluation of correlation calculation with permuted features

Using totalVI, we aimed to calculate a correlation matrix between all RNA and protein features free from nuisance variation such as sequencing depth and protein background. We took care to avoid the naive calculation of correlations directly between denoised features, noting that a recent study reported false positive correlations in smoothed scRNA-seq data [37]. Instead, we developed a novel sampling method for the calculation of denoised feature correlations that removes nuisance variation while avoiding imputation-induced artifacts (Methods 4.3).

To evaluate whether totalVI could calculate a denoised feature correlation matrix without introducing spurious relationships in the data, we permuted the expression of a set of genes to serve as a negative control. To create this set of negative control genes from the SLN111-D1 dataset, we selected the 100 genes with highest mean expression that were not already among the top highly variable genes used in the model (Methods 4.6). We randomly permuted the counts of these genes within each cell, rendering these genes independent of all other gene and protein features. After concatenating the SLN111-D1 dataset with the permuted gene expression for all cells, we ran the totalVI model.

We then calculated Pearson and Spearman correlations between features using three methods, referred to here as raw, naive totalVI, and totalVI correlations. Raw correlations were calculated between log library-size normalized RNA (Equation 30) and log(protein counts + 1). Naive totalVI correlations were calculated between totalVI denoised RNA (Methods 4.3) and totalVI denoised proteins (Methods 4.3). totalVI correlations were calculated by sampling denoised RNA and denoised protein values from the posterior (Methods 4.3).

We observed the correlations between all RNA and protein features as well as the 100 additional genes whose expression was randomly permuted. By comparing the raw correlations with denoised correlations, we observed whether the method of denoising could maintain the relationship between these permuted genes and other features, which, in expectation, are independent from each other. Here, we highlighted the correlations between all proteins and the randomly permuted genes, whose correlations are expected to be near zero.

#### Correlations of RNA-protein pairs

We calculated a feature correlation matrix for the SLN111-D1 dataset using either the totalVI sampling method or by calculating raw correlations as described above. The resulting feature correlation matrices for both Pearson and Spearman correlations were subset to each protein and its encoding RNA for all proteins with a unique encoding RNA in the dataset (i.e., excluding RNA with multiple isoforms such as Ptprc).

### 4.10 Integration of multiple datasets

We compared totalVI’s integration performance to that of Scanorama [39] and Seurat v3 [38]. We chose these methods because like totalVI, they provide both batch-corrected measurements and a low-dimensional integrated latent space. The input to both Scanorama (`scanorama.correct`) and Seurat v3 (`FindIntegrationAnchors, IntegrateData`) methods was a normalized matrix of concatenated genes and proteins. Genes were subset to the same subset used as input to totalVI. The RNA counts of this matrix were normalized following standard log library size normalization (Equation 30). For proteins, we used a *y* → log(*y* +1) transformation. Finally, we standard scaled each feature. Scanorama and Seurat v3 were run with default parameters. We compared the performance of totalVI, Scanorama, and Seurat v3 using the following metrics:

#### Latent mixing metric

The latent mixing metric measures how well the latent cell representations are mixed between batches relative to the global frequency of batches. First, a cell-cell similarity matrix is computed from a latent representation of cells. Next, select 100 cells uniformly at random, and calculate the frequency of batches represented in each cell’s 100 nearest neighbors. Let 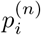 be the frequency of batch *i* in the 100 nearest neighbors of cell *n*. Let *q_i_* be the global frequency of batch *i*. Compute the negative relative entropy between the frequency of observed batches in the neighborhood, and the global frequency of batches:

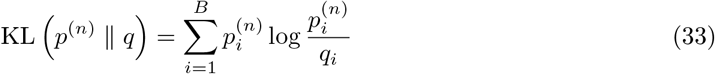

Repeat this 50 times and return the average negative relative entropy.

#### Measurement mixing metric

The measurement mixing metric describes how well the high-dimensional measurements are batch corrected, and for each feature, is related to the Mann-Whitney U statistic. Consider one feature in the batch-corrected data matrix placed in rank order. Let *R*_1_ be the sum of the ranks of the cells in batch 1 and *N*_1_ be the number of cells in batch 1. Define 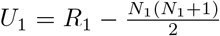. Similarly, compute *U*_2_ for batch 2 and return min(*U*_1_, *U*_2_). Higher values of this metric indicate better mixing within that feature.

#### Feature retention metric

The feature retention metric describes how spatial autocorrelation of both RNA and protein change when comparing cells from an integrated latent representation to a latent representation derived from each batch separately. Lower values of this metric indicate that the integration procedure reduced the localization of feature expression, indicating some degree of random mixing. We calculate it as follows. For two batches and a particular integration method, we calculate *Z*_1_ and *Z*_2_, the latent representations of the cells of batch 1 and batch 2, respectively. The latent space computation of the individual batches was chosen to closely match the integration method (see below). We also calculate an integrated latent representation of both batches 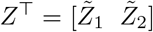. Let *D*_1_ = [*X*_1_ *Y*_1_] be the combined RNA and protein batch 1 in which RNA is library size log normalized and proteins are log-transformed. Let 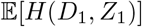 be the expected feature autocorrelation score as calculated by Hotspot [43]. Furthermore, let 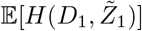 be the analogous quantity calculated using the latent cell representations of batch 1 subsetted from the joint, integrated representation. The feature retention metric is calculated as 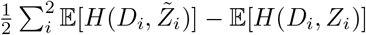. In the case of totalVI union, features were intersected to compute this metric.

For Scanorama, we define *Z*_1_ and *Z*_2_ to be a 100-dimensional matrix produced with principal components analysis (PCA), which is the same dimension reduction used in the integration method. For Seurat v3, we similarly use PCA to reduce *D*_1_ and *D*_2_ to 30 dimensions, the same number of dimensions used for integration. The input to PCA was the same as the input for the respective method, except for Scanorama, where we additionally *L*_2_ normalized each cell, because this step is done automatically by Scanorama’s `correct` method.

#### Missing protein imputation

For Seurat v3, we imputed proteins based on mutual nearests neighbors in the RNA data using the `FindTransferAnchors` and `TransferData` functions. Again, RNA data were log library size normalized. Proteins were not normalized as input to Seurat. For totalVI, after fitting the model, cells from the batch with held-out proteins were decoded conditioned on being in the batch with observed protein data. We used the Pearson correlation of values on the log scale to assess imputation accuracy. We note that CTP-net [91] is another method for imputing protein expression from scRNA-seq data; however, at the time of writing, the CTP-net package only offers a neural network that was pre-trained on specific CITE-seq datasets from human cells, with no option training with a new dataset from either mice or humans. Therefore, a direct comparison to totalVI and Seurat v3 is not straightforward.

### 4.11 Stratification of cells in SLN-all

We stratified cells of the mouse spleen and lymph node based on the SLN-all dataset (totalVI-intersect model fit as described above in Section 2.6). We clustered cells in the totalVI latent space with Scanpy’s implementation of the Leiden algorithm at resolution 1, resulting in 32 clusters [70, 92]. We repeated this approach to sub-cluster cells, finding a total of 43 clusters. We used Vision [61] with default parameters for data exploration, including its implementation of the Wilcoxon rank sum test, to identify cluster markers. Clusters were manually annotated based on a curated list of cell type markers (Table 4). Clusters annotated as doublets, low quality cells, or cells undergoing the cell cycle were removed from further analysis. Again, we visualized the totalVI latent space in two dimensions using Scanpy’s implementation of the UMAP algorithm.

### 4.12 Differential expression analysis

The t-test and Wilcoxon test for each differential expression scenario were run on protein features only using the Scanpy library, which produces adjusted *p*-values corrected by Benjamini-Hochberg procedure [116]. A protein was considered to be differentially expressed if the adjusted *p*-value was less than 0.05. Each application of totalVI differential expression tests to a dataset requires a trained totalVI model. For each dataset used in DE analysis, all cells were included to train the model. Throughout, we used our manual annotations from the SLN-all totalVI-intersect model run. The cells in nuisance clusters (described in previous section) were removed before running totalVI differential expression functions.

In a given totalVI differential expression test, we identified cell type markers by first filtering features for significance (log Bayes factor > 0.7), and then sorting by the median log fold change. We only retained genes with non-zero UMI counts in at least 10% of the subset of cells.

In the comparison to scVI gene Bayes factors, each method was trained independently on the SLN111-D1 dataset. We ran scVI with 20 latent dimensions and negative binomial conditional distribution to be consistent with totalVI. Differential expression of genes in scVI was computed using the same LFC-based method, which is implemented in the scvi package. In reproducibility benchmarking, totalVI was trained independently on the replicates. In the test between ICOS-high Tregs and CD4 conventional T cells, we used the same totalVI-intersect model fit that was used to manually annotate the cells.

#### DE on imputed proteins

In one totalVI model fit, SLN111-D1 and SLN111-D2 were integrated with the proteins of SLN111-D2 held out. In the second totalVI model fit, these two datasets were integrated with all data. In testing differential expression of proteins, and for each model fit, we conditioned on SLN111-D1. This is an application of Equation 22, except that the prior *p*(*s*′) is 1 if *s*′ = SLN111-D1 and 0 otherwise.

### 4.13 Archetypal analysis

This analysis was performed on the SLN-all intersect mode model run. As *z_n_* is distributed as logistic normal, the latent space is then constrained to the probability simplex (i.e., each *z_n_* is non-negative and sums to one). Archetypes correspond to vertices of the totalVI latent space, which means they can be represented by the identity matrix *I_d_*, where *d* is the number of latent dimensions (20 in all experiments). In this setting, the latent space is the 19-dimensional standard simplex. We first identified and removed four archetypes from further interpretation that suffered from inactivity (a known issue in training VAEs) [117]. For the remaining 16 latent dimensions, we decoded the archetypes to obtain denoised RNA and protein archetypal expression profiles, all conditioned on batch 0 (the SLN111-D1 experiment). We then computed denoised RNA and protein expression profiles for all cells in SLN-all, conditioned on SLN111-D1. To derive signatures for each archetype, we computed the mean and standard deviation of each feature in the denoised RNA and protein expression matrices (without the archetypes) and standard scaled the archetypal profiles with respect to this mean and standard deviation. We refer to this quantity as the archetype score. The top features for each archetype were those with an archetype score greater than 2. The distance to the archetype is computed as the Manhattan distance from each cell’s latent representation to the archetype. The distances per archeytpe were scaled into the range [0,1].

### 4.14 B cell analysis

For this analysis, we used the totalVI-intersect model fit on the SLN-all dataset as described above in Section 2.6. The SLN-all dataset was filtered to include all high-quality, non-doublet clusters annotated as B cells (excluding plasma B cells), resulting in 15,560 cells (Methods 4.11).

#### Calculation of signature scores

Gene signature analysis was conducted using Vision [61] with default parameters. Gene signatures, including interferon response signatures, were downloaded from MSigDB gene sets [118]. Signature scores were calculated on all cells in the SLN-all dataset based on cell similarities defined by the totalVI latent space.

#### Identification of transitional and mature B cell feature modules

totalVI Spearman correlations between all features were calculated separately within the transitional B cell cluster and the mature B cell cluster. Features were subset by the following method. From a one-vs-one DE test between transitional and mature B cells, we selected the top ten marker genes and top three marker proteins for each cluster (Methods 4.12). We added to this list the four features most highly correlated with each differentially expressed feature within its respective cluster. This resulted in a list of both transitional and mature features which we used to subset the full feature correlation matrix. Features were hierarchically clustered separately for transitional and mature B cells using Seaborn’s `clustermap` with default parameters.

When plotting totalVI expression of each feature as a function of 1 − *Z*_16_, each feature was standard scaled and smoothed with a loess curve. Spearman correlations were calculated between each feature and 1 − *Z*_16_. The p-values of these correlations were all significant (Benjamini-Hochberg adjusted *P* < 0.001).

## Code availability

The code to reproduce the experiments of this manuscript is available at https://github.com/YosefLab/totalVI_reproducibility. The reference implementation of totalVI is available via the scVI package at https://github.com/YosefLab/scVI.

## Data availability

Processed data are available in the reproducibility GitHub repository. Raw data are being uploaded to GEO. The SLN-all dataset can be explored interactively with Vision at http://s133.cs.berkeley.edu:9000/Results.html.

## 5 Supplementary Figures

**Figure S1:**
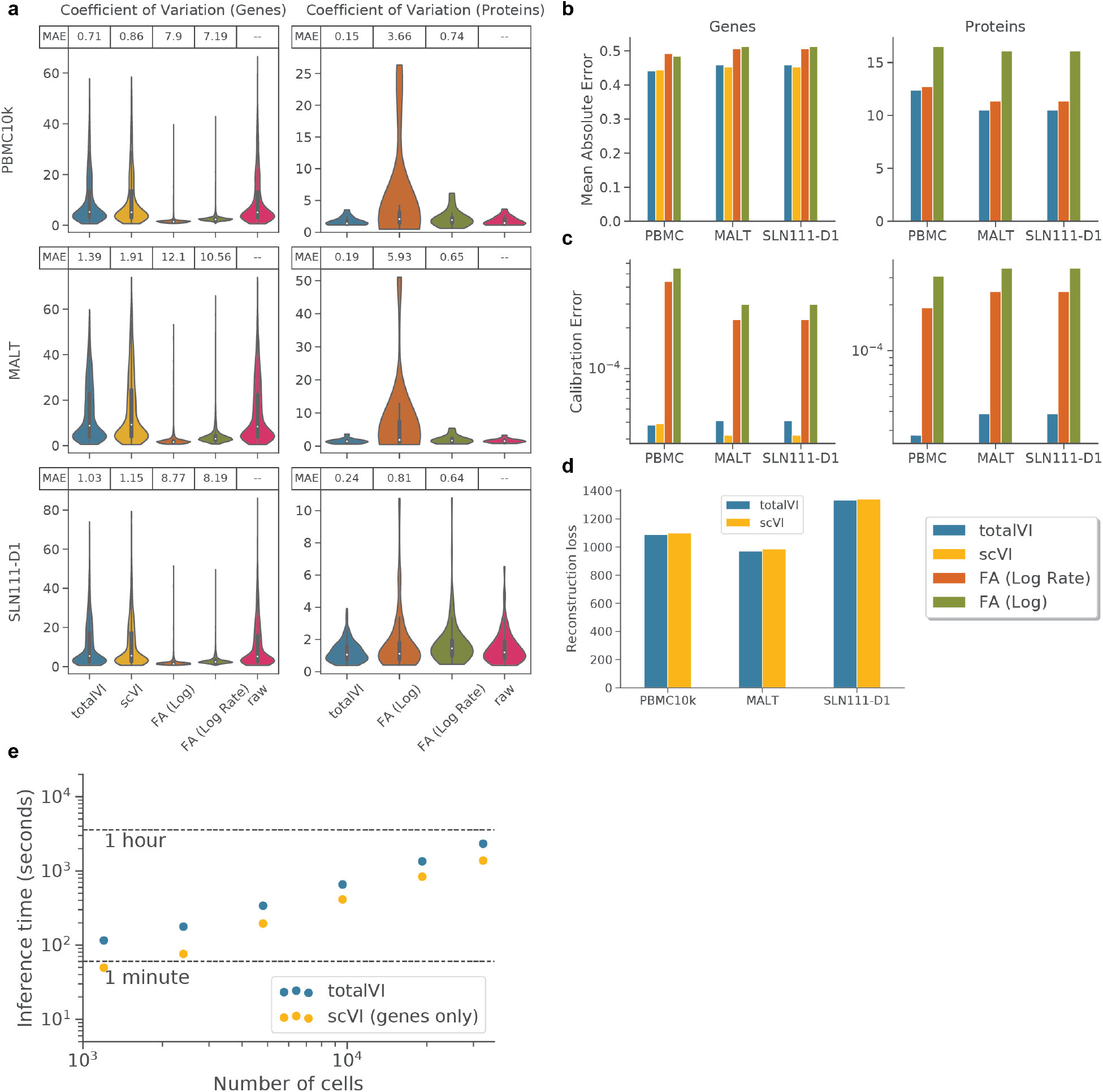
Evalutation of totalVI model. **a**, Posterior predictive check of coefficient of variation (CV) of genes and proteins. For each of the PBMC10k, MALT, and SLN111-D1 datasets and for each model (totalVI, scVI, factor analysis with normalized input) the average coefficient of variation from posterior predictive samples was computed for each feature. Violin plots summarize the distribution of CVs for genes and proteins. Mean absolute error (MAE) between raw data CVs and average posterior predictive CV are reported. **b**, MAE between held out data and posterior predictive mean separated by genes and proteins for each model and dataset. **c**, Calibration error of held-out data reported separately for genes and proteins. **d**, Held-out reconstruction loss of genes for scVI and totalVI. **e**, Inference time for totalVI and scVI across cells randomly subsampled to different levels from SLN-all. scVI was run with only genes.

**Figure S2:**
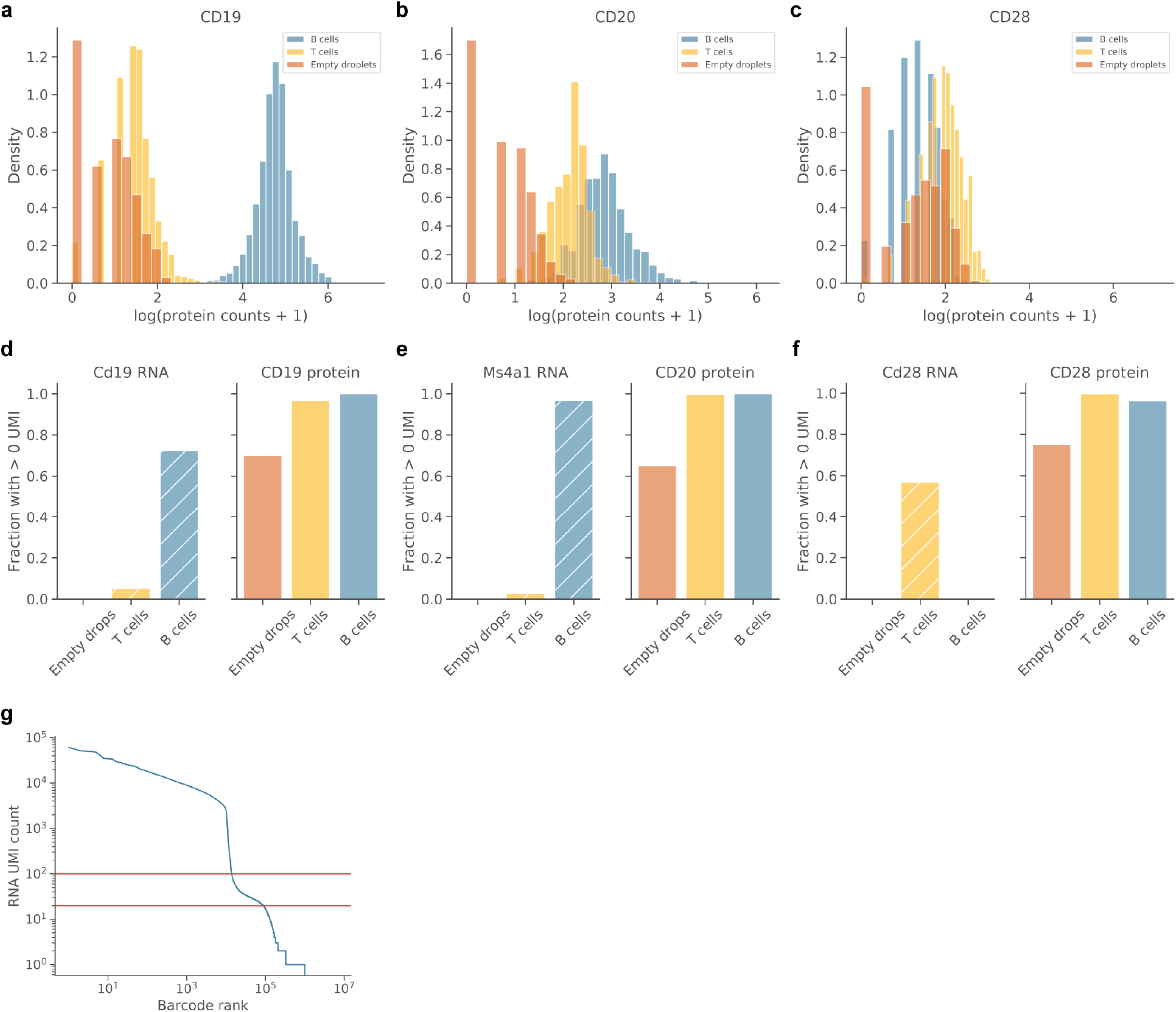
Protein background in cells and empty droplets. **a-c**, Histogram of log(protein counts +1) in the SLN111-D1 dataset for B cells, T cells, and empty droplets (Methods 4.8) for CD19 **(a)**, CD20 **(b)**, and CD28 **(c)**. **d-f**, Fraction of empty droplets, B cells, or T cells with > 0 UMIs detected for a given RNA (left, hatched) or protein (right, solid). RNA/proteins displayed are Cd19/CD19 **(d)**, Ms4a1/CD20 **(e)**, and Cd28/CD28 **(f)**. **g**, Barcode rank plot for all barcodes detected in the SLN111-D1 dataset. Red lines at 20 and 100 RNA UMI counts indicate the lower and upper bounds, respectively, used to define empty droplets in (a-f).

**Figure S3:**
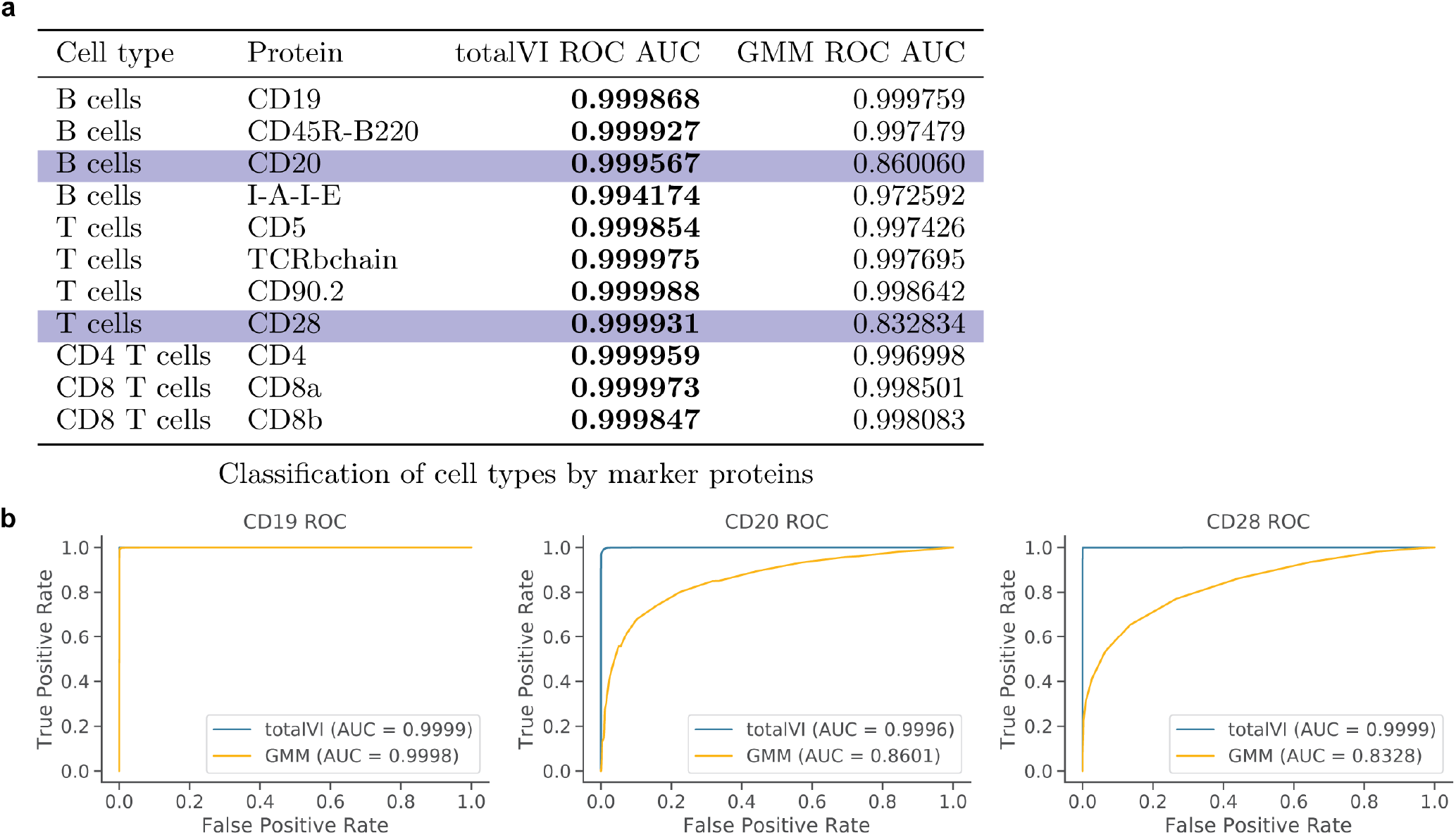
Classification of cell types by marker proteins. **a**, Performance of totalVI and a Gaussian mixture model (GMM) fit on all cells for each protein of the SLN111-D1 dataset. Area under the receiver operating characteristic curve (ROC AUC score) was calculated using as input either the totalVI foreground probability or GMM foreground probability where the indicated cell type was the positive population out of all B and T cells. Bolded ROC AUC scores indicate higher values (better performance). Highlighted in blue are two proteins for which totalVI noticeably outperformed the GMM. **b**, Receiver operating characteristic (ROC) curves for CD19 (B cells), CD20 (B cells), or CD28 (T cells).

**Figure S4:**
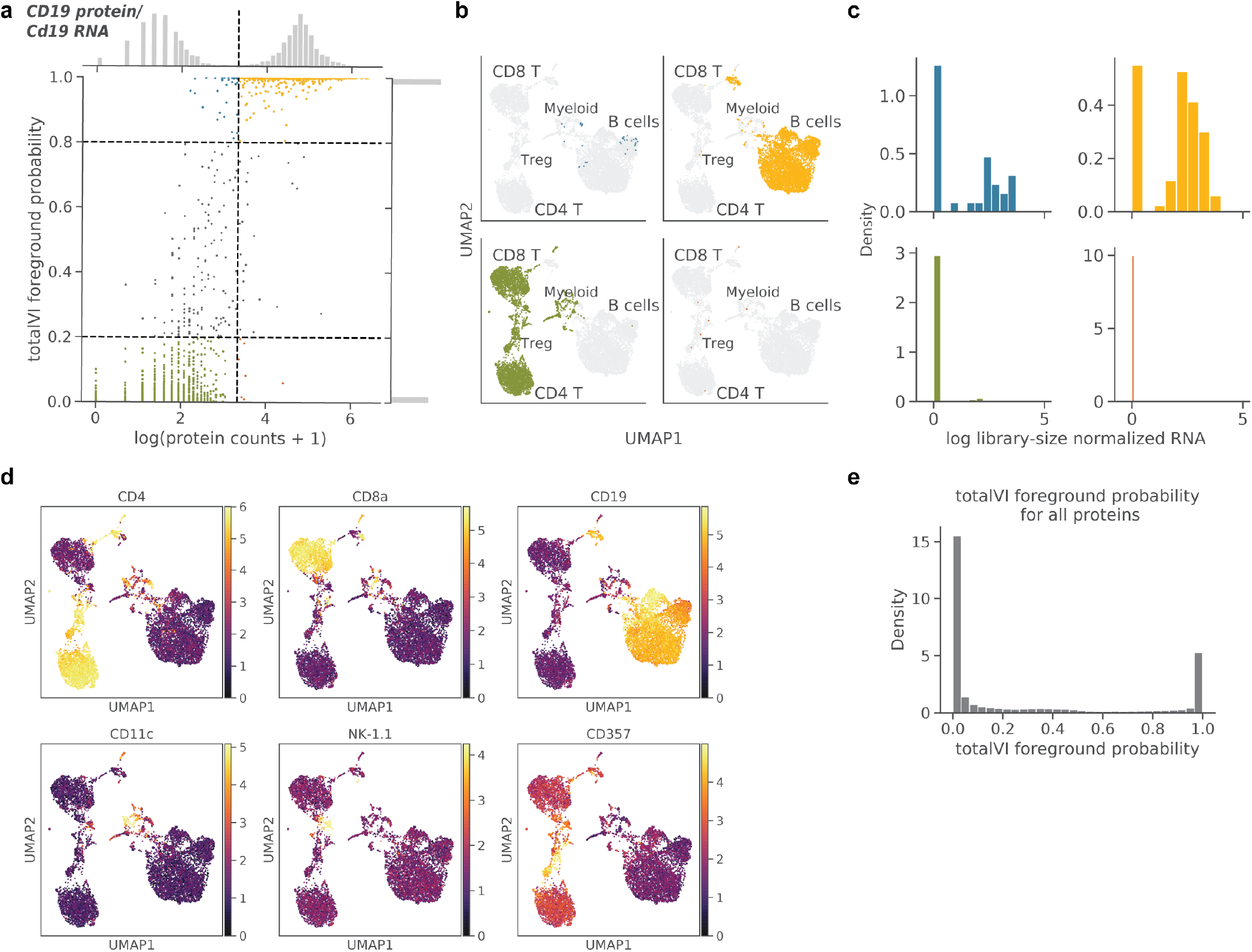
totalVI decouples protein foreground and background. totalVI was applied to the SLN111-D1 dataset. **a-c**, CD19 protein (encoded by Cd19 RNA). **(a)** totalVI foreground probability vs log(protein counts + 1). Vertical line denotes protein foreground/background cutoff determined by a GMM. Horizontal lines denote foreground probability of 0.2 and 0.8. Cells with foreground probability greater than 0.8 or less than 0.2 are colored by quadrant, while the remaining cells are gray. **(b)** UMAP plots of the totalVI latent space. Each quadrant contains cells from the corresponding quadrant of (a) in color with the remaining cells in gray. **(c)** RNA expression (log library-size normalized; Methods 4.8) for cells colored in (a). **d**, UMAP plots of the totalVI latent space colored by (log(counts + 1)) of cell type marker proteins (Table 4). **e**, totalVI foreground probability for all proteins across all cells in the SLN111-D1 dataset.

**Figure S5:**
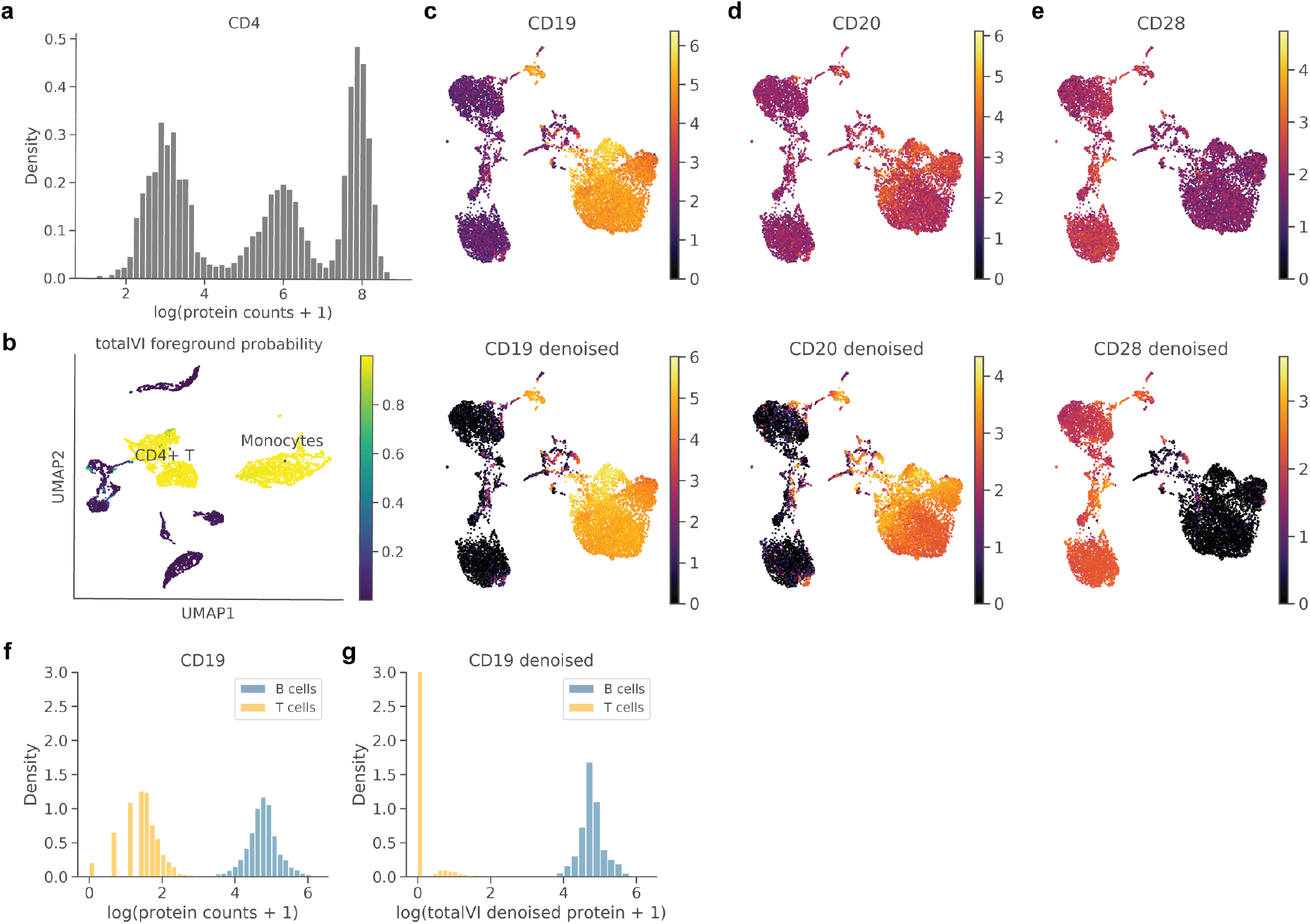
totalVI decouples foreground and background for trimodal protein distributions and denoises protein data. **a, b** CD4 protein expression in the PBMC10k dataset. **(a)** Trimodal distribution of log(protein counts + 1). **(b)** UMAP plot of the totalVI latent space colored by totalVI foreground probability. **c-e**, UMAP plots of the totalVI latent space for the SLN111-D1 dataset. Plots are colored by log(protein counts + 1) (top) and log(totalVI denoised protein + 1) (bottom) for CD19 **(c)**, CD20 **(d)**, and CD28 **(e)**. **f, g**, Distributions of log(protein counts + 1) **(f)** and log(totalVI denoised protein + 1) **(g)** for CD19 protein in B and T cells. y-axis is truncated at 3.

**Figure S6:**
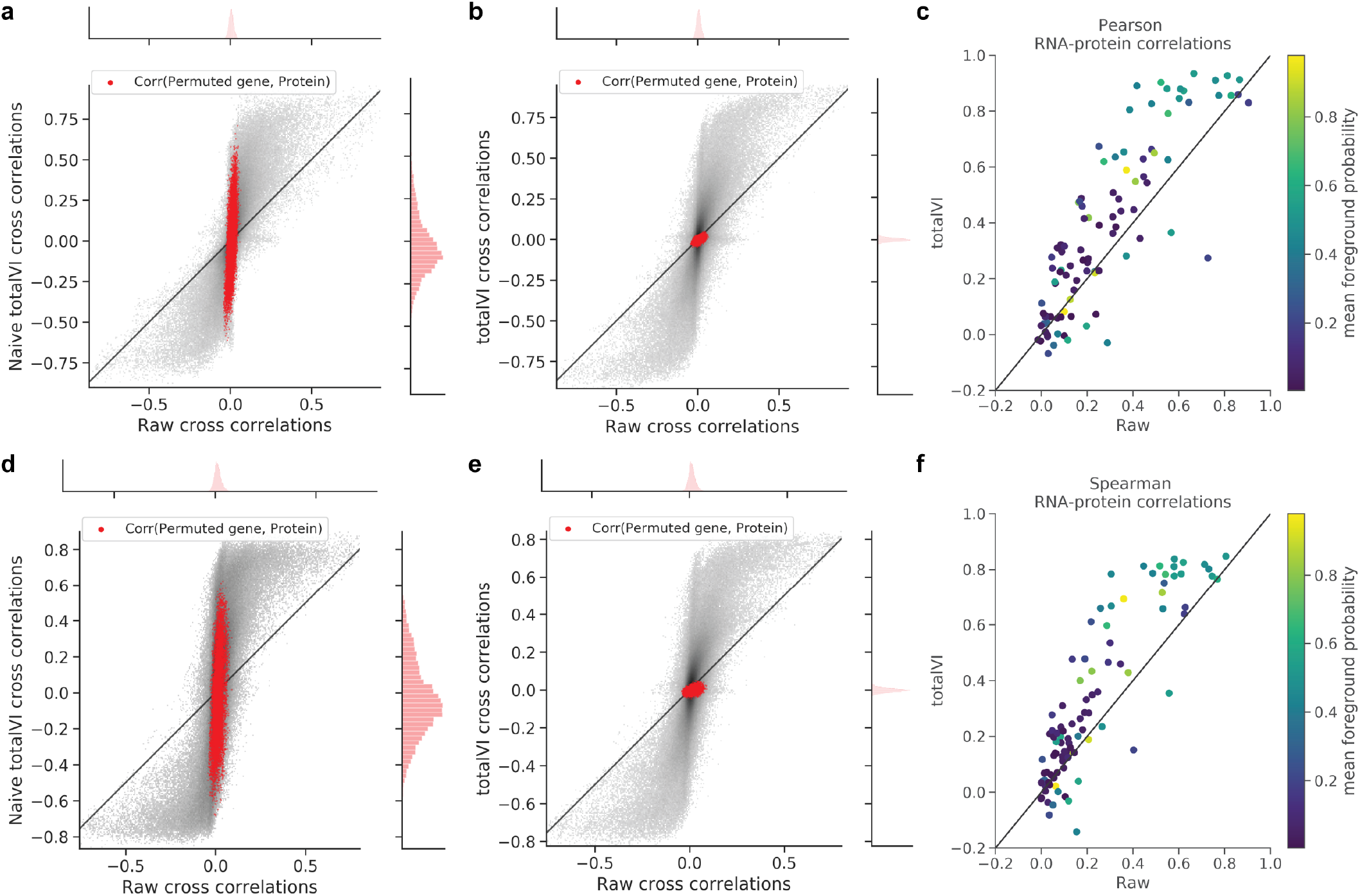
RNA-protein correlations. **a, b**, 2d density plots of Pearson correlations between all RNA and protein features in the SLN111-D1 dataset as well as 100 additional genes whose expression was randomly permuted. Correlations between all proteins and the randomly permuted genes are colored in red. Raw correlations were calculated between log library-size normalized RNA and log(protein counts + 1). **(a)**, Naive totalVI correlations were calculated between totalVI denoised RNA and totalVI denoised proteins. **(b)**, totalVI correlations were calculated between denoised RNA and proteins sampled from the posterior (Methods 4.3). **c**, Pearson correlations between each protein and its encoding RNA for all proteins with a unique encoding RNA, colored by the mean probability foreground of the protein across all cells. totalVI correlations were calculated as in (b) and raw correlation were calculated as in (a, b). **d-f**, Same as (a-c), but for Spearman correlations.

**Figure S7:**
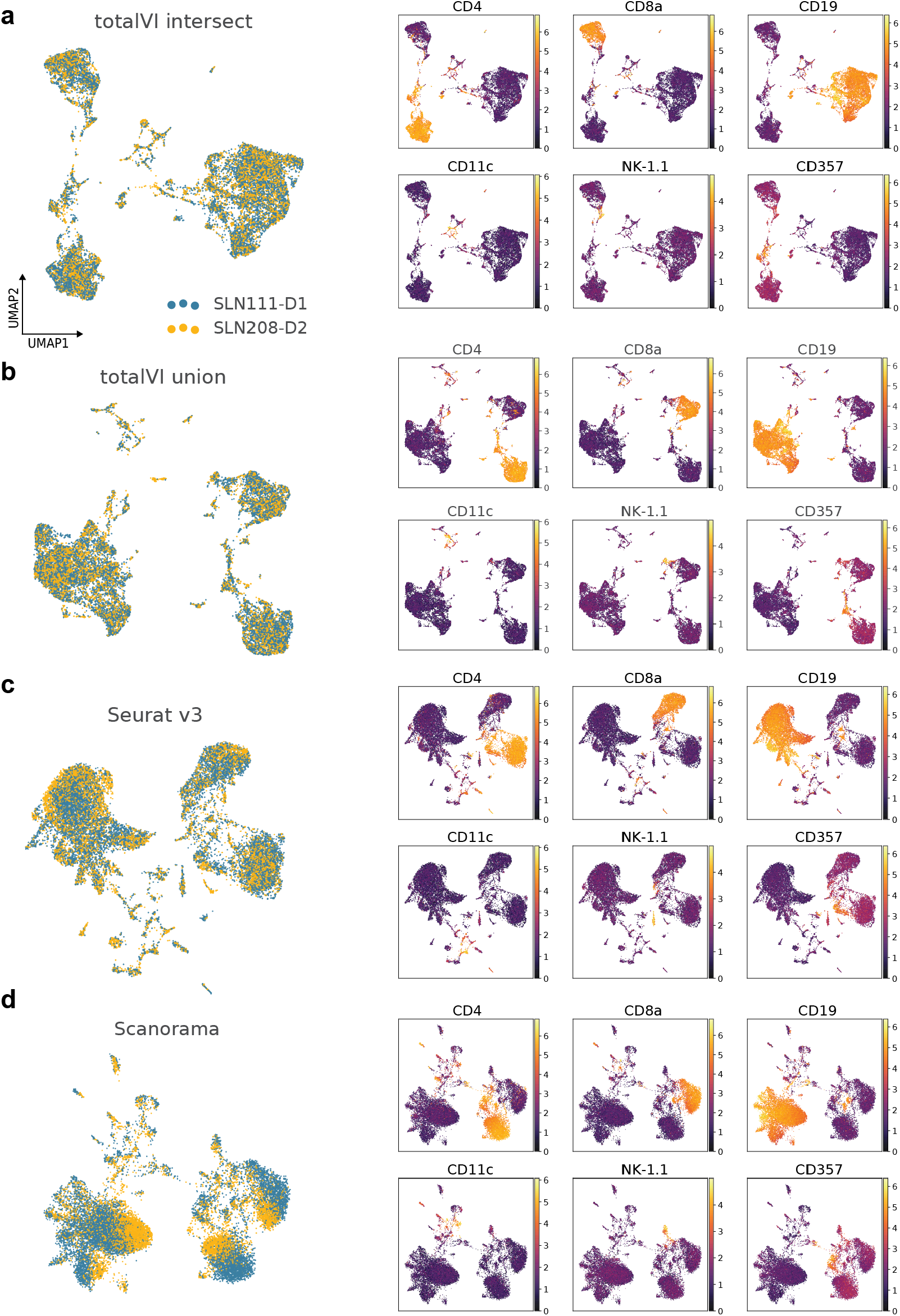
UMAP embeddings of integration methods on spleen and lymph node data. **a-d**, For each method, UMAP plots colored by dataset, and by log(counts + 1) of key marker proteins (Table 4).

**Figure S8:**
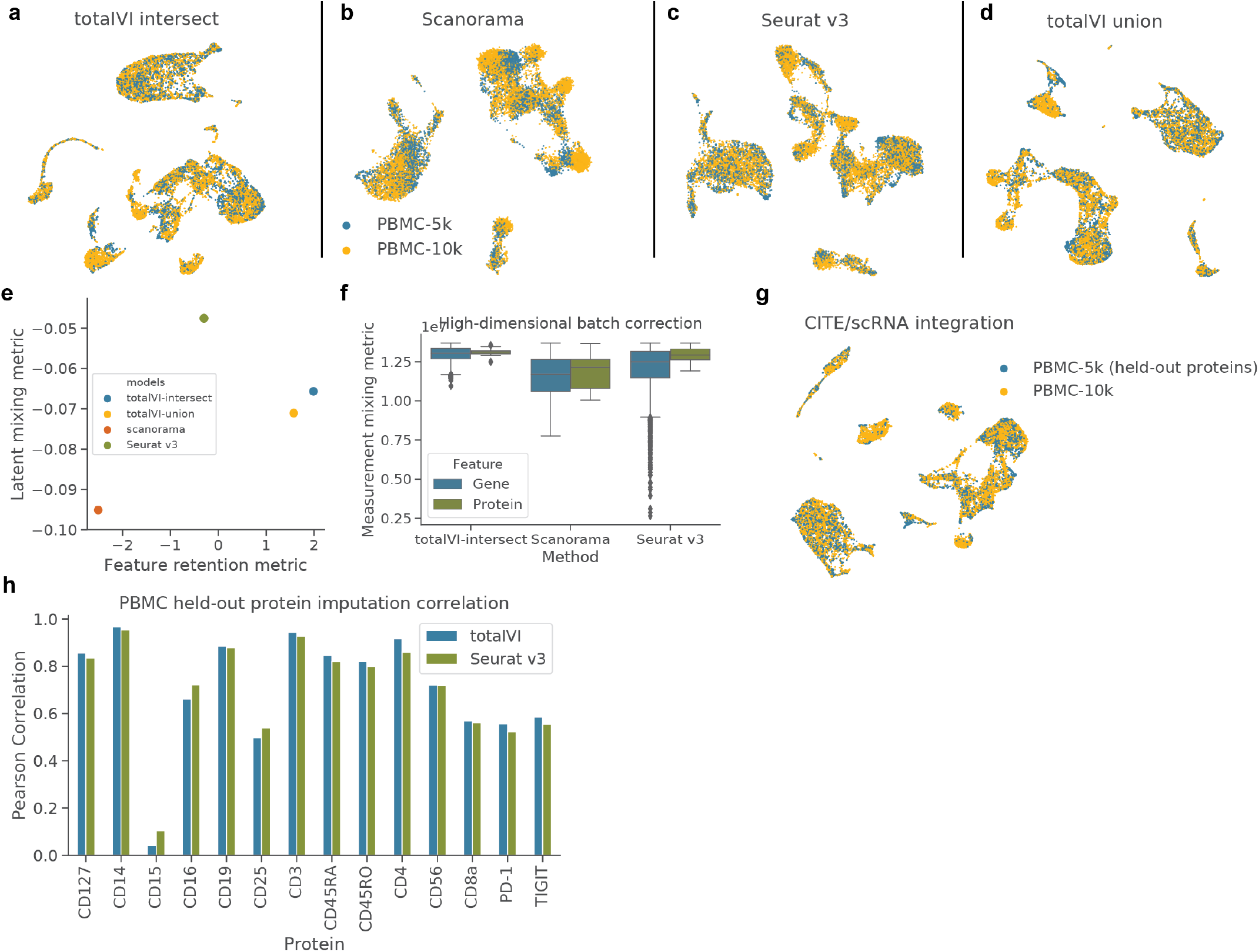
Benchmarking of integration methods on PBMC data. Integration methods were applied to PBMC10k and PBMC5k. **a-d**, UMAP plots of integrated latent spaces. **e**, Latent mixing metric and feature retention metric for each method. **f**, Measurement mixing metric applied indivdually to each batch-corrected feature. **g**, UMAP plot of integrated latent space from totalVI union mode when holding out the proteins from PBMC5k. **h**, Pearson correlation between imputed and observed proteins (log scale) from PBMC5k for totalVI and Seurat v3.

**Figure S9:**
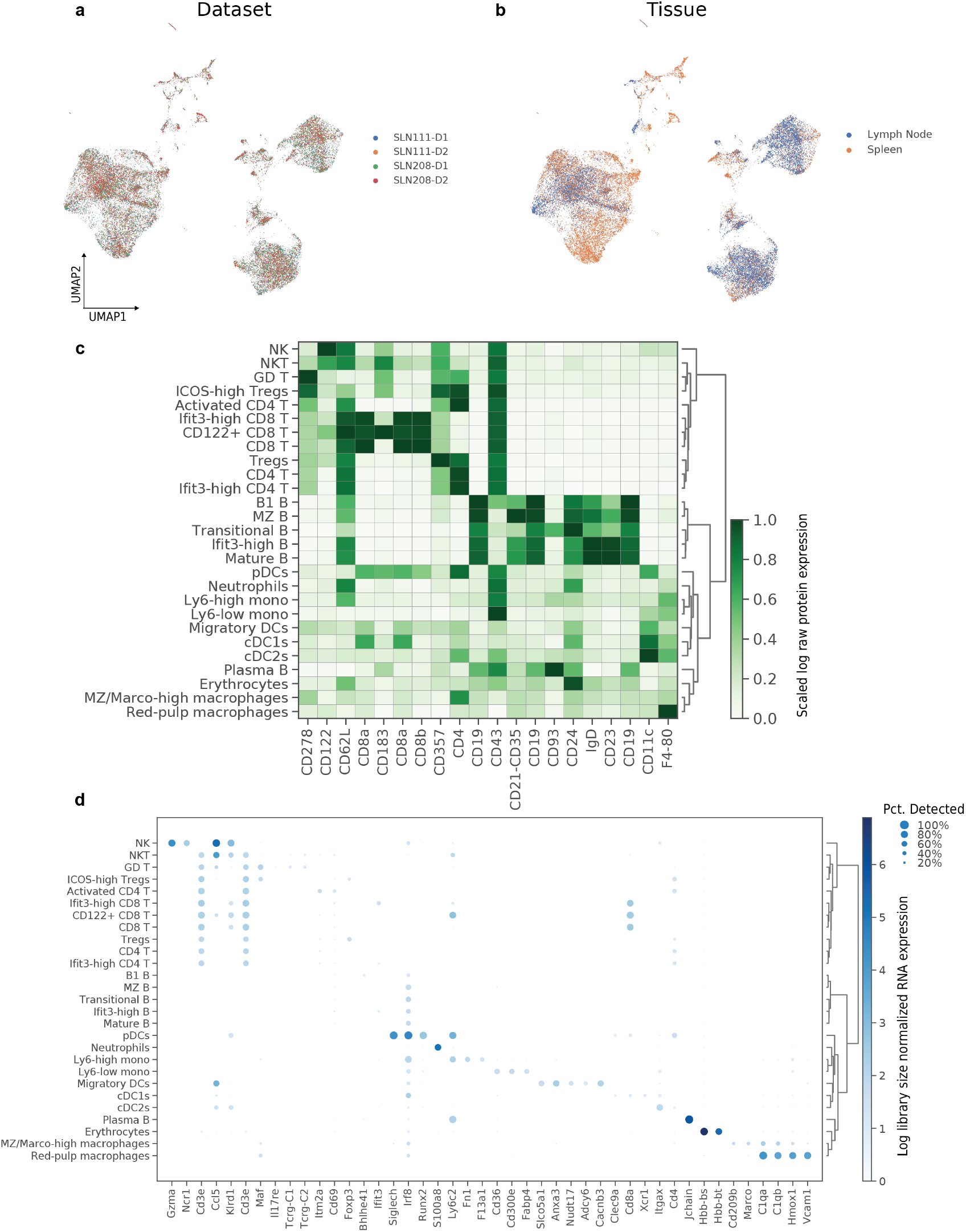
Integration of SLN-all with totalVI-intersect. **a, b**, UMAP plot of SLN-all colored by **(a)** dataset, and **(b)** tissue. **c**, Heatmap of proteins used for annotation. Proteins (columns) are log(protein counts + 1) scaled by column for visualization. **d**, Dotplot of RNA markers used for annotation. RNA is log library size normalized.

**Figure S10:**
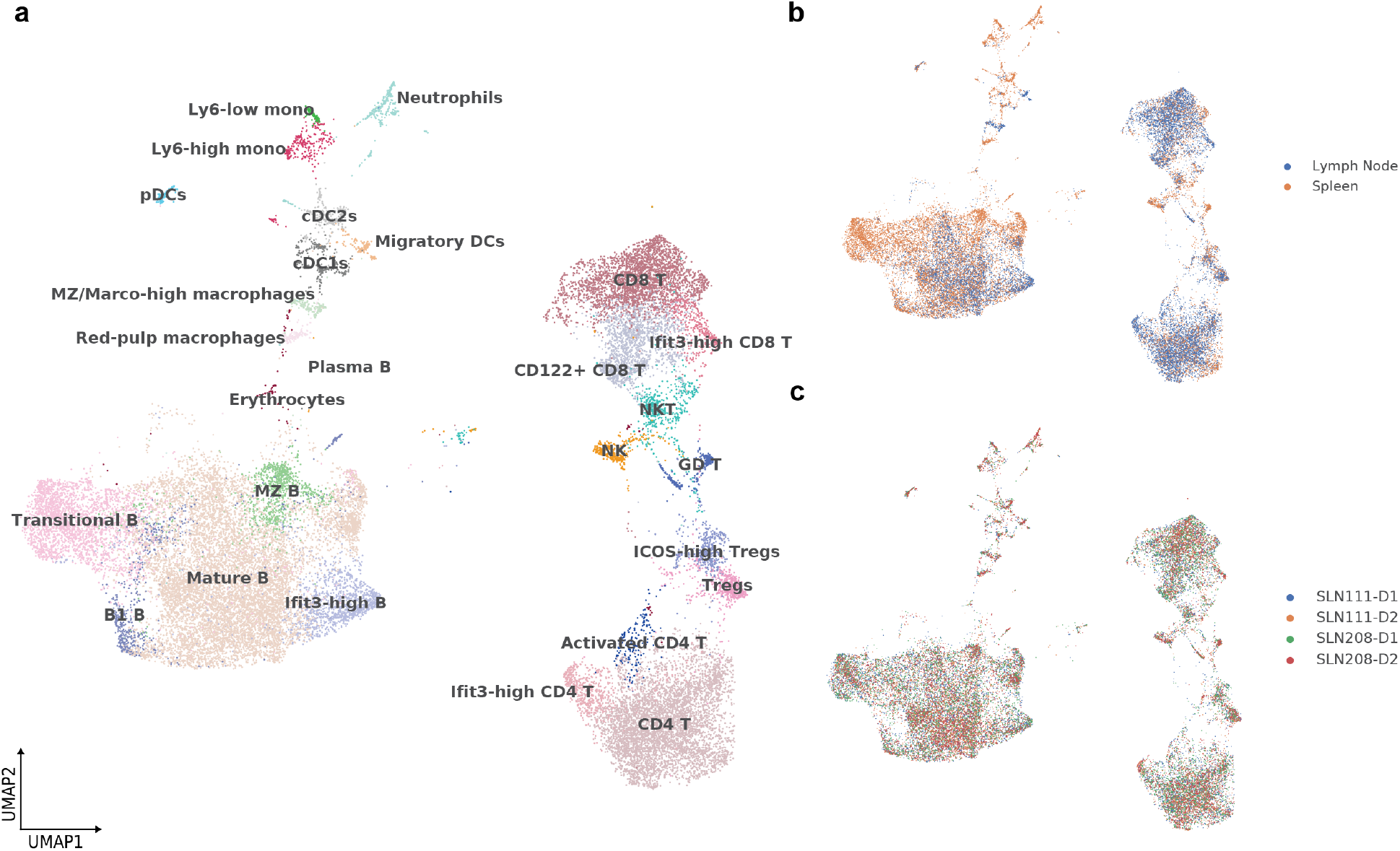
Integration of SLN-all with union of panels. **a-c**, UMAP plot of SLN-all colored by **(a)** cell types derived from manual annotation of model run with intersection of panels, **(b)** tissue, and **(c)**, dataset.

**Figure S11:**
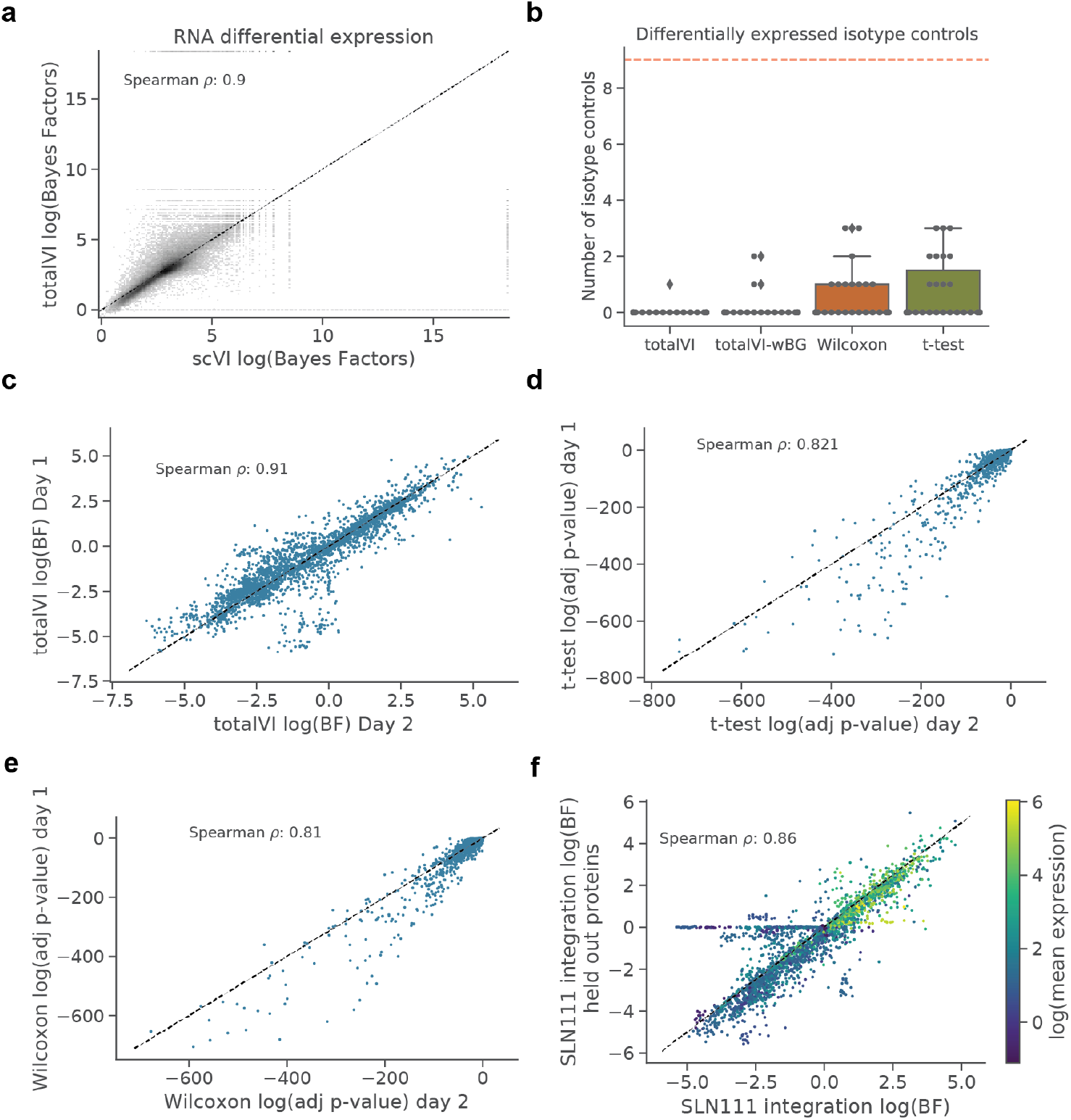
Differential expression analysis. **a**, 2d density plot of totalVI and scVI log Bayes factors for genes. Bayes factors were computed for each gene in one-vs-all tests on the SLN111-D1 dataset. **b**, Number of isotype controls called differentially expressed in one-vs-all tests for totalVI, totalVI-wBG (totalVI test without background removal), Wilcoxon rank-sum, and t-test. Tests were applied to SLN208-D1, for which isotype controls were retained. **c-e**, Significance level (Bayes factors for totalVI, adjusted *p*-value for frequentist tests) for proteins in one-vs-all tests computed on SLN111-D1 and SLN111-D2 for each of **(c)** totalVI, **(d)** t-test, **(e)** Wilcoxon. **f**, Bayes factors for proteins in one-vs-all tests computed on the SLN111 datasets integrated with and without the SLN111-D2 proteins held-out. Differential expression tests for both model fits were conditioned on SLN111-D1. Bayes factors are colored by the average protein expression from SLN111-D1.

**Figure S12:**
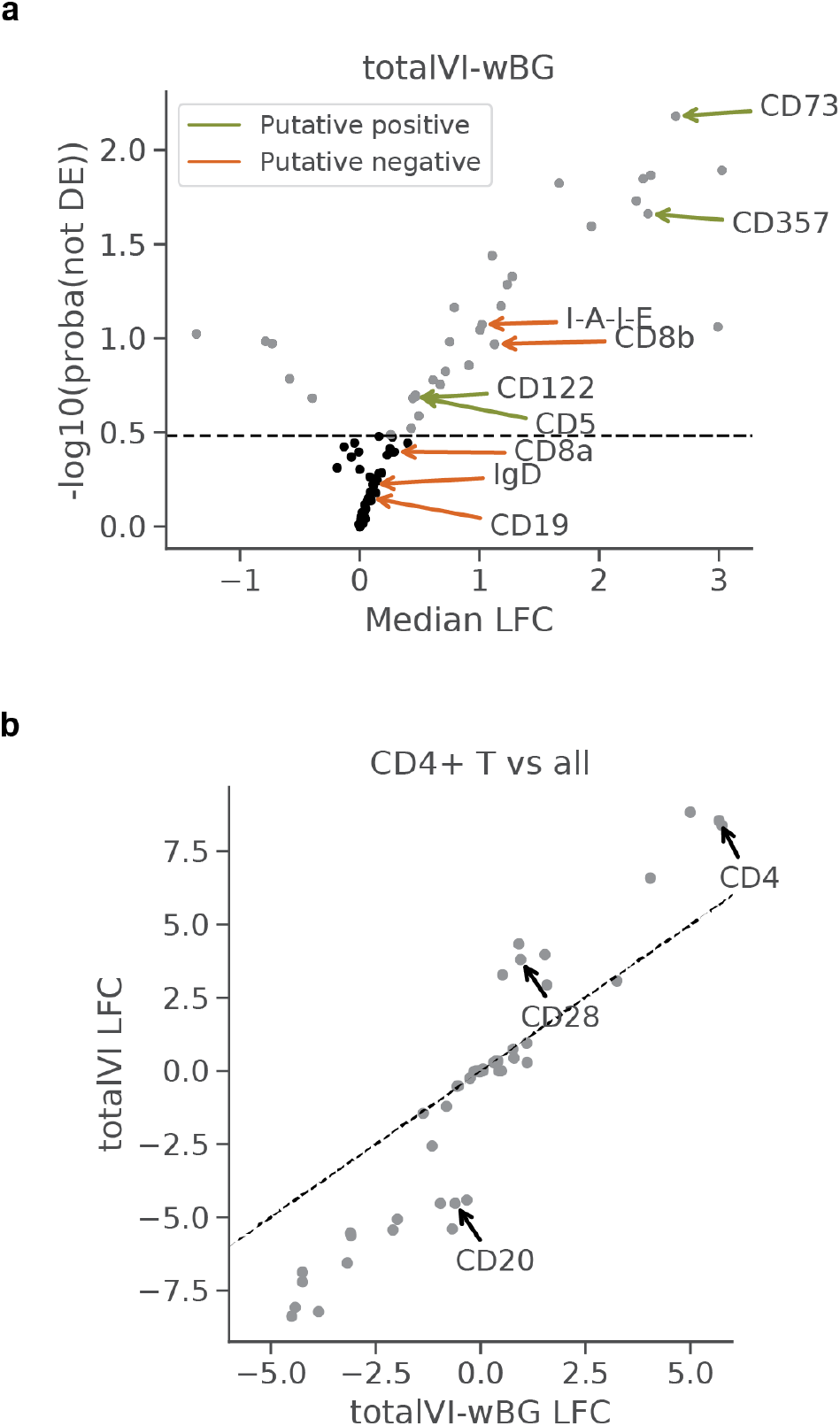
Differential expression using totalVI-wBG, which does not correct for the protein background component. **a** Volcano plot for the ICOS-high Tregs vs CD4 T cells test. The same putative positives and negatives as Figure 4d are highlighted here. **b** The LFC estimates for totalVI and totalVI-wBG on the CD4 T cells versus all others test.

**Figure S13:**
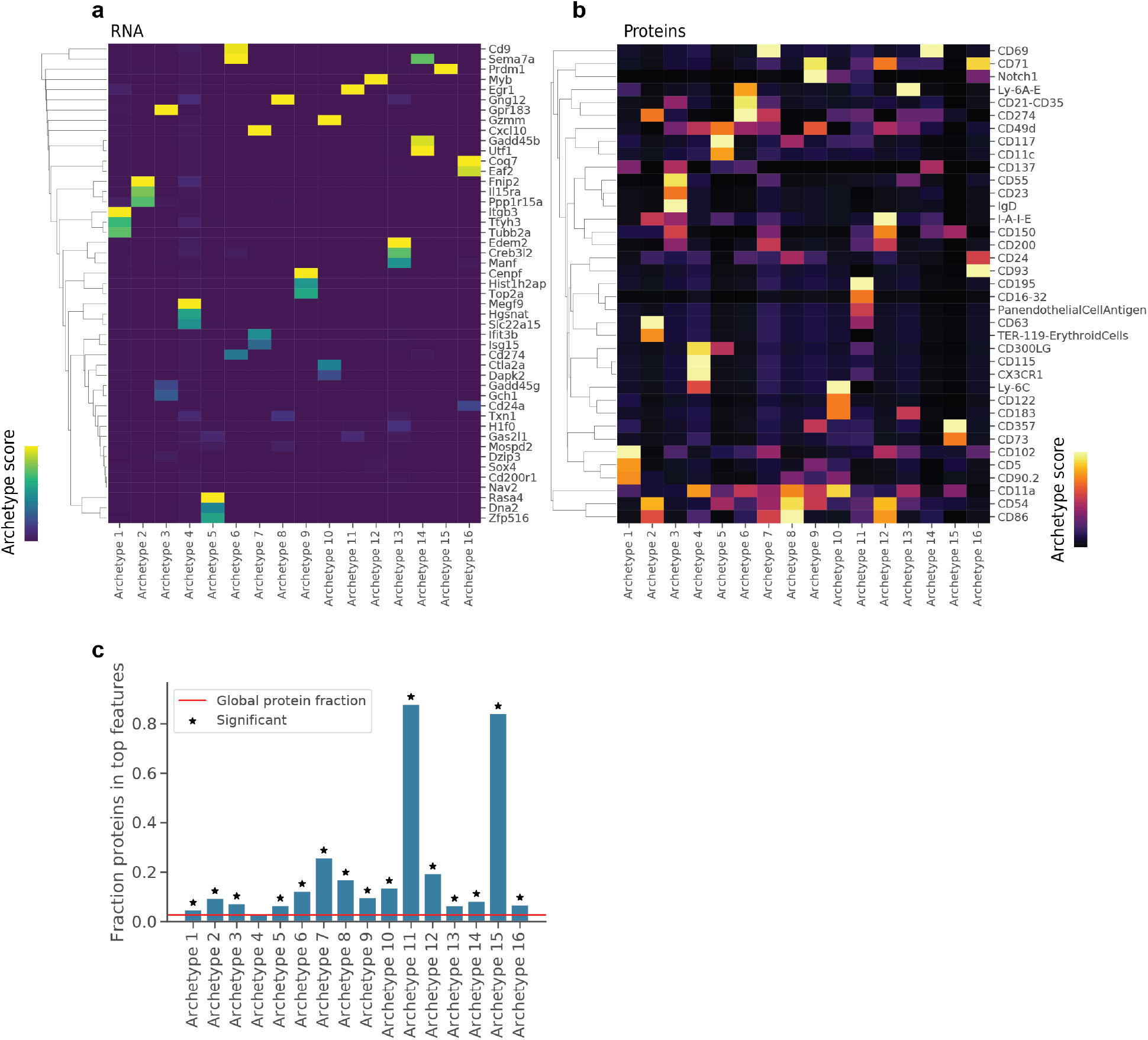
Interpreting totalVI latent dimensions with archetypal analysis. **a, b**, Heatmap of top **(a)** gene and **(b)** protein features for each archetype. The archetype score corresponds to the standard scaled archetypal expression profiles (Methods 4.13). Heatmaps are individually column normalized for visualization. **c**, Fraction of proteins in top archetypal features for each archetype. Features in each archetype were selected in the “top” if they had an archetype score of greater than 2. For these features, we performed a hypergeometric test to determine if proteins were over-represented in this feature set relative to the global distribution of feature types.

**Figure S14:**
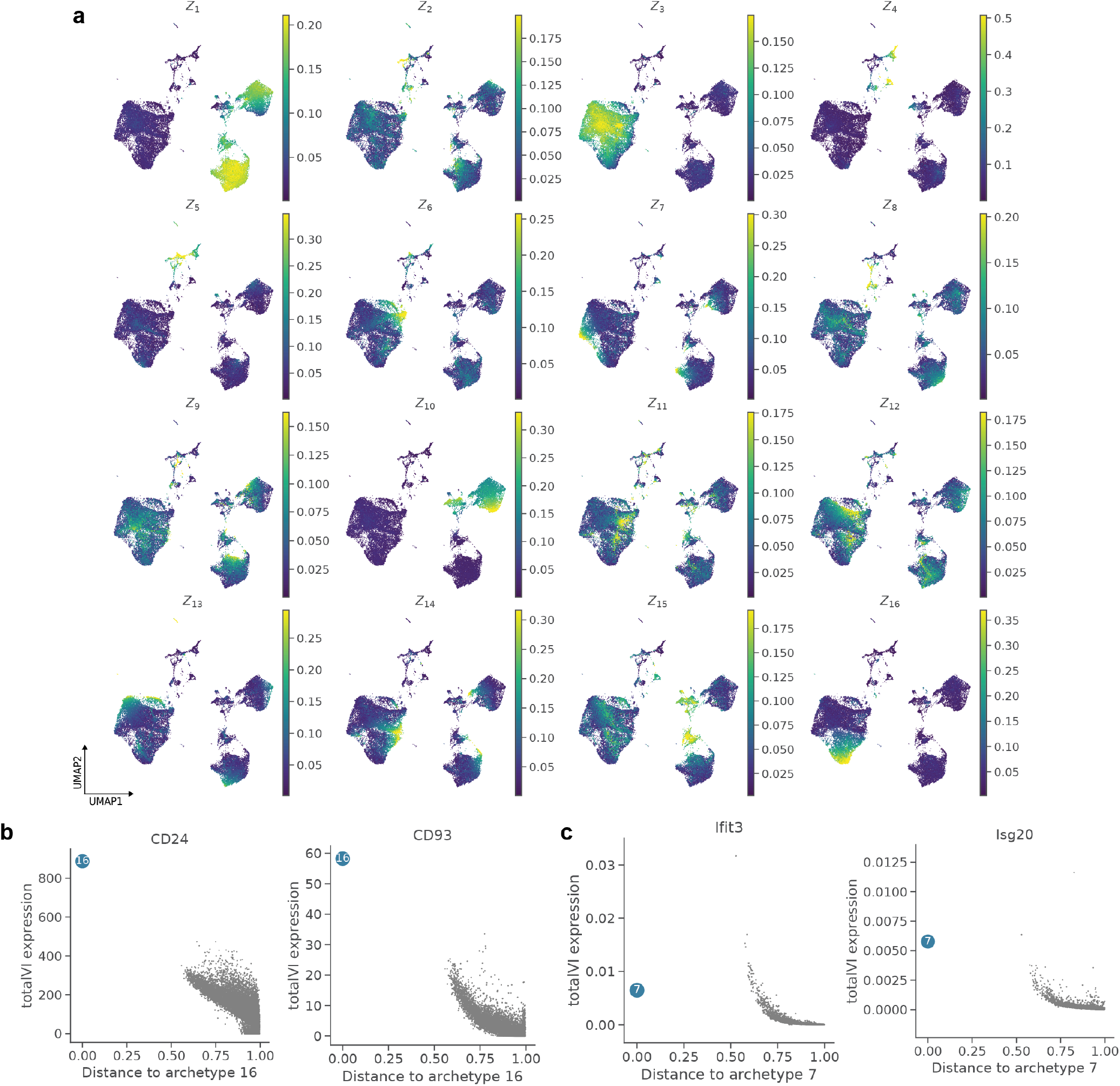
Visualization of archetypes in totalVI-intersect model of SLN-all. **a**, UMAP plots of SLN-all cells colored by latent dimension value. **b**, totalVI protein expression for CD24 and CD93 proteins as a function of distance to archetype 16. **c**, totalVI denoised expression for Isg20 and Ifit3 genes as a function of distance to archetype 7. Archetype is colored in blue, all other cells in grey.

**Figure S15:**
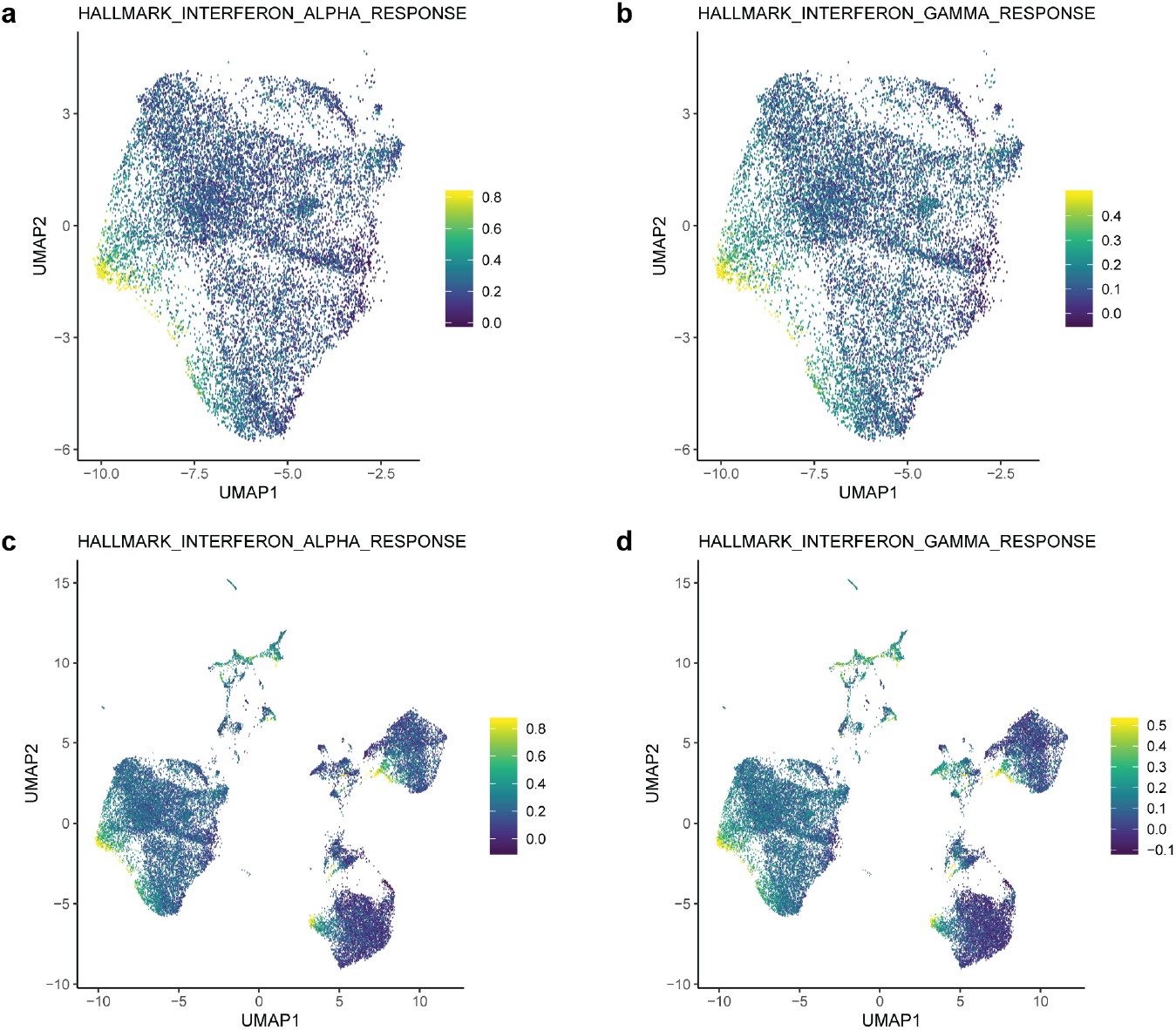
Interferon signatures in the mouse spleen and lymph node. **a, b**, UMAP of totalVI latent space for B cells of the SLN-all dataset **(a)** colored by the Hallmark Interferon Alpha Response signature score and **b** colored by the Hallmark Interferon Gamma Response signature score (Methods 4.14). **c, d**, Same as in (a, b), but for all cells in SLN-all.

**Figure S16:**
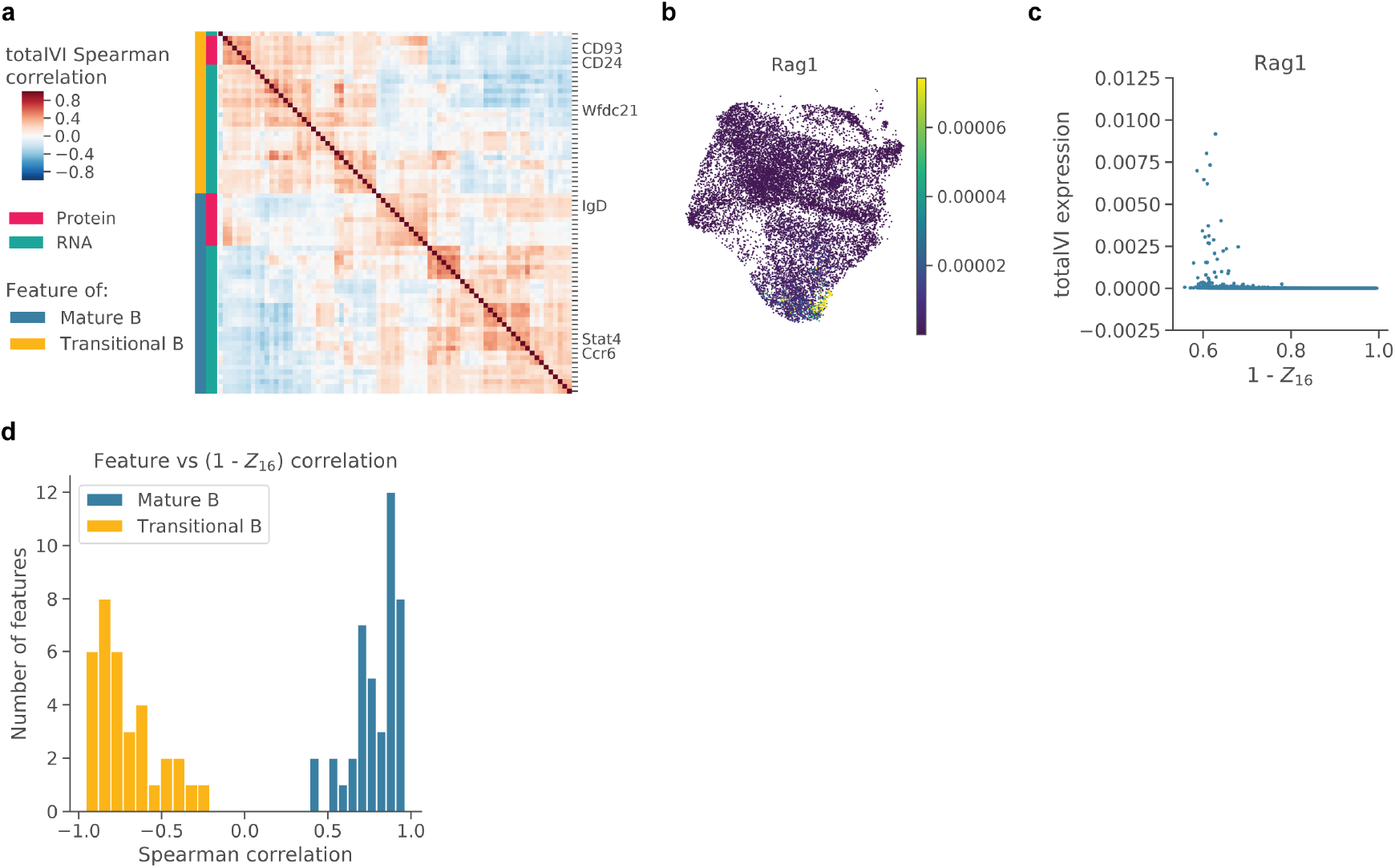
totalVI identifies correlated modules of RNA and proteins that are associated with the maturation of transitional B cells. **a**, totalVI Spearman correlations in mature B cells between the same RNA and proteins as in Figure 5h. Features were hierarchically clustered within mature B cells. **b**, UMAP of the totalVI latent space colored by totalVI RNA expression of Rag1. **c**, totalVI RNA expression of Rag1 as a function of 1 − *Z*_16_ (the totalVI latent dimension associated with transitional B cells). **d**, Histogram of Spearman correlations between each feature in (a) and 1 − *Z*_16_.

**Figure S17:**
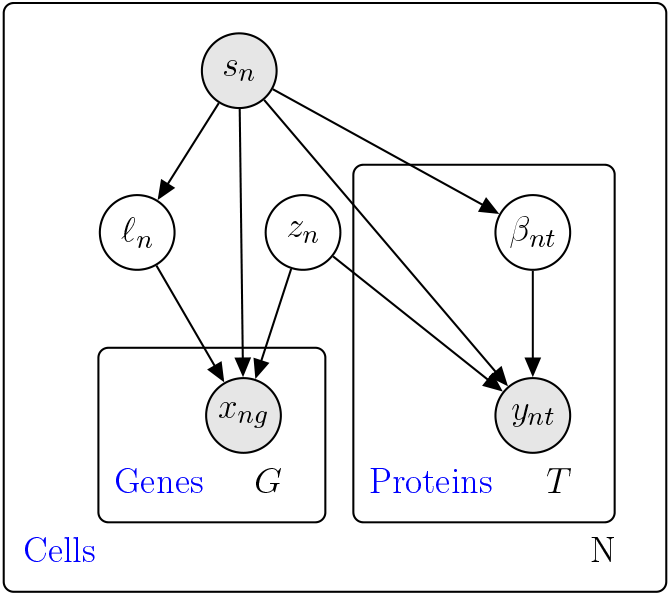
totalVI probabilistic graphical model. Shaded nodes represent observed random variables. Unshaded nodes represent latent variables. Edges denote conditional independence. Rectangles (“plates”) represent independent replication.

## A Protein considerations

### A.1 Experimental considerations

Sources of technical variation in CITE-seq experiments, particularly protein background, are dependent on the experimental method itself. There are a number of potential experimental sources of background. We primarily discussed ambient antibodies and non-specific antibody binding. Another potential source of background could arise from oligonucleotide barcodes that become dissociated from their conjugated antibody. Similarly to barcoded antibodies, ambient oligonucleotide barcodes could contaminate cell-containing droplets or could non-specifically bind to the cell surface. In this study, we do not distinguish between background due to antibodies or background due to free oligonucleotide barcodes.

Although in our experiments we used the standard CITE-seq protocol, there are a number of protocol modifications that could change the amount of background. For instance, increasing the number of washes after staining cells with antibodies could reduce ambient background. Alternatively, a buffer modification could reduce the amount of non-specific binding. Both washing and blocking are frequently considered in flow cytometry protocol designs. However, implementing these protocol changes in an effort to eliminate background could come with trade-offs; reducing background by washing and blocking would likely reduce true signal by reducing total cell numbers or blocking specific binding, respectively.

Another common experimental practice to modulate the amount of background is antibody titration, meaning that different antibodies are added to the experiment at different concentrations. At the optimal titration, an antibody would have the maximal signal-to-noise ratio. This would require the antibody to be present at a sufficient concentration to specifically bind its target protein and generate a detectable signal, but not at so high a concentration as to increase protein background by binding non-specifically or remaining at high concentration in the ambient solution. In a CITE-seq experiment, it is possible that the recommended antibody concentration is too low to detect foreground signal from a given protein. Finding the optimal concentration for each antibody might be challenging, since the optimal concentration might be different for different cell types or experimental systems. If titrations are modified per antibody, there are a few points to consider. When antibodies are titrated at different concentrations, it becomes infeasible to quantify absolute protein levels. For example, it would not be possible to determine if one protein was expressed at a higher level than another protein. In addition, even if every protein in the cell were measured with a theoretical unbiased antibody panel, the sum of all protein counts from a cell (referred to as the protein library size) could not serve as a meaningful estimate of total protein molecules in a cell or cell size because the relative amounts of each protein have been manipulated.

In our modeling and analysis, we considered whether protein library size should be taken into account. In scRNA-seq data, we consider library size to be a nuisance factor that is reflective of a combination of sequencing depth and cell size. In CITE-seq experiments where only selected markers are measured, there is no guarantee that the markers selected are representative of the total protein content in the cell. For example, a cell with few detected protein counts might in reality express other unmeasured proteins at higher levels, meaning this cell’s total counts reflect the selection of markers rather than reflecting nuisance variation like sequencing depth or cell size. Therefore, treating protein library size as a nuisance factor does not make sense in this context. Because measured proteins are biased in this manner, we do not assume that the protein data is compositional, as is assumed by other methods that use a centered log ratio transformation for normalization. In the future, more unbiased protein panels might necessitate further consideration of protein library size in CITE-seq data analysis.

For proteins that are expressed at low levels or are only expressed in rare cell types, it might be necessary to increase sequencing depth to increase the sensitivity to detect these molecules. In addition, sequencing depth could play a role in the ability of totalVI to decouple protein foreground and background (i.e., with more counts, it might become easier to separate what appear to be overlapping foreground and background distributions). Determining the optimal sequencing depth for protein panels could be an important cost consideration in CITE-seq experiments, particularly as the size of protein panels increases. Since in the CITE-seq protocol RNA and protein libraries are prepared independently, future work could determine the value of these two molecules in various downstream analysis tasks to make recommendations for sequencing depth for each library.

Because the barcoded antibodies used in this study came from clones that have been previously validated, we were surprised to find that some common protein markers (e.g., IgM, CCR7) appeared to have little or no signal. Aside from the consideration of titration and sequencing depth discussed above, an additional explanation could be the uniform staining conditions for all antibodies in the CITE-seq panel simultaneously. For example, the chemokine receptor CCR7 is a well-documented marker in T cells and typically requires staining at higher temperature and for longer times than other antibodies due to its constant cycling onto and off of the cell surface. For future CITE-seq experiments, it might be worthwhile to consider the optimal staining conditions (e.g., time and temperature) for each antibody independently rather than staining with all antibodies at once.

### A.2 Modeling considerations

Guided by the points raised in Appendix A.1, we considered a variety of protein likelihoods before settling on the version used in this manuscript. Among our considerations were the interpretability of the parameters as well as how well the likelihood captured our view on the CITE-seq protein data generating process. For example, we considered models that included a latent variable for the protein library size (analogous to *ℓ_n_* for RNA), though as discussed in Appendix A.1, protein library size and RNA library size do not convey the same information.

We also considered alternative models to decouple protein background. Initially, we attempted to use models that assume every cell receives the same distribution of protein background scaled by some cell-specific scalar. However, we found these models inadequate for decoupling the protein signal, which again suggests that ambient antibodies can not fully explain the protein background.

The likelihood we used in this manuscript,

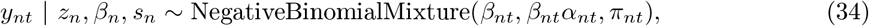

also has some import downstream considerations. First, the mixture assumes that the observed counts for a given cell *n* and protein *t* are generated from either the background component (with probability *π_nt_*) or the foreground component (with probability 1 − *π_nt_*). Despite the fact that the background mean parameter *β_nt_* appears in the foreground mean *β_ntαnt_*, this likelihood does not allow us to correct the foreground for possible background contamination. Here, the double usage of *β_nt_* is to help identify the mixture model. In other words, we cannot “subtract the background” from *y_nt_* that are determined to be in the foreground. Perhaps this limitation could be addressed in future work, in which different latent variables are associated with local and global sources of background; though this will require greater understanding of the experimental mechanisms previously discussed.

## B Integrating out latent variables

Here we show that if

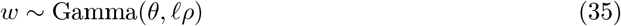

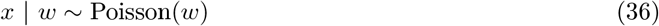

then *x* ∼ NegativeBinomial(*ℓ_ρ_, θ*). Note that we have dropped all subscripts and each variable here is a scalar. While we parameterize the Gamma with its shape and mean, a more conventional form is with its shape and rate, so *w* ∼ Gamma(*θ, θ/ℓ_ρ_*)

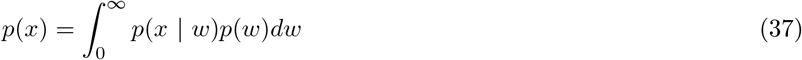

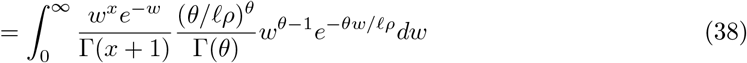

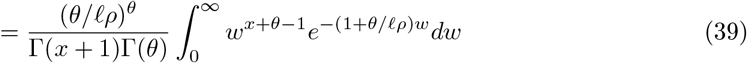

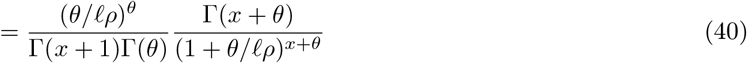

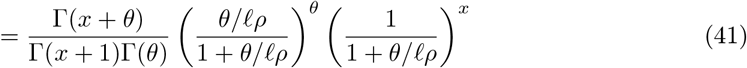

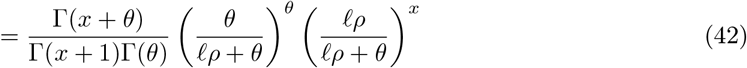

In the fourth line, we use the fact that the integrand is an unnormalized gamma distribution. The final line is a negative binomial distribution with mean *ℓ_ρ_* and inverse dispersion *θ*. Therefore, we have a direct link between the parameters of the negative binomial and the underlying parameters of the Poisson rate. Finally, we note that we could have written *w* ∼ Gamma(*θ, ρ*) and *x* | *w* ∼ Poisson(*ℓ_w_*) and achieved the same result.

## C totalVI implementation details

### Evidence lower bound derivation

Here we derive the Evidence lower bound (ELBO), which is ultimately used in optimizing the model and variational parameters. For shorthand, we drop subscripts and inference and generative parameters *ν* and *η*. The joint likelihood based on the totalVI generative model for a single cell factorizes as

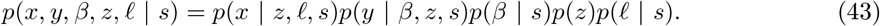

In the model specification, we use the latent variable *z* ∼ LogisticNormal(0, *I*). Here we use the logistic normal definition of [119], in which a normal random variable *δ* ∼ Normal(0, *I*) is transformed by a softmax function, embedding the random variable in the simplex. Thus, *z* = softmax(*δ*). However, the softmax function is not invertible, so for simplicity we consider the underlying latent variable *δ*. In this setting, *z*, which is ultimately the input to the decoder, is treated as a likelihood parameter. Therefore, we can rewrite the joint likelihood as

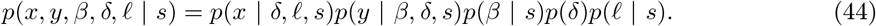

To perform variational inference, we define the variational posterior distribution as

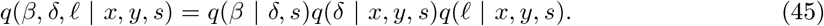

The Evidence LOwer Bound (ELBO) is derived using Jensen’s inequality. We use the shorthand notation *q*(*β*, *δ*, *l*) = *q*(*β*, *δ*, *l* | *x, y, s*).

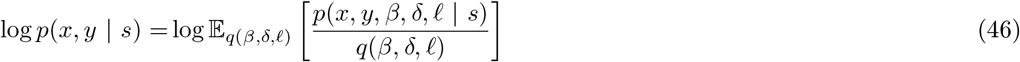

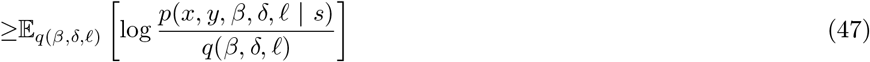

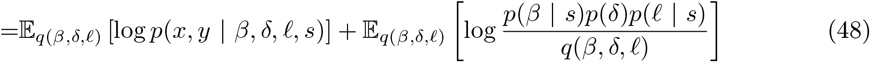

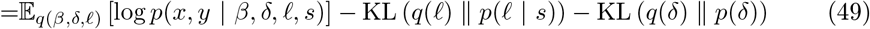

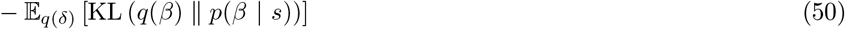

To compute the KL divergences of lognormal random variables we note that the KL divergence is invariant to invertible transformations, so the KL can be computed in closed form using the KL between normal random variables. The log likelihood of a negative binomial mixture distribution is computed using numerically stable functions in Pytorch (Appendix D). The ELBO derived here is amenable to the reparameterization trick used to train VAEs [25]. Estimates of the expectations in the ELBO are taken via Monte Carlo and noisy gradients of the ELBO are used in a stochastic optimization scheme. A sketch of the inference procedure for totalVI is in Algorithm 2.

#### Algorithm 2

Inference for totalVI

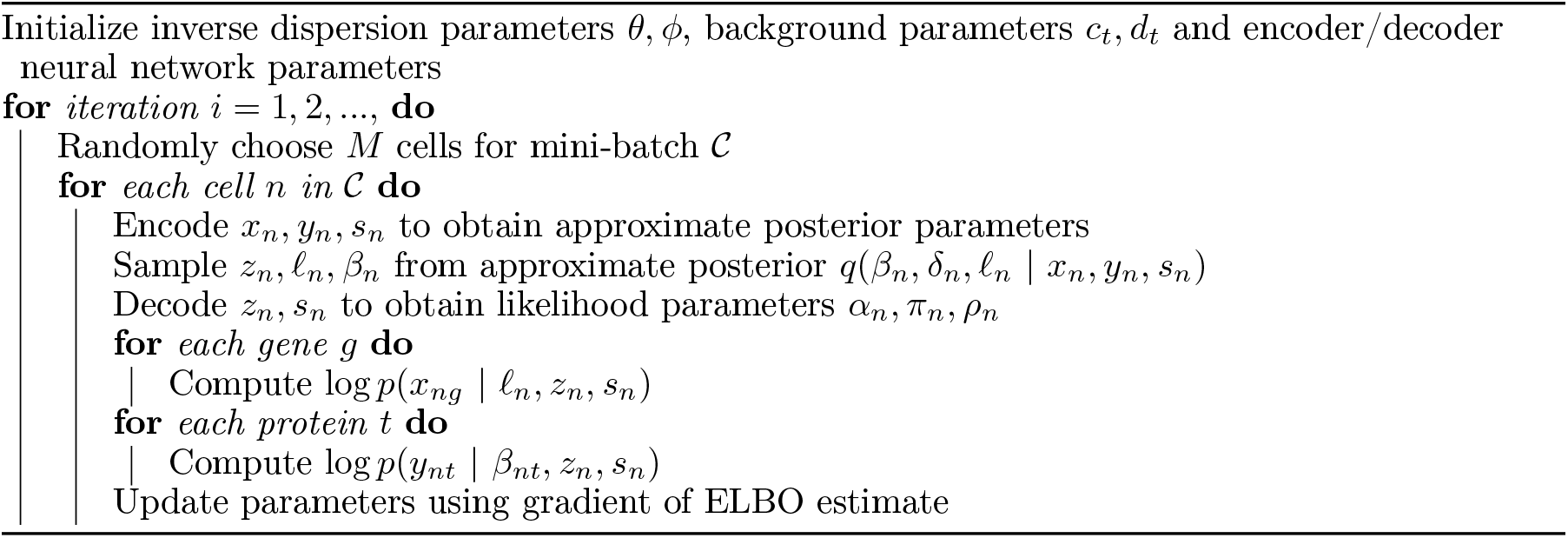

### Approximate posterior specification

The approximate posterior distributions are specified by neural networks. In particular, one neural network takes as input the triple (*x, y, s*) and outputs the parameters of *q*(*δ* | *x, y, s*)*q*(*ℓ* | *x, y, s*). An additional neural network maps (*δ, s*) to the mean and variance parameters of *q*(*β z, s*) through *z*. The variational distributions match their priors in family (e.g., *q*(*δ* | *x, y, s*) is a Gaussian with diagonal covariance matrix). A posterior draw of *z*, which we used as input to clustering and visualization algorithms, as well as used for archetypal analysis is then obtained by

1. Draw *δ* from *q*(*δ* | *x, y, s*)
2. Set *z* = softmax(*δ*)

### Neural networks

The encoder neural network has one shared hidden layer of 256 nodes followed by a layer of 512 nodes. The output of the 512 nodes are split in half and are used as input for linear layers that parameterize *q*(*δ* | *x, y, s*) and *q*(*ℓ* | *x, y, s*), respectively. The final encoder neural network of one hidden layer and 256 hidden nodes takes as input (*z, s*) and outputs the parameters of *q*(*β* | *δ, s*). The parameters of *δ* are 20-dimensional mean and variance parameters. The decoder consists of three individual neural networks each with one hidden layer and 256 nodes. The first maps to the parameters of the mean of the RNA likelihood (*ρn*). The second maps to the foreground mean of the protein likelihood (*α_n_*). The third maps to the mixing parameter of the protein likelihood mixture (*π_n_*). Each of these decoder networks takes as input (*z, s*). Furthermore, (*z, s*) are reinjected at each hidden layer. All neural networks use batch normalization [120], dropout regularization [121], and ReLU activations in hidden layers. The model parameters *θ*, *ϕ*, *c*, and *d* are treated as global neural network parameters, optimized to maximize the ELBO.

### Hyperparameters

The neural network architecture previously described was used throughout this manuscript without modification. There are a number of other hyperparameters used to train neural networks that we also held constant in all experiments. This includes the learning rate of the optimizer (`lr=4e-3`), the size of the training test (90%), the KL warmup scheme (`0.75 × NumCells` minibatches, or 213 epochs with the fixed training set size of 90%), and the number of training epochs (500 epochs). An early stopping scheme is performed with respect to the 10% of cells in the test set. If there is no improvement of the held-out ELBO of the test set with 30 epochs, the learning rate is multiplied by 0.6. If there is no improvement after 45 epochs, the inference procedure is stopped. The totalVI package includes a module for likelihood-guided hyperparameter tuning, which can be used to revisit the default parameters, especially as the experimental protocol evolves.

## D Numerical considerations for the negative binomial mixture distribution

For a single negative binomial mixture component, we use numerically stable functions provided by PyTorch (e.g., the log gamma function). For a mixture of negative binomials, we rewrite the distribution to use numerically stable functions like `logsumexp` and `softplus`.

Let *p^b^*(*y*) = *p*(*y* | *z, μ, s, v* = 1) be the probability mass function for the background and *p^f^* (*y*) = *p*(*y* | *z, μ, s, v* = 0) be the probability mass function for the foreground. Then by integrating over *v* (recalling that *v* ∼ Bernoulli(*π*)),

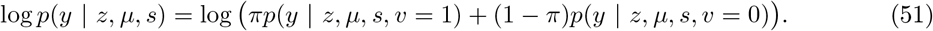

We now rewrite this in a form more amenable for optimization. We recall from Algorithm 1 that *π* = *h_π_*(*z, s*; Ω). Thus, with *c*(*z, s*) = logit(*h_π_*(*z, s*)), then *π* = 1/(1 + exp(−*c*(*z, s*))). Also, let 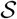 be the softplus function: *x* → log(1 + *e^x^*).

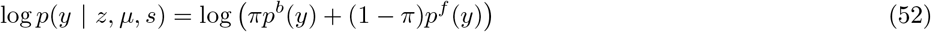

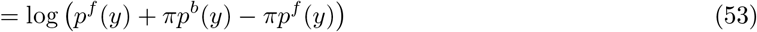

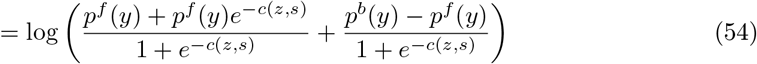

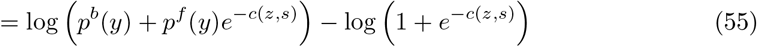

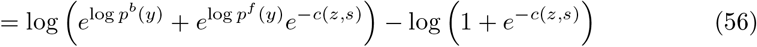

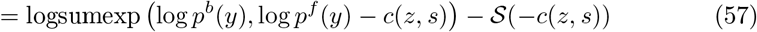

